# Functional projection of cognitive functions through the human corpus callosum

**DOI:** 10.64898/2026.06.16.732587

**Authors:** Rolando Bonandrini, Marco Tettamanti, Claudio Luzzatti

## Abstract

Reconciling the anatomical observation that the human brain comprises two asymmetrical halves with the phenomenal unity of the mind is a puzzle that has challenged neuroscientists since the dawn of research in the field. White-matter commissural fibres of the corpus callosum constitute a critical anatomical substrate for the functional resolution of this anatomical duality. However, the extent of the functional involvement of the callosum in different domains of cognition represents, to this day, a mostly uncharted territory. Here we present a probabilistic characterization of callosal involvement in a set of cognitive functions. In particular, we estimated structural callosal connections by means of the Disconnectome approach applied to a reference sample of healthy participants while using the macro-anatomical cortical areas contained in the Harvard-Oxford template as seeds. By multiplying structural connectivity by the involvement of each cortical area in a set of cognitive functions (as derived from Neurosynth meta-analyses), we produced a voxel-wise characterization of the corpus callosum in different functional domains. We were able to highlight greater involvement of posterior callosal regions in vision and episodic memory, greater involvement of more anterior callosal regions in decision making and working memory, with somatosensory and motor functions more related to the central dorsal portion of the callosum.

## Introduction

The involvement of white matter in cognition is a long-standing issue in human neuroscience. Functional neuroimaging has indeed produced substantial literature mapping cognitive functions to grey matter (Dolan, 2008; Friston, 2009; Frith & Friston, 2013; Perani, 2008; Price, 2012). However, despite structural techniques have allowed reconstructions of white matter bundles (Catani, 2007; Le Bihan et al., 2001; Maier-Hein et al., 2017; Tournier et al., 2011) and despite the Aristotelian principle whereby structure and function intuitively “go together” (Blits, 1999; Catani, 2007), there is still limited data on the functional characterization of white matter (Catani & Thiebaut de Schotten, 2012).

In this context, the corpus callosum (“tough body” in Latin) is of particular interest: it contains 200–300 million axons and it constitutes the largest commissural tract in the human brain (Aboitiz et al., 1992; Catani et al., 2012; Tomasch, 1954). From an anatomical point of view (e.g., Shah et al., 2021), it is subdivided in the rostrum (i.e., “beak”: the most ventral part), the genu (i.e., “knee”: the most anterior portion, which bends into the rostrum), the body (i.e., the intermediate part), the isthmus (i.e., narrow part that connects the body with the most posterior portion), and the splenium (i.e., “band”: the most posterior part, that looks rounded in a medial sagittal section of the brain)^1^. Its connections follow a rostro-caudal (i.e., antero-posterior) gradient (Bernhardt et al., 2022)^2^: more anterior portions of the callosum connect more anterior cortical areas and vice-versa (Barakovic et al., 2021; Friedrich et al., 2020; Hofer & Frahm, 2006; Barbaresi et al., 2024). In addition, a dorso-ventral gradient in callosal connections has been recently described (Xiong et al., 2024).

Although traditionally thought to serve a purely structural purpose (i.e., preventing the two cerebral hemispheres from collapsing onto one another; Mooshagian, 2008), neuroscience inquiry in the second half of the 20^th^ century has brought the callosum under the spotlight of cognitive research (e.g., Sperry, 1964). Indeed, empirical evidence highlighted its crucial role in giving rise to the phenomena of inter-hemispheric communication, competition and integration (for a review, see Bloom & Hynd, 2005; van der Knaap & van der Ham, 2011; see also Hinkley et al., 2012; Santander et al., 2025; Bekir et al., 2025) which crucially contribute to resolving the anatomical duality of the brain into the phenomenal unity of the mind (Gazzaniga, 2000). Still, in spite of the extensive wealth of studies on split-brain patients that highlights the role of the corpus callosum in cognition as a whole (e.g., Berlucchi et al., 1997; Gazzaniga, 1995; Gazzaniga, 2005; LeDoux & Gazzaniga, 1981; Levy & Trevarthen, 1977; Metcalfe et al., 1995; Zaidel et al., 1999), fine-grained evidence on the functional role of each portion of the callosum is still limited. Indeed, data from neuropsychological patients (extensively reviewed in Catani et al., 2012, see also Bozzali et al., 2012; Caleo, 2018; Li et al., 2015; Rode et al., 2010) point towards the involvement of the body of the callosum in somatosensory and motor functions, as well as the involvement of the splenium in vision. However, these inferences are confined to a relatively limited collection of cases, which hampers population-wise inference. In addition, the spatial resolution of such anatomo-clinical associations (especially in less recent studies) rarely exceeds that of macroanatomical callosal areas (i.e., rostrum vs. genu, etc).

An alternative framework has attempted a functional characterization of the corpus callosum using fMRI. Indeed, Blood Oxygenation Level Dependent (BOLD) response was observed in the corpus callosum, although studies were limited to cognitive tasks explicitly requiring inter-hemispheric transfer (D’Arcy et al., 2006; Gawryluk et al., 2011; Gawryluk et al., 2014; Tettamanti et al., 2002). A notable exception in this regard is the study by Fabri et al. (2011), in which anterior, central, central/posterior and splenial portions of the callosum were activated by taste stimuli, motor tasks, tactile stimuli and visual stimuli, respectively. Still, the extent of these associations does not go beyond a handful of volume units in the brain (voxels), which is likely due to the fact that blood volume and flow (on which the BOLD signal relies) are three to seven times lower in white matter than in grey matter (Gawryluk et al., 2014; Helenius et al., 2003; Preibisch & Haase, 2001; Rostrup et al., 2000). This limitation, combined with those derived from anatomo-clinical associations, has produced a rather piecemeal understanding of the involvement of the callosum in cognitive functions.

It is worth mentioning that recent studies have combined structural and functional resting-state MRI and identified white-matter functional networks towards which structural and functional connectivity from the corpus callosum align (Peer et al., 2017; Wang et al, 2020; Wang et al., 2021). Diffusion MRI data has also helped unravel the involvement of the corpus callosum in individually targeted domains of cognition, with notable examples involving emotion recognition (Orlando et al., 2023), motor performance (Hannanu et al., 2022), sensorimotor synchronization (Blecher, Tal & Ben-Shachar, 2016) attention (Chechlacz et al., 2015; Mesaros et al., 2009), vocabulary and matrix reasoning (Danielsen et al., 2020).

However, the available literature has not yet produced a comprehensive atlas of the functional anatomy of the corpus callosum across specific cognitive functions. Diffusion MRI studies indeed investigated individual cognitive domains in isolation, inferring callosal involvement from associations between white-matter properties and behavioural performance (i.e., “where in the callosum does fractional anisotropy correlate with this particular task?”). While this approach is well suited to identifying structure-function relationships within a given domain, it does not assure that results from a single study can be reliably generalized to the whole cognitive domain they pertain to, nor it provides a common reference framework in which multiple cognitive functions can be directly compared. On the other hand, studies that focused on gradient- or functional-connectivity-based organization of the corpus callosum (Peer et al., 2017; Wang et al, 2020; Wang et al., 2021; Friedrich et al., 2020) describe large-scale principles of structural or functional organization, but they do not associate specific cognitive operations with individual callosal voxels. Here we sought to directly address the insofar unanswered question of what type of computation does every voxel of the corpus callosum support during cognitive tasks, while adopting an approach that assures comparability across domains and meta-analytical convergence within each cognitive domain (e.g., “Which portions of the callosum support the cognitive operations underlying—for instance—‘working memory’, defined through convergence between dozens of experiments?”).

One promising methodological standpoint from which to address this issue derives from the so-called *Functionnecome*. The *Functionnecome* approach proposes to combine task-based fMRI and tractographic data to project function onto structure (by using voxel-wise structural priors), with the goal to infer brain functioning at the network level (Nozais et al., 2021; Nozais et al., 2023). Remarkably, this approach allows, although indirectly (i.e., white matter signal is projected from grey matter, not measured directly), to characterize white matter tracts at a functional level. In the present work, we sought to address the current limitations of the literature by adopting a methodology of structure-function projection inspired by the *Functionnectome,* to conduct a large-scale functional characterization of the corpus callosum based on functional data from twelve cognitive domains. Critically, with respect to the original formulation of the *Functionnectome*, we (1) adopted an atlas-based approach instead of voxel-wise structural priors to reduce the computational workload, (2) adopted meta-analytical data instead of first-hand fMRI data to minimize the impact of study-specific methodological choices, and (3) introduced a validation procedure back-projecting function from structure to minimize the chances that results may simply be statistical artefacts.

The overarching goal of this work is to assign voxel-level cognitive significance to the anatomical organization of the corpus callosum, by constructing a meta-analytically informed probabilistic atlas of its functional topography across multiple cognitive domains.

## Materials and Methods

Here we present a novel pipeline (inspired by the *Functionnectome*), which builds on the combination of the Disconnectome and Neurosynth approaches. Intermediate steps were validated through the estimation of structural gradients in structural connections. Functional projections of cognitive functions on the corpus callosum were validated through back projection from structure to function. With the only exception of Disconnectome analyses (which were conducted using standard code provided by developers of the toolbox), all other analytical steps detailed below were performed through MATLAB (R2024a) – and in particular SPM 12 (Ashburner et al., 2014) - and R (version 4.3.3) custom-made code. In what follows, the analytical steps (see Figure 1) are discussed in detail.

**Figure 1.**
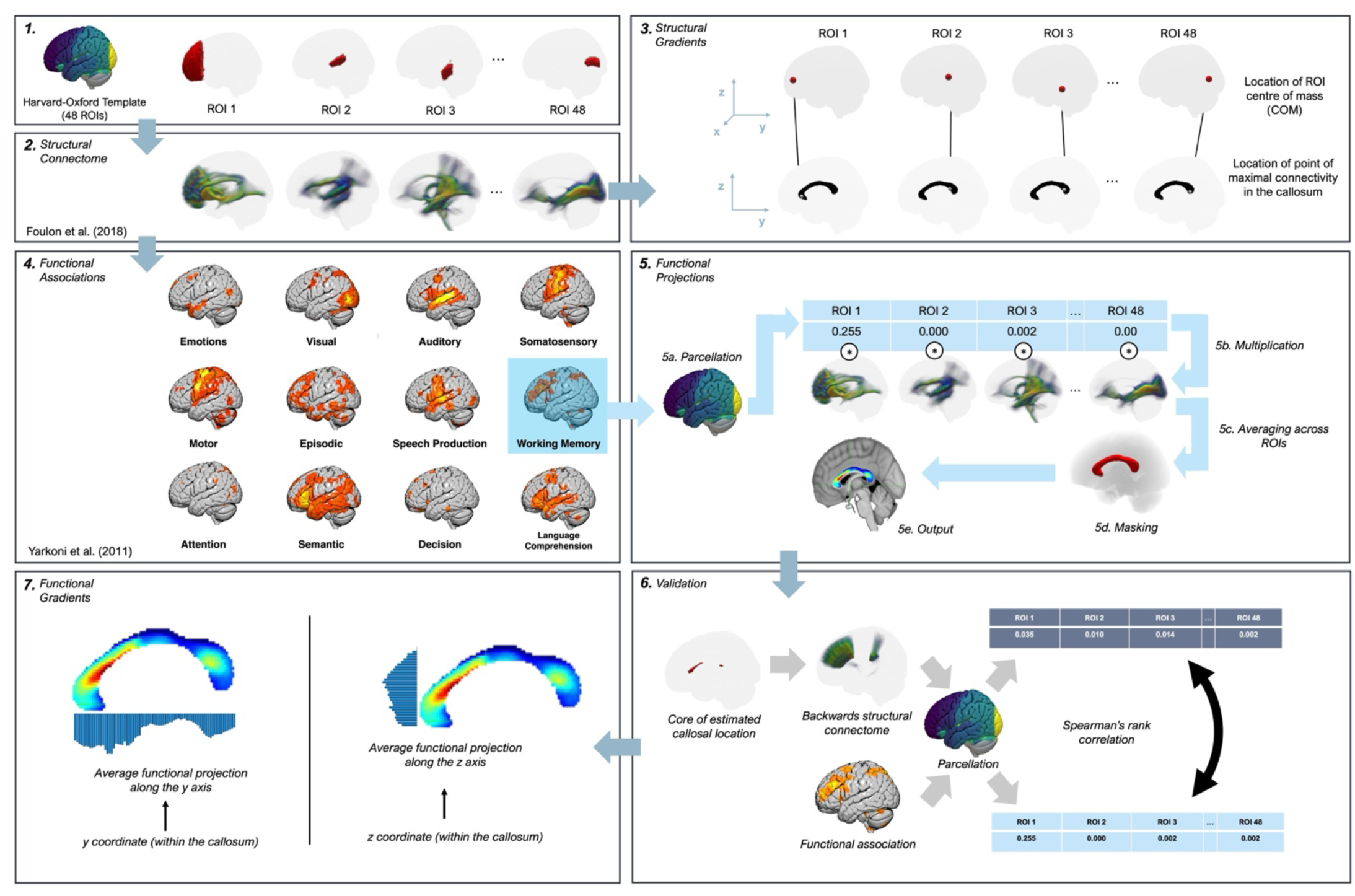
Overview of the analytical pipeline. Please note that ROI order has been changed for illustration purposes. Panel 1 shows examples of the cortical ROIs from the Harvard Oxford atlas used as input for connectome estimation. Panel 2 shows the output of ROI-based connectome analyses. Panel 3 shows the logic behind the estimation of structural gradients (i.e., relating ROI centre of mass and the locus of maximal connectivity in the callosum). Panel 4 shows Neurosynth meta-analytical maps for the functional domains taken into account. Panel 5 highlights the steps underlying callosal projection of cognitive functions: parcellation of functional maps, by-ROI multiplication by parcellated structural connections, averaging across ROIs, and masking. Panel 6 indicates the steps underlying validation: identification of the core of the functional projection, estimation of backward structural connections, parcellation and correlation with parcellated functional associations from Neurosynth. Panel 7 shows the estimation of functional gradients, relating (in separate analyses) y and z coordinates within the corpus callosum and average functional projections.

### Estimation of structural connections

We estimated the structural connections (Figure 1.1; 1.2) passing through the corpus callosum by means of the “disconnectome” tool implemented in the BCBToolkit software (Foulon et al., 2018), as recently applied to template-based structural (dis)connection mapping (Veronelli et al., 2024). In particular, we used the 96 (48 per hemisphere) regions of interest (ROIs) from the cortical^3^ Harvard-Oxford template (HarvardOxford-cort-maxprob-thr0-1mm; Desikan et al., 2006; Frazier et al., 2005; Goldstein et al., 2007; Makris et al., 2006) as seeds (Figure 1.1). The disconnectome approach uses diffusion-weighted magnetic resonance images of healthy participants (internally coded in the distributed BCBToolkit) to track the fibers passing through the location of a region of interest, which is typically (although not necessarily) a brain lesion. The method estimates the probability - from 0 to 1 - of each voxel of being connected from the target site (or disconnected, if the target site is a lesion; Thiebaut de Schotten et al., 2015). To do so, each target location is registered to each participant’s native space and used as a seed for tractography (Thiebaut de Schotten et al., 2011) in Trackvis (Wang et al., 2007). Each tractography from each control subject is then converted into a visitation map, binarized and back-transformed to the Montreal Neurological Institute (MNI) space. As the final output of the “disconnectome” tool, a percentage overlap map is produced by summing, for each voxel of the MNI space, the normalized visitation maps of the reference subjects (Figure 1.2). Our analysis was conducted on fully processed tractographic data contained in the freely available Disconnectome Package X (https://storage.googleapis.com/bcblabweb/open_data.html; Salvalaggio et al., 2020) including 178 healthy participants (109 female, 69 male; 21 participants aged 22-25; 85 participants aged 26-30; 70 participants aged 31-35; 2 participants aged 36+), in turn derived from the Human Connectome Project (HCP7T, Van Essen et al., 2012; for details on acquisition, see Vu et al., 2015). We did not collect nor pre-process data first-hand. For further information on data acquisition and pre-processing, please refer to the original articles from which the dataset for our (dis)connectome analyses was derived. The (dis)connectome analysis was conducted without thresholding (Threshold= 0.0) to capture the full range of probability of structural connection. Structural connectivity maps were then averaged for each couple of homologous contralateral ROIs and masked using the JHU callosal template to exclude other anatomical structures (Hua et al., 2008; Mori et al., 2005; Wakana et al., 2007; see Supplementary Material: Medial Sagittal Callosal Atlas).

### Estimation of structural gradients

In this analysis we sought to obtain a comprehensive account of the direction of structural connections travelling through the corpus callosum. This was performed as a preliminary analytical step before estimating the involvement of the corpus callosum in cognitive functions. In particular, we explored the association between the location of a pair of homologous contralateral cortical areas (e.g., the left and right middle occipital gyri) and the location of the white matter callosal bundles resulting from them. More specifically, the location of cortical areas was operationalized as the centre of mass (COM, Fesl et al., 2008)^4^ of each area (i.e., a ROI in the Harvard-Oxford template), whereas the location of the white matter callosal bundle was operationalized as the location in the medial sagittal slice of the callosum of the point of highest probability of structural connection from the target cortical region (after averaging the (dis)connectome map from the left and right ROI). We sought to model the association between the location of cortical regions (along the rostro-caudal, dorso-ventral and medial-lateral axes) with the location (along the rostro-caudal and dorso-ventral axes) of the point of maximal connectivity^5^ in the medial slice of the callosum (see Xiong et al., 2024 for a similar approach) (Figure 1.3). To do so, we ran two multiple linear regressions: one with the medial-lateral, rostro-caudal, and dorso-ventral location of the cortical ROI as independent variables and the rostro-caudal location of the point of maximal structural connectivity as dependent variable; another one with the medial-lateral, rostro-caudal, and dorso-ventral location of the cortical ROI as independent variables and the dorso-ventral location of the point of maximal structural connectivity as dependent variable. In both models, both linear and quadratic associations between the independent and the dependent variables were modelled.

In the context of this analysis, the linear association between the rostro-caudal location of a given ROI and the rostro-caudal location of the point of resulting maximal structural connectivity in the callosum served as a replication of the rostro-caudal gradient in callosal connections ubiquitously observed in the literature. The same analysis in the dorso-ventral direction represents a replication of the recent findings by Xiong et al., 2024.

### Estimation of functional projections

We produced twelve whole-brain meta-analytic maps with the following keywords (Figure 1.4): “Emotions” (444 studies), “Visual” (3110 studies), “Auditory” (1252 studies), “Somatosensory” (674 studies), “Motor” (2565 studies), “Episodic memory” (332 studies), “Working memory” (1091 studies), “Semantic” (1031 studies), “Decision making” (509 studies), “Language comprehension” (107 studies), “Speech production” (107 studies), “Attention” (1831 studies) using Neurosynth (Yarkoni et al., 2011), and in particular through the Neurosynth functional association test, which indicates whether activation in a region occurs more consistently for studies that mention the current term than for studies that do not. Keywords were selected based on the work by Karolis et al. (2019), which highlighted that functional brain lateralization is organized around four functional axes, namely symbolic communication, perception/action, emotion, and decision-making. Our 12 keywords were selected to include these functional axes and to complement them with a more in-depth specification of perception and communication, as well as with memory and attentional domains. After thresholding meta-analytic maps to discard negative associations, they were then parcellated using the cortical Harvard-Oxford template (2mm resolution, to match the space of the meta-analytic map) to produce, for each ROI in the template, the average association with each of the twelve keywords across all voxels included in each given ROI. We then computed the functional projection on the corpus callosum of each of the twelve keywords: for each keyword, we multiplied (Figure 1.5.b) the medial sagittal structural (dis)connectome from each Harvard-Oxford ROI with the Neurosynth-based average functional association of that specific ROI with that specific keyword. For each keyword, this was iterated over each ROI, and the results were eventually averaged across ROI (Figure 1.5.c), to produce – upon masking (Figure 1.5.d) - a voxel-wise map of functional projection of each cognitive function in the medial sagittal slice of the corpus callosum (Figure 1.5.e).

### Validation analysis of functional projections

Given the explorative nature of our estimates of functional projections of cognitive functions on the corpus callosum, we decided to put a validation procedure in place. This was done to minimize the chance for us to overestimate structure-function associations in the corpus callosum. We therefore decided to embrace a conservative standpoint in this regard. The assumption behind our validation procedure is that we consider as reliable only the functional projections of cognitive functions in the corpus callosum that show a bidirectional association between structure and function: the portions of the corpus callosum that receive white matter projections from a given set of areas (associated to a given function) should also project back to the same areas in a similar way. In this context, given the functional projection of a cognitive function on the corpus callosum, the distribution of the involvement of grey matter regions in the back-projections from these callosal areas to the cortex (i.e., structural connectome from the putative callosal target to the cortex, parcellated using the cortical HarvardOxford atlas) should be correlated with the distribution of the functional involvement (Neurosynth meta analytical maps parcellated with the same atlas) of grey matter areas in that cognitive function. Therefore (Figure 1.6), in order to validate the functional projections, we generated a binary map for each functional domain (i.e., keyword) containing (in the medial sagittal section of the callosum) 1 in voxels whose estimated functional projection was greater than the midpoint of the range (in parallel with the default 0.5 threshold in Disconnectome analyses) and 0 otherwise. This binary mask for each domain was then used as seed for a disconnectome analysis (Threshold = 0.0 to capture the full range of connection probability). The resulting (dis)connectome map was then parcellated using the cortical Harvard-Oxford atlas, so that each portion of the atlas was associated with the average structural connectivity with the callosal location estimated to be the projection of a particular function. For each domain, parcellated structural connectomes were correlated with parcellated functional associations derived from Neurosynth (Figure 1.6).

In addition, in seeking further convergent validation of our findings by also considering a neuropsychological perspective, we conducted a complementary analysis based on the atlas of brain disconnections associated with neuropsychological test scores in a population of stroke patients by Talozzi et al. (2023). This complementary validation was run for the domains corresponding to the keywords “Motor”, “Attention”, “Episodic Memory”, “Language Comprehension” and “Speech Production” (for details, see Supplementary Materials).

### Estimation of functional gradients

Linear and quadratic functional gradients for validated functional domains were eventually computed to explore whether, within each cognitive domain, functional projections to the corpus callosum show a rostro-caudal or a dorso-ventral pattern. An example of significant linear rostro-caudal gradient for a given cognitive domain would entail stronger projections of the functions anteriorly than posteriorly within the callosum. An example of significant linear dorso-ventral gradient for a given cognitive domain would entail stronger projections of the functions dorsally than ventrally within the callosum. This was accomplished by estimating the link between each location (within the medial sagittal slice of the callosum) along the antero-posterior and dorso-ventral axis and the average functional projection of each function. This was done by means of generalized linear models with gamma distribution and logarithmic link function.

## Results

### Structural gradients

We observed both a linear (t= 11.681, p< 0.001) and a quadratic (t=2.941, p=0.005) antero-posterior gradient in structural connections. We also detected a linear association between the point of maximal connectivity along the antero-posterior axis and the location of the ROI along the dorso-ventral axis (t= 2.368, p= 0.023; Table 1; Figure 2a). In addition, a linear dorso-ventral gradient was observed (t= 4.446, p< 0.001), complemented by a quadratic association highlighting that the most anterior and posterior cortices have interhemispheric connections involving the most ventral part of the corpus callosum (t= −4.640, p< 0.001; Table 2; Figure 2b).

**Figure 2.**
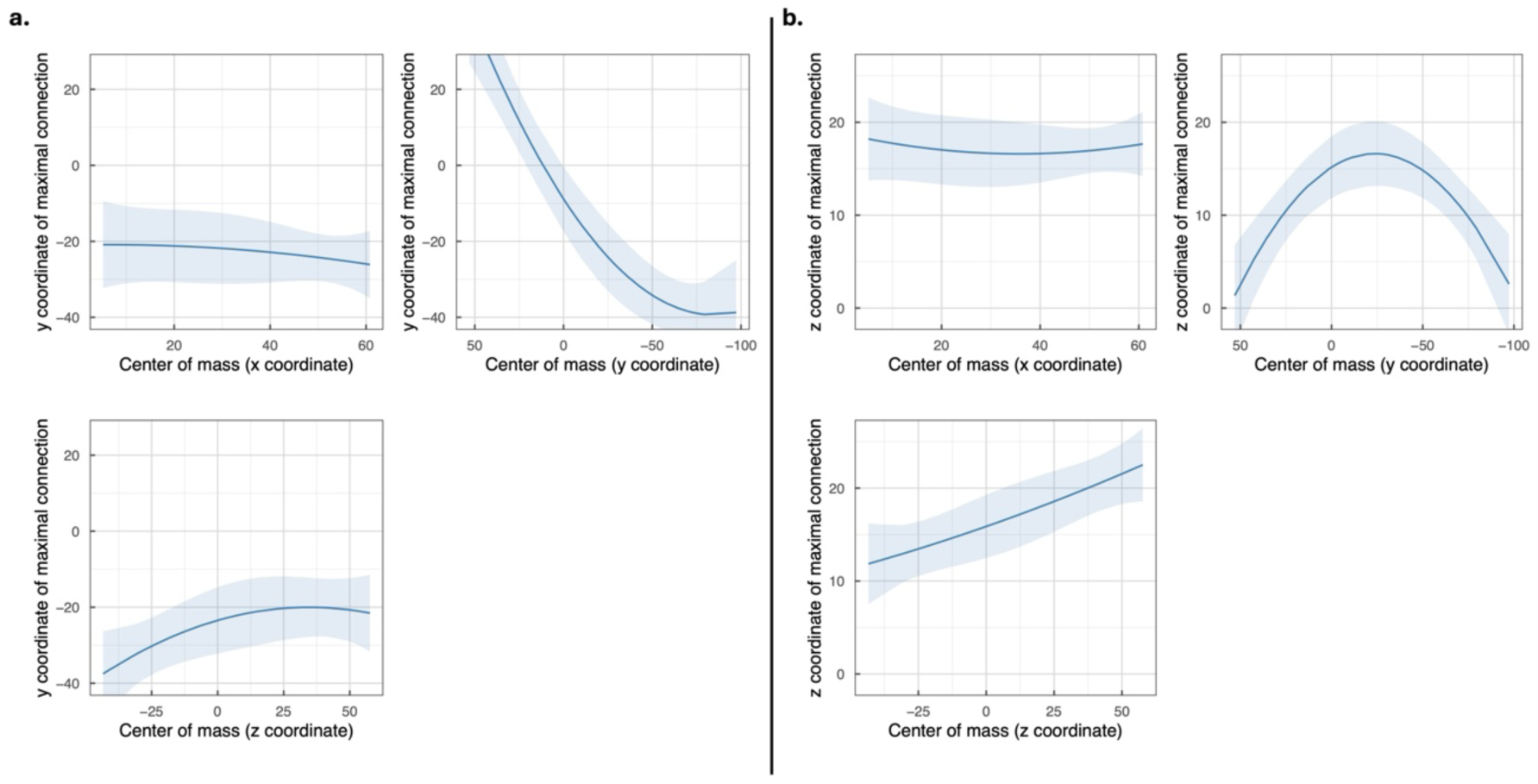
Estimated association between the stereotactic coordinates of the center of mass (COM) of each ROI and the location of maximal connection probability along the antero-posterior axis (a) or dorso-ventral axis (b). Values in the lateral-medial axis (x coordinates) that are closer to zero indicate areas closer to the sagittal midline. Negative values in the antero-posterior axis (y coordinates) indicate more posterior areas. Negative values in the dorso-ventral axis (z coordinates) indicate more ventral areas.

**Table 1.** Associations between location of maximal connection probability along the antero-posterior (y) axis and the x, y, and z coordinates of the center of mass of each ROI. Statistically significant effects (p< 0.05) are highlighted with an asterisk.

|  | Estimate | Std. Error | t value | p value |
| --- | --- | --- | --- | --- |
| (Intercept) | -19.333 | 1.593 | -12.140 | < 0.001* |
| ROI center of mass (x) : linear | -11.029 | 13.440 | -0.821 | 0.417 |
| ROI center of mass (x) : quadratic | -3.118 | 11.681 | -0.267 | 0.791 |
| ROI center of mass (y) : linear | 131.861 | 11.289 | 11.681 | < 0.001* |
| ROI center of mass (y) : quadratic | 41.265 | 14.029 | 2.941 | 0.005* |
| ROI center of mass (z) : linear | 26.950 | 11.381 | 2.368 | 0.023* |
| ROI center of mass (z) : quadratic | -15.397 | 12.847 | -1.198 | 0.238 |

**Table 2.**
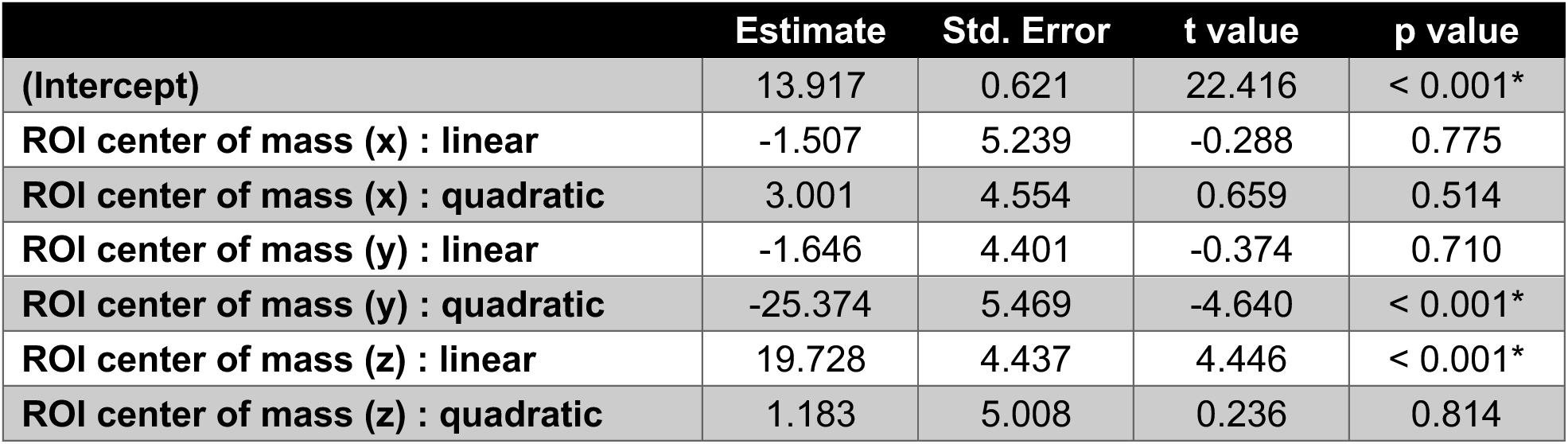
Associations between location of maximal connection probability along the dorso-ventral (z) axis and the x, y, and z coordinates of the center of mass of each ROI. Statistically significant effects (p< 0.05) are highlighted with an asterisk.

### Functional validation

Successful validation for functional projections in the corpus callosum was achieved for the “Visual” (ρ= 0.393, p= 0.003), “Somatosensory” (ρ= 0.287, p= 0.024), “Motor” (ρ= 0.598, p< 0.001), “Episodic memory” (ρ= 0.310, p= 0.016), “Working Memory” (ρ= 0.375, p= 0.004), and “Decision making” (ρ= 0.381, p= 0.004) domains. In all other domains (“Emotions”, “Auditory”, “Semantic”, “Language Comprehension”, “Speech Production”, “Attention”) validation did not reach statistical significance (Table 3). Only domains in which the correlation between structural and functional estimates were positive and significant were further considered and discussed. We note that there is no significant correlation between validation success and number of studies included in the Neurosynth meta-analysis (ρ (10)= 0.403, p= 0.194). This rules out the possibility that our validation approach could be biased by the number of studies available for functional meta-analyses.

**Table 3.**
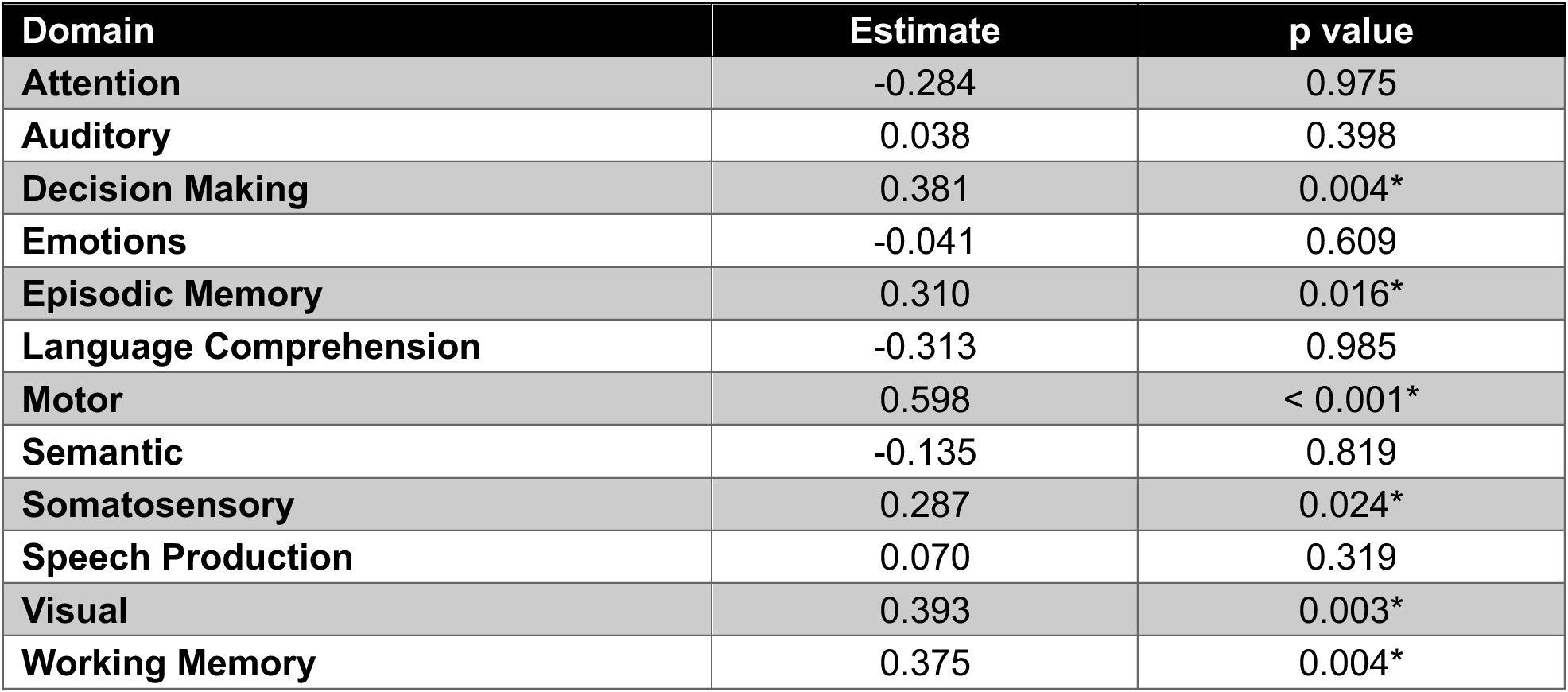
Results of the validation analysis (Spearman correlations). Statistically significant effects (p< 0.05) are highlighted with an asterisk.

| Domain | Estimate | p value |
| --- | --- | --- |
| Attention | -0.284 | 0.975 |
| Auditory | 0.038 | 0.398 |
| Decision Making | 0.381 | 0.004* |
| Emotions | -0.041 | 0.609 |
| Episodic Memory | 0.310 | 0.016* |
| Language Comprehension | -0.313 | 0.985 |
| Motor | 0.598 | < 0.001* |
| Semantic | -0.135 | 0.819 |
| Somatosensory | 0.287 | 0.024* |
| Speech Production | 0.070 | 0.319 |
| Visual | 0.393 | 0.003* |
| Working Memory | 0.375 | 0.004* |

For the results of the complementary neuropsychological validation, please see Supplementary Material and Supplementary Table S1.

### Functional gradients

We characterized significant linear and quadratic rostro-caudal (y axis) and dorso-ventral (z axis) functional gradients for keywords for which validation (i.e., back-projecting function from structure; see Functional validation paragraph) was successful (Figure 3). All linear and quadratic functional gradients turned out to be significant for both the antero-posterior (min |t| = 3.693, p< 0.001; Table 4) and the dorso-ventral axis (min |t| = 2.657, p= 0.013; Table 5). The “Visual” and “Episodic memory” keywords were associated with the inferior, middle and superior portion of the most posterior section of the callosum (i.e., the Splenium). The “Somatosensory” and “Motor” keywords were associated with regions of the callosum between the Splenium, the Isthmus, and the Body, with the former keyword being associated with more posterior callosal regions than the latter. The “Decision Making” keyword was associated with the most anterior portions of the callosum (Genu and Rostrum). The keyword “Working Memory” showed association with the Splenium and more anterior sections (from the Body onwards).

**Figure 3:**
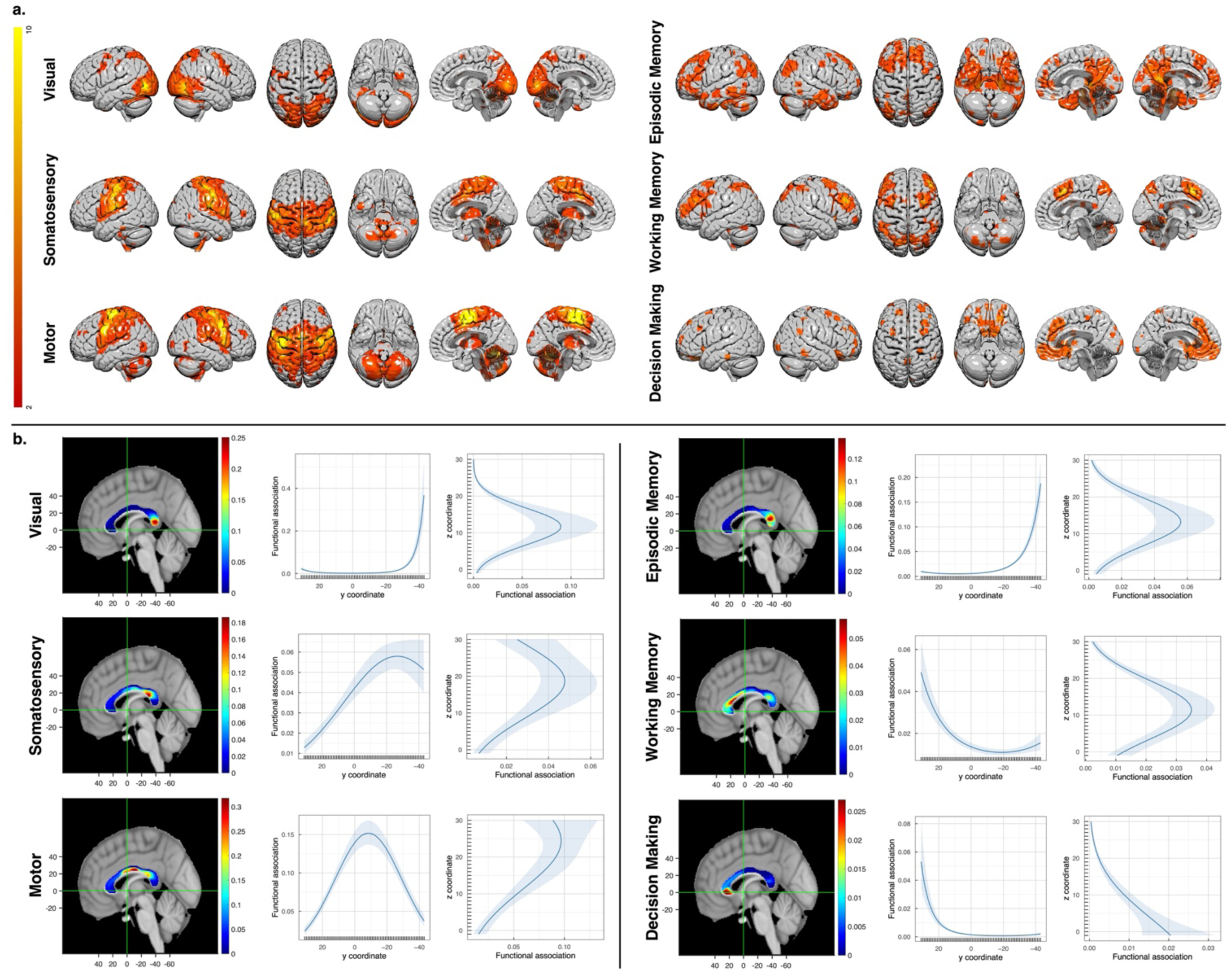
(a) meta-analytic maps as extracted from Neurosynth (the colorbar indicates Z scores of the association with the keyword). (b) Estimated projection of each cognitive function through the callosum, together with the estimated link between the functional projection and location along the antero-posterior axis and the dorso-ventral axis. These plots indicate the average magnitude of the functional projection of each given function on the corpus callosum for each point along the rostro-caudal and dorso-ventral axes. Negative values in the antero-posterior axis (y coordinates) indicate more posterior areas. Negative values in the antero-posterior axis (z coordinates) indicate more ventral areas.

**Table 4.** Functional gradients analysis: link between the y coordinate in the callosum and average functional projection of the validated cognitive functions. Statistically significant effects (p< 0.05) are highlighted with an asterisk.

|  |  | Estimate | Std. Error | t value | p value |
| --- | --- | --- | --- | --- | --- |
| **Visual** | (Intercept) | -4.929 | 0.065 | -76.270 | <0.001* |
|  | y coordinate : linear | -7.079 | 0.560 | -12.650 | <0.001* |
|  | y coordinate : quadratic | 10.209 | 0.560 | 18.240 | <0.001* |
| **Somatosensory** | (Intercept) | -3.254 | 0.043 | -75.461 | <0.001* |
|  | y coordinate : linear | -3.514 | 0.373 | -9.411 | <0.001* |
|  | y coordinate : quadratic | -1.648 | 0.373 | -4.412 | <0.001* |
| **Motor** | (Intercept) | -2.444 | 0.034 | -71.035 | <0.001* |
|  | y coordinate : linear | -1.100 | 0.298 | -3.693 | <0.001* |
|  | y coordinate : quadratic | -4.272 | 0.298 | -14.338 | <0.001* |
| **Episodic Memory** | (Intercept) | -4.372 | 0.038 | -114.150 | <0.001* |
|  | y coordinate : linear | -7.536 | 0.332 | -22.720 | <0.001* |
|  | y coordinate : quadratic | 4.893 | 0.332 | 14.750 | <0.001* |
| **Working Memory** | (Intercept) | -4.126 | 0.045 | -91.416 | <0.001* |
|  | y coordinate : linear | 2.903 | 0.391 | 7.427 | <0.001* |
|  | y coordinate : quadratic | 2.179 | 0.391 | 5.576 | <0.001* |
| **Decision Making** | (Intercept) | -6.070 | 0.073 | -83.519 | <0.001* |
|  | y coordinate : linear | 8.299 | 0.629 | 13.185 | <0.001* |
|  | y coordinate : quadratic | 6.037 | 0.629 | 9.592 | <0.001* |

**Table 5.** Functional gradients analysis: link between the z coordinate in the callosum and average functional projection of the validated cognitive functions. Statistically significant effects (p< 0.05) are highlighted with an asterisk.

|  |  | Estimate | Std. Error | t value | p value |
| --- | --- | --- | --- | --- | --- |
| **Visual** | (Intercept) | -4.284 | 0.117 | -36.596 | <0.001* |
|  | z coordinate : linear | -5.842 | 0.662 | -8.821 | <0.001* |
|  | z coordinate : quadratic | -8.690 | 0.662 | -13.122 | <0.001* |
| **Somatosensory** | (Intercept) | -3.554 | 0.096 | -37.139 | <0.001* |
|  | z coordinate : linear | 2.174 | 0.541 | 4.017 | <0.001* |
|  | z coordinate : quadratic | -2.147 | 0.541 | -3.967 | <0.001* |
| **Motor** | (Intercept) | -2.859 | 0.070 | -40.665 | <0.001* |
|  | z coordinate : linear | 2.954 | 0.398 | 7.427 | <0.001* |
|  | z coordinate : quadratic | -1.226 | 0.398 | -3.084 | 0.004* |
| **Episodic Memory** | (Intercept) | -3.951 | 0.103 | -38.361 | <0.001* |
|  | z coordinate : linear | -1.618 | 0.583 | -2.777 | 0.010* |
|  | z coordinate : quadratic | -5.312 | 0.583 | -9.117 | <0.001* |
| **Working Memory** | (Intercept) | -4.092 | 0.067 | -61.255 | <0.001* |
|  | z coordinate : linear | -2.511 | 0.378 | -6.644 | <0.001* |
|  | z coordinate : quadratic | -3.371 | 0.378 | -8.918 | <0.001* |
| **Decision Making** | (Intercept) | -5.486 | 0.075 | -72.894 | <0.001* |
|  | z coordinate : linear | -6.752 | 0.426 | -15.860 | <0.001* |
|  | z coordinate : quadratic | -1.131 | 0.426 | -2.657 | 0.013* |

## Discussion

In this study, we combined a meta-analytical functional approach (Yarkoni et al., 2011) with a neuroimaging-based human structural connectome technique (Foulon et al., 2018) to chart the functional involvement of the corpus callosum in different cognitive domains.

We replicated previous findings suggesting the existence of a rostro-caudal gradient in the structural connections of the callosum (Barakovic et al., 2021; Friedrich et al., 2020; Hofer & Frahm, 2006). We also characterized a dorso-ventral gradient (in line with Xiong et al., 2024) and an antero-posterior/rostro-caudal link, whereby connections of the most anterior and posterior cortices involve the most ventral parts of the corpus callosum. These structure-only data point towards the existence of anatomical regularities in callosal anatomical connectivity, which complement previous ROI-based studies that outlined the connections of the corpus callosum with macro-anatomical cortical areas (Catani et al., 2012).

The observed task-independent anatomical gradients constitute ideal foundations for the exploration of the hypothetically graded involvement of different portions of the corpus callosum in different aspects of cognition. In this regard, the groundwork of this endeavour was laid down by Friedrich and colleagues (2020), who highlighted that posterior portions of the corpus callosum connect lower-level sensorimotor cortices, while more anterior portions of the callosum connect areas involved in higher-order functions.

Previous diffusion MRI data have suggested the involvement of the callosum in some cognitive domains: the literature has highlighted the involvement of central-anterior portions of the callosum in emotion recognition (Orlando et al., 2023), of the callosal *genu* in motor performance (Hannanu et al., 2022), of the precentral portions in sensorimotor synchronization (Blecher, Tal & Ben-Shachar, 2016), of anterior, central and posterior portions in attention (Chechlacz et al., 2015; Mesaros et al., 2009), and of central-posterior regions in vocabulary and matrix reasoning (Danielsen et al., 2020).This evidence is complemented by the observation that the corpus callosum displays converging patterns of structural and functional connectivity towards white-matter functional networks (Peer et al., 2017; Wang et al, 2020; Wang et al., 2021).

On the other hand, the *Functionnectome* framework (Nozais et al., 2021; Nozais et al., 2023) has revealed that it is possible to infer brain functioning at the network level, by projecting neurofunctional data onto structural connections. Here we built on these theoretical and methodological pillars to estimate the participation of different portions of the corpus callosum in different aspects of cognition, by projecting meta-analytical functional data onto probabilistic maps of structural connections. By doing so, we produced the first large-scale probabilistic outline of the functional anatomy of the corpus callosum, through which we were able to highlight that that vision and episodic memory involve the most posterior portions of the callosum, somatosensory and motor functions involve the central dorsal portion, whereas decision making and working memory involve anterior fibers.

Critically, these findings extend previous gradient- and connectivity-based descriptions of the corpus callosum in a fundamental way. Existing gradient-based evidence has established how callosal fibres are spatially organized and the order of cognitive function they subserve (Friedrich et al., 2020; Xiong et al., 2024), whereas studies on structural and functional connectivity have demonstrated their participation in large-scale intrinsic networks (Peer et al., 2017; Wang et al., 2020; Wang et al., 2021). The present framework instead assigns voxel-wise cognitive significance to callosal structures by projecting multiple meta-analytically defined cognitive domains onto a common structural reference. This allows the relative functional organization of different cognitive operations to be characterized within a unified probabilistic framework, moving beyond isolated structure–function associations towards a comprehensive atlas of the functional topography of the corpus callosum. Importantly, this approach also overcomes a key limitation of diffusion MRI-based accounts, which typically infer functional involvement from correlations between white-matter structure (e.g., fractional anisotropy) and behavioural performance on individual tasks, thereby restricting inference to task-specific effects that are not immediately comparable across cognitive domains.

We note that data originating functional projections come from functional neuroimaging, which limits any conclusive claim on specificity and exclusivity of the present results. Still, our findings align with existing neuropsychological evidence (see also our neuropsychological validation, Supplementary Material and Supplementary Table S1). Indeed, splenial lesions have been extensively associated with (often comorbid) clinical conditions involving the visual domain, like hemianopia, color anomia, deficits in binocular stereopsis, pure alexia, and optic aphasia (Berlucchi & Aglioti, 2020; Bonandrini et al., 2020; Coslett & Saffran, 1989; Epelbaum et al., 2008; Geschwind & Fusillo, 1966; Luzzatti, 2003; Luzzatti et al., 1998; Montant & Behrmann, 2000; Saffran & Coslett, 1998; Veronelli et al., 2024). Although the magnitude of the involvement of the splenium in different computational subcomponents of visual perception is yet to be quantified, it was hypothesized that this portion of the corpus callosum mediates integration and synchronization of the activity of the two contralateral visual cortices, and it plays a critical role in the development of the alpha (8-12 Hz) rhythm (for a review, see Knyazeva, 2013). From a more cognitive standpoint, recent evidence suggests that the splenium might be involved in automatic integration of visual information targeting the two visual hemifields (Pinto et al., 2023).

As for sensorimotor functions, our results comply with evidence linking left tactile agnosia and apraxia with lesions of the body of the callosum (Balsamo et al., 2008; Graff-Radford et al., 1987; Heilman & Watson, 2008). We hypothesize this portion of the callosum to be involved in the minimisation of contralateral sensorimotor conflict (whose failure is associated with the so-called alien hand syndrome; Geschwind et al., 1995). Indeed, evidence from previous literature suggests that callosal fibres are involved in inter-hemispheric inhibition across homologous contralateral areas of the motor system (Ferbert et al., 1992), and that they mediate activation of the posterior parietal and parietal opercular cortices in the hemisphere ipsilateral to a hand undergoing tactile stimulation (even though activation is predominantly contralateral to the side of tactile stimulation; Fabri et al., 1999). The association between the present results and previous neuropsychological literature is less direct for working memory and decision making: data from Bechara and colleagues (Bechara et al., 1998) suggest the severing of anterior callosal fibers in at least one of their patients with working memory and decision-making deficits. Yet, the lack of systematic support for this claim in the literature suggests that a callosal lesion alone may not be sufficient to cause deficits in these domains (we assume that this lack of convergence between our data and the neuropsychological literature could to some extent depend on a relatively limited functional lateralization of these cognitive domains). Nevertheless, previous data suggest a link between working memory and diffusion MRI-based fractional anisotropy measures in the anterior part of the body of the corpus callosum (Takeuchi et al., 2010). In this regard, it is possible to hypothesize that the corpus callosum plays a critical role in allowing the bi-hemispheric communication in the fronto-parietal network that supports the computational underpinnings of verbal and spatial working memory (Treble et al., 2013). Anterior portions of the corpus callosum have also been related to the pathophysiology of impulsive and suicidal behaviors in bipolar disorder (Matsuo et al., 2010). From a more cognitive perspective, these callosal fibers are deemed to play a critical role in the resolution of interhemispheric inhibition in tasks requiring competition between conflicting representations (Schulte & Müller-Oehring, 2010).

As for episodic memory, retro-splenial lesions - that are most likely to affect splenial connections - have been associated to memory deficits (Li et al., 2018; Saito et al., 2003; Valenstein et al., 1987; Yasuda et al., 1997). In addition, Overman et al (2021) observed that verbal retrieval deficits in a group of patients with posterior cortical atrophy was associated with microstructural damage of the splenium of the corpus callosum. Overman et al. interpreted this finding considering the structural connections between the splenium and thalamic radiations and the internal capsule and hypothesized that the splenium may play a role in attentional mechanisms subserving memory functions.

It is interesting to note that reliable structural-functional associations were only detected for a relatively limited subset of functions. Notably, linguistic functions are not among these. This may depend on our validation procedure being too conservative to capture more subtle structure-functional associations: here we assumed that the connectivity from the estimated functional callosal locus to the cerebral cortex is proportional to the degree of involvement of each cortical region in that function. This assumption may simply not fit all cognitive functions in the same way, also depending on their focal or distributed organization. This may be particularly relevant for linguistic functions, which tend to be lateralized to the left hemisphere (whereas structural back-projections from the corpus callosum are inherently bilateral), although ample variability exists between different subcomponents (see for instance Bonandrini et al., 2024; Branch et al., 1964; Gerrits et al., 2019; Parker et al., 2024; Parker et al., 2022; Vigneau et al., 2011).

As for the similar pattern of results in the attention domain, we suggest that the whole corpus callosum is likely to be a crucial anatomical structure supporting attention (Chechlacz et al., 2015; Mesaros et al., 2009), which makes it difficult to isolate specific subsections for which reliable bi-directional structure-functional associations may take place. More specifically, we note that functional projections for the cognitive domain of attention produced a localised pattern in posterior callosal regions but not in anterior ones (which theoretically connect the frontal areas). We believe this may have played a critical role in the negative validation result we obtained.

On the other hand, these results could suggest that structural-functional projections in the callosum can only be established for relatively basic cognitive functions, which tend to be more precisely localized in the cerebral cortex. The use of meta-analytical data prevents us from drawing conclusive inference, since the different number of studies included in each meta-analysis makes them not comparable. Still, the adoption of meta-analytical data assures robustness of the functional associations, while minimizing the chance that results could depend on the idiosyncrasies of any given empirical study. It is also worth noting that we also double-checked the consistency of our results, by conducting a confirmatory analysis based upon the asymmetric atlas by Schaefer et al (2018), as an alternative to the symmetric cortical Harvard-Oxford atlas used in the main analysis, through which we obtained virtually identical results (which excludes our results to depend on the cortical parcellation approach; see Supplementary Tables S2-S7 and supplementary Figures S1 and S2).

In conclusion, our study revealed that the posterior portion of the callosum is involved in vision and episodic memory, that the central, most dorsal part is involved in somatosensory and motor functions, and that more anterior portions are involved in working memory and decision making. Although historically considered a structural scaffolding device needed to prevent the two cerebral hemispheres from collapsing onto one another (Mooshagian, 2008), the corpus callosum has been characterized by contemporary neuroscience as rather fundamental in human cognition. More precisely, in the second half of the 20^th^ century interest towards the involvement of the callosum in cognition was ignited by research on split-brain patients (Berlucchi & Aglioti, 2020; Berlucchi et al., 1997; Cohen et al., 2000; Gazzaniga, 1995; LeDoux & Gazzaniga, 1981; Levy & Trevarthen, 1977; Metcalfe et al., 1995; Zaidel, 1973; Zaidel, 1982; Zaidel, 1983; Zaidel et al., 1999; Santander et al., 2025; Bekir et al., 2025), which paved the way for the contemporary study of functional brain lateralization (e.g., Behrmann & Plaut, 2015; Bonandrini et al., 2024; Bonandrini et al., 2023; Bryden, 1988; Davidson, 2014; Dundas et al., 2013; Gerrits et al., 2020; Gerrits et al., 2019; Ley & Bryden, 1979; Ocklenburg et al., 2024; Parker et al., 2024; Parker et al., 2022; Plaut & Behrmann, 2011; Vigneau et al., 2011; Vingerhoets et al., 2012; Vingerhoets & Stroobant, 1999). The present work seeks to complement this tradition with the localization of projections of different cognitive functions onto specific portions of the corpus callosum. We believe that these results may help tailor more sensitive clinical neuropsychological assessments in case of callosal lesions. On the other hand, the present study can inform future anatomically-plausible models of cognitive processes, as well as novel empirical neuroimaging and brain stimulation studies.

## Footnotes

1 An alternative anatomical subdivision of the corpus callos um is the one entailing forceps major (u-shaped bundle originating from the splenium and running posteriorly), forceps minor (bundle originating from the genu and connecting frontal areas), body and tapetum (portion of the corpus callosum that constitutes the superior boundary of the lateral ventricles.

2 Here and henceforth, by “gradient”, we define (in line with Bernhardt et al., 2022) “spatial trends of neural organization in which structure-function relationships can be investigated”.

3 Given our interest towards the link between the corpus callosum and cortical areas, we adopted a reference atlas that does not include cerebellum and subcortical structures (e.g., basal ganglia)

4 COM coordinates were averaged between each couple of contralateral homologous areas.

5 if multiple tied maxima were found, the one located closer to the origin of the MNI space was chosen for further analyses, in order to minimize spatial inflation of results.

## References

Aboitiz, F., Scheibel, A. B., Fisher, R. S., & Zaidel, E. (1992). Fiber composition of the human corpus callosum. Brain research, 598(1-2), 143–153.

Ashburner, J., Barnes, G., Chen, C. C., Daunizeau, J., Flandin, G., Friston, K., … & Penny, W. (2014). SPM12 manual. Wellcome Trust Centre for Neuroimaging, London, UK, 2464(4), 53.

Balsamo, M., Trojano, L., Giamundo, A., & Grossi, D. (2008). Left hand tactile agnosia after posterior callosal lesion. Cortex, 44(8), 1030–1036.

Barbaresi, P., Fabri, M., Lorenzi, T., Sagrati, A., & Morroni, M. (2024). Intrinsic organization of the corpus callosum. Frontiers in Physiology, 15, 1393000.

Barakovic, M., Girard, G., Schiavi, S., Romascano, D., Descoteaux, M., Granziera, C.,…Daducci, A. (2021). Bundle-specific axon diameter index as a new contrast to differentiate white matter tracts. Frontiers in neuroscience, 15, 646034.

Bechara, A., Damasio, H., Tranel, D., & Anderson, S. W. (1998). Dissociation of working memory from decision making within the human prefrontal cortex. Journal of neuroscience, 18(1), 428–437.

Behrmann, M., & Plaut, D. C. (2015). A vision of graded hemispheric specialization. Ann N Y Acad Sci, 1359, 30–46. 10.1111/nyas.12833

Bekir, S., Hopf, J. L., Paul, T., Wiemer, V. M., Santander, T., Skinner, H. E., … & Miller, M. B. (2025). No disconnection syndrome after near-complete callosotomy. Communications Psychology.

Berlucchi, G., & Aglioti, S. (2020). Interhemispheric disconnection syndromes. In Handbook of clinical and experimental neuropsychology (pp. 635–670). Psychology Press.

Berlucchi, G., Mangun, G., & Gazzaniga, M. (1997). Visuospatial attention and the split brain. Physiology, 12(5), 226–231.

Bernhardt, B. C., Smallwood, J., Keilholz, S., & Margulies, D. S. (2022). Gradients in brain organization. NeuroImage, 251, 118987.

Blecher, T., Tal, I., & Ben-Shachar, M. (2016). White matter microstructural properties correlate with sensorimotor synchronization abilities. NeuroImage, 138, 1–12.

Blits, K. C. (1999). Aristotle: form, function, and comparative anatomy. The Anatomical Record: An Official Publication of the American Association of Anatomists, 257(2), 58–63.

Bloom, J. S., & Hynd, G. W. (2005). The role of the corpus callosum in interhemispheric transfer of information: excitation or inhibition? Neuropsychology review, 15, 59–71.

Bonandrini, R., Gornetti, E., & Paulesu, E. (2024). A meta-analytical account of the functional lateralization of the reading network. cortex, 177, 363–384.

Bonandrini, R., Paulesu, E., Traficante, D., Capelli, E., Marelli, M., & Luzzatti, C. (2023). Lateralized reading in the healthy brain: A behavioral and computational study on the nature of the visual field effect. Neuropsychologia, 108468.

Bonandrini, R., Veronelli, L., Licciardo, D., Caporali, A., Judica, E., Corbo, M., & Luzzatti, C. (2020). Can the right hemisphere read? A behavioral and disconnectome study on implicit reading in a patient with pure alexia. Neurocase, 26(6), 321–327. 10.1080/13554794.2020.1830118

Bozzali, M., Mastropasqua, C., Cercignani, M., Giulietti, G., Bonnì, S., Caltagirone, C., & Koch, G. (2012). Microstructural damage of the posterior corpus callosum contributes to the clinical severity of neglect. PloS one, 7(10), e48079.

Branch, C., Milner, B., & Rasmussen, T. (1964). Intracarotid sodium amytal for the lateralization of cerebral speech dominance: observations in 123 patients. Journal of neurosurgery, 21(5), 399–405.

Bryden, M. P. (1988). An overview of the dichotic listening procedure and its relation to cerebral organization.

Caleo, M. (2018). Plasticity of transcallosal pathways after stroke and their role in recovery. The Journal of Physiology, 596(10), 1789.

Catani, M. (2007). From hodology to function. Brain, 130(3), 602–605.

Catani, M., Schotten, M. T. d., & Thiebaut de Schotten, M. (2012). Commissural Pathways. In Atlas of Human Brain Connections (pp. 0). Oxford University Press. 10.1093/med/9780199541164.003.0105

Catani, M., & Thiebaut de Schotten, M. (2012). Atlas of human brain connections. Oxford University Press, USA.

Chechlacz, M., Humphreys, G. W., Sotiropoulos, S. N., Kennard, C., & Cazzoli, D. (2015). Structural organization of the corpus callosum predicts attentional shifts after continuous theta burst stimulation. Journal of Neuroscience, 35(46), 15353–15368.

Cohen, L., Dehaene, S., Naccache, L., Lehéricy, S., Dehaene-Lambertz, G., Hénaff, M. A., & Michel, F. (2000). The visual word form area: spatial and temporal characterization of an initial stage of reading in normal subjects and posterior split-brain patients. Brain, 123 (Pt 2), 291–307.

Coslett, H. B., & Saffran, E. M. (1989). Evidence for preserved reading in ‘pure alexia’. Brain, 112 (Pt 2), 327–359. 10.1093/brain/112.2.327

D’Arcy, R. C., Hamilton, A., Jarmasz, M., Sullivan, S., & Stroink, G. (2006). Exploratory data analysis reveals visuovisual interhemispheric transfer in functional magnetic resonance imaging. Magnetic Resonance in Medicine: An Official Journal of the International Society for Magnetic Resonance in Medicine, 55(4), 952–958.

Danielsen, V. M., Vidal-Piñeiro, D., Mowinckel, A. M., Sederevicius, D., Fjell, A. M., Walhovd, K. B., & Westerhausen, R. (2020). Lifespan trajectories of relative corpus callosum thickness: regional differences and cognitive relevance. Cortex, 130, 127–141.

Davidson, R. J. (2014). Hemispheric asymmetry and emotion. In Approaches to emotion (pp. 39-57). Psychology Press.

Desikan, R. S., Ségonne, F., Fischl, B., Quinn, B. T., Dickerson, B. C., Blacker, D.,…Hyman, B. T. (2006). An automated labeling system for subdividing the human cerebral cortex on MRI scans into gyral based regions of interest. Neuroimage, 31(3), 968–980.

Dolan, R. J. (2008). Neuroimaging of cognition: past, present, and future. Neuron, 60(3), 496–502.

Dundas, E. M., Plaut, D. C., & Behrmann, M. (2013). The joint development of hemispheric lateralization for words and faces. Journal of Experimental Psychology: General, 142(2), 348.

Epelbaum, S., Pinel, P., Gaillard, R., Delmaire, C., Perrin, M., Dupont, S.,…Cohen, L. (2008). Pure alexia as a disconnection syndrome: new diffusion imaging evidence for an old concept. cortex, 44(8), 962–974.

Fabri, M., Polonara, G., Mascioli, G., Salvolini, U., & Manzoni, T. (2011). Topographical organization of human corpus callosum: an fMRI mapping study. Brain research, 1370, 99–111.

Fabri, M., Polonara, G., Quattrini, A., Salvolini, U., Del Pesce, M., & Manzoni, T. (1999). Role of the corpus callosum in the somatosensory activation of the ipsilateral cerebral cortex: an fMRI study of callosotomized patients. European Journal of Neuroscience, 11(11), 3983–3994.

Ferbert, A., Priori, A., Rothwell, J. C., Day, B. L., Colebatch, J. G., & Marsden, C. (1992). Interhemispheric inhibition of the human motor cortex. The Journal of physiology, 453(1), 525–546.

Fesl, G., Braun, B., Rau, S., Wiesmann, M., Ruge, M., Bruhns, P.,…Tonn, J.-C. (2008). Is the center of mass (COM) a reliable parameter for the localization of brain function in fMRI? European radiology, 18, 1031–1037.

Foulon, C., Cerliani, L., Kinkingnehun, S., Levy, R., Rosso, C., Urbanski, M.,…Thiebaut de Schotten, M. (2018). Advanced lesion symptom mapping analyses and implementation as BCBtoolkit. Gigascience, 7(3), giy004.

Frazier, J. A., Chiu, S., Breeze, J. L., Makris, N., Lange, N., Kennedy, D. N.,…Dieterich, M. E. (2005). Structural brain magnetic resonance imaging of limbic and thalamic volumes in pediatric bipolar disorder. American Journal of Psychiatry, 162(7), 1256–1265.

Friedrich, P., Forkel, S. J., & de Schotten, M. T. (2020). Mapping the principal gradient onto the corpus callosum. Neuroimage, 223, 117317.

Friston, K. J. (2009). Modalities, modes, and models in functional neuroimaging. Science, 326(5951), 399–403.

Frith, C. D., & Friston, K. J. (2013). Studying brain function with neuroimaging. In Cognitive neuroscience (pp. 169–195). Psychology Press.

Gawryluk, J. R., D’Arcy, R. C., Mazerolle, E. L., Brewer, K. D., & Beyea, S. D. (2011). Functional mapping in the corpus callosum: A 4 T fMRI study of white matter. Neuroimage, 54(1), 10–15.

Gawryluk, J. R., Mazerolle, E. L., & D’Arcy, R. C. (2014). Does functional MRI detect activation in white matter? A review of emerging evidence, issues, and future directions. Frontiers in neuroscience, 8, 239.

Gazzaniga, M. S. (1995). Principles of human brain organization derived from split-brain studies. Neuron, 14(2), 217–228.

Gazzaniga, M. S. (2000). Cerebral specialization and interhemispheric communication: does the corpus callosum enable the human condition? Brain, 123(7), 1293–1326.

Gazzaniga, M. S. (2005). Forty-five years of split-brain research and still going strong. Nat Rev Neurosci, 6(8), 653–659. 10.1038/nrn1723

Gerrits, R., De Clercq, P., Verhelst, H., & Vingerhoets, G. (2020). Evaluating the performance of the visual half field paradigm as a screening tool to detect right hemispheric language dominance. Laterality, 25(6), 722–739.

Gerrits, R., Van der Haegen, L., Brysbaert, M., & Vingerhoets, G. (2019). Laterality for recognizing written words and faces in the fusiform gyrus covaries with language dominance. Cortex, 117, 196–204.

Geschwind, N., & Fusillo, M. (1966). Color-naming defects in association with alexia. Archives of Neurology, 15(2), 137–146.

Geschwind, D. H., Iacoboni, M., Mega, M. S., Zaidel, D. W., Cloughesy, T., & Zaidel, E. (1995). Alien hand syndrome: interhemispheric motor disconnection due to a lesion in the midbody of the corpus callosum. Neurology, 45(4), 802–808.

Goldstein, J. M., Seidman, L. J., Makris, N., Ahern, T., O’Brien, L. M., Caviness Jr, V. S.,…Tsuang, M. T. (2007). Hypothalamic abnormalities in schizophrenia: sex effects and genetic vulnerability. Biological psychiatry, 61(8), 935–945.

Graff-Radford, N. R., Welsh, K., & Godersky, J. (1987). Callosal apraxia. Neurology, 37(1), 100–100.

Hannanu, F. F., Naegele, B., Hommel, M., Krainik, A., Detante, O., & Jaillard, A. (2022). White matter tract disruption is associated with ipsilateral hand impairment in subacute stroke: a diffusion MRI study. Neuroradiology, 64(8), 1605–1615.

Heilman, K. M., & Watson, R. T. (2008). The disconnection apraxias. Cortex, 44(8), 975–982.

Helenius, J., Perkiö, J., Soinne, L., Østergaard, L., Carano, R. A., Salonen, O.,…Tatlisumak, T. (2003). Cerebral hemodynamics in a healthy population measured by dynamic susceptibility contrast MR imaging. Acta radiologica, 44(5), 538–546.

Hinkley, L. B., Marco, E. J., Findlay, A. M., Honma, S., Jeremy, R. J., Strominger, Z., … & Sherr, E. H. (2012). The Role of Corpus Callosum Development in Functional Connectivity and Cognitive Processing. PLoS ONE, 7(8), e39804.

Hofer, S., & Frahm, J. (2006). Topography of the human corpus callosum revisited—comprehensive fiber tractography using diffusion tensor magnetic resonance imaging. Neuroimage, 32(3), 989–994.

Hua, K., Zhang, J., Wakana, S., Jiang, H., Li, X., Reich, D. S.,…Mori, S. (2008). Tract probability maps in stereotaxic spaces: analyses of white matter anatomy and tract-specific quantification. Neuroimage, 39(1), 336–347.

Karolis, V. R., Corbetta, M., & Thiebaut de Schotten, M. (2019). The architecture of functional lateralisation and its relationship to callosal connectivity in the human brain. Nature communications, 10 (1), 1417.

Knyazeva, M. G. (2013). Splenium of corpus callosum: patterns of interhemispheric interaction in children and adults. Neural plasticity, 2013(1), 639430.

Le Bihan, D., Mangin, J. F., Poupon, C., Clark, C. A., Pappata, S., Molko, N., & Chabriat, H. (2001). Diffusion tensor imaging: concepts and applications. Journal of Magnetic Resonance Imaging: An Official Journal of the International Society for Magnetic Resonance in Medicine, 13(4), 534–546.

LeDoux, J. E., & Gazzaniga, M. S. (1981). The brain and the split brain: A duel with duality as a model of mind. Behavioral and Brain Sciences, 4(1), 109–110.

Levy, J., & Trevarthen, C. (1977). Perceptual, semantic and phonetic aspects of elementary language processes in split-brain patients. Brain, 100 Pt 1, 105–118. 10.1093/brain/100.1.105

Ley, R. G., & Bryden, M. P. (1979). Hemispheric differences in processing emotions and faces. Brain and language, 7(1), 127–138.

Li, P., Shan, H., Liang, S., Nie, B., Duan, S., Huang, Q.,…Ma, L. (2018). Structural and functional brain network of human retrosplenial cortex. Neuroscience Letters, 674, 24–29.

Li, Y., Wu, P., Liang, F., & Huang, W. (2015). The microstructural status of the corpus callosum is associated with the degree of motor function and neurological deficit in stroke patients. PLoS One, 10(4), e0122615.

Luzzatti, C. (2003). Optic aphasia and pure alexia: contribution of callosal disconnection syndromes to the study of lexical and semantic representation in the right hemisphere.

Luzzatti, C., Rumiati, R. I., & Ghirardi, G. (1998). A functional model of visuo-verbal disconnection and the neuroanatomical constraints of optic aphasia. Neurocase, 4(1), 71–87, 1355-4794.

Maier-Hein, K. H., Neher, P. F., Houde, J.-C., Côté, M.-A., Garyfallidis, E., Zhong, J.,…Ji, Q. (2017). The challenge of mapping the human connectome based on diffusion tractography. Nature communications, 8(1), 1349.

Makris, N., Goldstein, J. M., Kennedy, D., Hodge, S. M., Caviness, V. S., Faraone, S. V.,…Seidman, L. J. (2006). Decreased volume of left and total anterior insular lobule in schizophrenia. Schizophrenia research, 83(2-3), 155–171.

Matsuo, K., Nielsen, N., Nicoletti, M. A., Hatch, J. P., Monkul, E. S., Watanabe, Y.,…Soares, J. C. (2010). Anterior genu corpus callosum and impulsivity in suicidal patients with bipolar disorder. Neuroscience letters, 469(1), 75–80.

Mesaros, S., Rocca, M. A., Riccitelli, G., Pagani, E., Rovaris, M., Caputo, D., … & Filippi, M. (2009). Corpus callosum damage and cognitive dysfunction in benign MS. Human brain mapping, 30(8), 2656–2666.

Metcalfe, J., Funnell, M., & Gazzaniga, M. S. (1995). Right-hemisphere memory superiority: Studies of a split-brain patient. Psychological Science, 6(3), 157–164.

Montant, M., & Behrmann, M. (2000). Pure alexia. Neurocase, 6(4), 265–294.

Mooshagian, E. (2008). Anatomy of the corpus callosum reveals its function. Journal of Neuroscience, 28(7), 1535–1536.

Mori, S., Wakana, S., Van Zijl, P. C., & Nagae-Poetscher, L. (2005). MRI atlas of human white matter. Elsevier.

Nozais, V., Forkel, S. J., Foulon, C., Petit, L., & Thiebaut de Schotten, M. (2021). Functionnectome as a framework to analyse the contribution of brain circuits to fMRI. Communications biology, 4(1), 1035.

Nozais, V., Theaud, G., Descoteaux, M., Thiebaut de Schotten, M., & Petit, L. (2023). Improved Functionnectome by dissociating the contributions of white matter fiber classes to functional activation. Brain Structure and Function, 228(9), 2165–2177.

Ocklenburg, S., Mundorf, A., Gerrits, R., Karlsson, E. M., Papadatou-Pastou, M., & Vingerhoets, G. (2024). Clinical implications of brain asymmetries. Nature Reviews Neurology, 1–12.

Orlando, I., Ricci, C., Griffanti, L., & Filippini, N. (2023). Neural correlates of successful emotion recognition in healthy elderly: a multimodal imaging study. Social Cognitive and Affective Neuroscience, 18(1), nsad058.

Overman, M. J., Zamboni, G., Butler, C., & Ahmed, S. (2021). Splenial white matter integrity is associated with memory impairments in posterior cortical atrophy. Brain Communications, 3(2), fcab060.

Parker, A. J., Hontaru, M.-E., Lin, R., Ollerenshaw, S., & Bonandrini, R. (2024). Opposite perceptual biases in analogous auditory and visual tasks are unique to consonant–vowel strings and are unlikely a consequence of repetition. Laterality, 1–30.

Parker, A. J., Woodhead, Z. V., Carey, D. P., Groen, M. A., Gutierrez-Sigut, E., Hodgson, J.,…Payne, H. (2022). Inconsistent language lateralisation–Testing the dissociable language laterality hypothesis using behaviour and lateralised cerebral blood flow. cortex, 154, 105–134.

Peer, M., Nitzan, M., Bick, A. S., Levin, N., & Arzy, S. (2017). Evidence for functional networks within the human brain’s white matter. Journal of Neuroscience, 37(27), 6394–6407.

Perani, D. (2008). Functional neuroimaging of cognition. Handbook of Clinical Neurology, 88, 61–111.

Pinto, Y., Villa, M. C., Siliquini, S., Polonara, G., Passamonti, C., Lattanzi, S., … & de Haan, E. H. (2023). Visual integration across fixation: automatic processes are split but conscious processes remain unified in the split-brain. Frontiers in Human Neuroscience, 17, 1278025.

Plaut, D. C., & Behrmann, M. (2011). Complementary neural representations for faces and words: a computational exploration. Cogn Neuropsychol, 28(3-4), 251–275. 10.1080/02643294.2011.609812

Preibisch, C., & Haase, A. (2001). Perfusion imaging using spin-labeling methods: Contrast-to-noise comparison in functional MRI applications. Magnetic Resonance in Medicine: An Official Journal of the International Society for Magnetic Resonance in Medicine, 46(1), 172–182.

Price, C. J. (2012). A review and synthesis of the first 20 years of PET and fMRI studies of heard speech, spoken language and reading. Neuroimage, 62(2), 816–847. 10.1016/j.neuroimage.2012.04.062

Rostrup, E., Law, I., Blinkenberg, M., Larsson, H., Born, A. P., Holm, S., & Paulson, O. (2000). Regional differences in the CBF and BOLD responses to hypercapnia: a combined PET and fMRI study. Neuroimage, 11(2), 87–97.

Rode, G., Cotton, F., Revol, P., Jacquin-Courtois, S., Rossetti, Y., & Bartolomeo, P. (2010). Representation and disconnection in imaginal neglect. Neuropsychologia, 48(10), 2903–2911.

Saffran, E. M., & Coslett, H. B. (1998). Implicit vs. letter-by-letter reading in pure alexia: A tale of two systems. Cognitive Neuropsychology, 15(1-2), 141–165, 0264-3294.

Saito, K., Kimura, K., Minematsu, K., Shiraishi, A., & Nakajima, M. (2003). Transient global amnesia associated with an acute infarction in the retrosplenium of the corpus callosum. Journal of the Neurological Sciences, 210(1-2), 95–97.

Santander, T., Bekir, S., Paul, T., Simonson, J. M., Wiemer, V. M., Skinner, H. E., … & Miller, M. B. (2025). Full interhemispheric integration sustained by a fraction of posterior callosal fibers. Proceedings of the National Academy of Sciences, 122(43), e2520190122.

Salvalaggio, A., De Filippo De Grazia, M., Zorzi, M., Thiebaut de Schotten, M., & Corbetta, M. (2020). Post-stroke deficit prediction from lesion and indirect structural and functional disconnection. Brain, 143(7), 2173–2188.

Schaefer, A., Kong, R., Gordon, E. M., Laumann, T. O., Zuo, X. N., Holmes, A. J., … & Yeo, B. T. (2018). Local-global parcellation of the human cerebral cortex from intrinsic functional connectivity MRI. Cerebral cortex, 28(9), 3095–3114.

Shah, A., Jhawar, S., Goel, A., & Goel, A. (2021). Corpus callosum and its connections: a fiber dissection study. World Neurosurgery, 151, e1024–e1035.

Schulte, T., & Müller-Oehring, E. M. (2010). Contribution of callosal connections to the interhemispheric integration of visuomotor and cognitive processes. Neuropsychology review, 20(2), 174–190.

Sperry, R. W. (1964). The great cerebral commissure. Scientific American, 210(1), 42–53.

Takeuchi, H., Sekiguchi, A., Taki, Y., Yokoyama, S., Yomogida, Y., Komuro, N.,…Kawashima, R. (2010). Training of working memory impacts structural connectivity. Journal of Neuroscience, 30(9), 3297–3303.

Talozzi, L., Forkel, S. J., Pacella, V., Nozais, V., Allart, E., Piscicelli, C., … & Thiebaut de Schotten, M. (2023). Latent disconnectome prediction of long-term cognitive-behavioural symptoms in stroke. Brain, 146(5), 1963–1978.

Tettamanti, M., Paulesu, E., Scifo, P., Maravita, A., Fazio, F., Perani, D., & Marzi, C. A. (2002). Interhemispheric transmission of visuomotor information in humans: fMRI evidence. Journal of Neurophysiology, 88(2), 1051–1058.

Thiebaut de Schotten, M., Dell’Acqua, F., Ratiu, P., Leslie, A., Howells, H., Cabanis, E.,…Dronkers, N. (2015). From Phineas Gage and Monsieur Leborgne to HM: revisiting disconnection syndromes. Cerebral Cortex, 25(12), 4812–4827.

Thiebaut de Schotten, M., Dell’Acqua, F., Forkel, S., Simmons, A., Vergani, F., Murphy, D. G., & Catani, M. (2011). A lateralized brain network for visuo-spatial attention. Nature Precedings, 1–1.

Tomasch, J. (1954). Size, distribution, and number of fibres in the human corpus callosum. The Anatomical Record, 119(1), 119–135.

Tournier, J.-D., Mori, S., & Leemans, A. (2011). Diffusion tensor imaging and beyond. Magnetic resonance in medicine, 65(6), 1532.

Treble, A., Hasan, K. M., Iftikhar, A., Stuebing, K. K., Kramer, L. A., Cox Jr, C. S., … & Ewing-Cobbs, L. (2013). Working memory and corpus callosum microstructural integrity after pediatric traumatic brain injury: a diffusion tensor tractography study. Journal of neurotrauma, 30(19), 1609–1619.

Valenstein, E., Bowers, D., Verfaellie, M., Heilman, K. M., Day, A., & Watson, R. T. (1987). Retrosplenial amnesia. Brain, 110(6), 1631–1646.

van der Knaap, L. J., & van der Ham, I. J. (2011). How does the corpus callosum mediate interhemispheric transfer? A review. Behavioural brain research, 223(1), 211–221.

Van Essen, D. C., Ugurbil, K., Auerbach, E., Barch, D., Behrens, T. E., Bucholz, R., … & WU-Minn HCP Consortium. (2012). The Human Connectome Project: a data acquisition perspective. Neuroimage, 62(4), 2222–2231.

Veronelli, L., Bonandrini, R., Caporali, A., Licciardo, D., Corbo, M., & Luzzatti, C. (2024). Clinical and structural disconnectome evaluation in a case of optic aphasia. Brain Structure and Function, 229(7), 1641–1654.

Vigneau, M., Beaucousin, V., Hervé, P.-Y., Jobard, G., Petit, L., Crivello, F.,…Tzourio-Mazoyer, N. (2011). What is right-hemisphere contribution to phonological, lexico-semantic, and sentence processing?: Insights from a meta-analysis. Neuroimage, 54(1), 577–593.

Vingerhoets, G., Acke, F., Alderweireldt, A. S., Nys, J., Vandemaele, P., & Achten, E. (2012). Cerebral lateralization of praxis in right-and left-handedness: Same pattern, different strength. Human brain mapping, 33(4), 763–777.

Vingerhoets, G., & Stroobant, N. (1999). Lateralization of cerebral blood flow velocity changes during cognitive tasks: A simultaneous bilateral transcranial Doppler study. Stroke, 30(10), 2152–2158.

Vu, A. T., Auerbach, E., Lenglet, C., Moeller, S., Sotiropoulos, S. N., Jbabdi, S., … & Ugurbil, K. (2015). High resolution whole brain diffusion imaging at 7 T for the Human Connectome Project. Neuroimage, 122, 318–331.

Wakana, S., Caprihan, A., Panzenboeck, M. M., Fallon, J. H., Perry, M., Gollub, R. L.,…Dubey, P. (2007). Reproducibility of quantitative tractography methods applied to cerebral white matter. Neuroimage, 36(3), 630–644.

Wang, R., Benner, T., Sorensen, A. G., & Wedeen, V. J. (2007). Diffusion toolkit: a software package for diffusion imaging data processing and tractography. Proc Intl Soc Mag Reson Med,

Wang, P., Meng, C., Yuan, R., Wang, J., Yang, H., Zhang, T., … & Biswal, B. B. (2020). The organization of the human corpus callosum estimated by intrinsic functional connectivity with white-matter functional networks. Cerebral Cortex, 30(5), 3313–3324.

Wang, P., Wang, J., Tang, Q., Alvarez, T. L., Wang, Z., Kung, Y. C., … & Biswal, B. B. (2021). Structural and functional connectivity mapping of the human corpus callosum organization with white-matter functional networks. Neuroimage, 227, 117642.

Xiong, Y., Yang, L., Wang, C., Zhao, C., Luo, J., Wu, D., … & Gong, G. (2024). Cortical mapping of callosal connections in healthy young adults. Human Brain Mapping, 45(3), e26629.

Yarkoni, T., Poldrack, R. A., Nichols, T. E., Van Essen, D. C., & Wager, T. D. (2011). Large-scale automated synthesis of human functional neuroimaging data. Nat Methods, 8(8), 665–670. 10.1038/nmeth.1635

Yasuda, Y., Watanabe, T., Tanaka, H., Tadashi, I., & Akiguchi, I. (1997). Amnesia following infarction in the right retrosplenial region. Clinical neurology and neurosurgery, 99(2), 102–105.

Zaidel, E. (1973). Linguistic competence and related functions in the right cerebral hemisphere of man following commissurotomy an hemisphererectomy California Institute of Technology].

Zaidel, E. (1982). Reading by the disconnected right hemisphere: An aphasiological perspective. In Dyslexia: Neuronal, Cognitive & Linguistic Aspects (pp. 67-91). Elsevier.

Zaidel, E. (1983). A response to Gazzaniga: Language in the right hemisphere, convergent perspectives.

Zaidel, E., Zaidel, D. W., & Bogen, J. E. (1999). The split brain. Encyclopedia of neuroscience. Amsterdam: Elsevier Science.

