## Supplementary Material for "Functional projection of cognitive functions through the human corpus callosum"

Medial Sagittal Callosal Atlas

***Correspondence:**
Rolando Bonandrini

Frontal Medial Cortex……………………………………………………………….. Page 9
Frontal Pole…………………………………………………………………………... Page 10
Cingulate Gyrus, anterior division………………………………………………….. Page 11
Paracingulate Gyrus…………………………………………………………………. Page 12
Central Opercular Cortex……………………………………………………………. Page 17
Postcentral Gyrus……………………………………………………………………. Page 18
Superior Parietal Lobule…………………………………………………………….. Page 19
Angular Gyrus…………………………………………………………………………. Page 20
Supramarginal Gyrus, anterior division……………………………………………. Page 21
Parietal Operculum Cortex…………………………………………………………... Page 22
Supramarginal Gyrus, posterior division…………………………………………... Page 23
Cingulate Gyrus, posterior division………………………………………………... Page 24
Precuneus…………………………………………………………………………….. Page 25
Occipital Fusiform Gyrus……………………………………………………………. Page 32
Lateral Occipital Cortex, inferior division…………………………………………. Page 33
Temporal Occipital Fusiform Cortex……………………………………………….. Page 34
Inferior Temporal Gyrus, temporooccipital part…………………………………… Page 35
Middle Temporal Gyrus, temporooccipital part……………………………………. Page 36
Inferior Temporal Gyrus, posterior division……………………………………….. Page 37
Temporal Fusiform Cortex, posterior division……………………………………... Page 38
Parahippocampal Gyrus, posterior division……………………………………….. Page 39
Temporal Pole………………………………………………………………………... Page 50

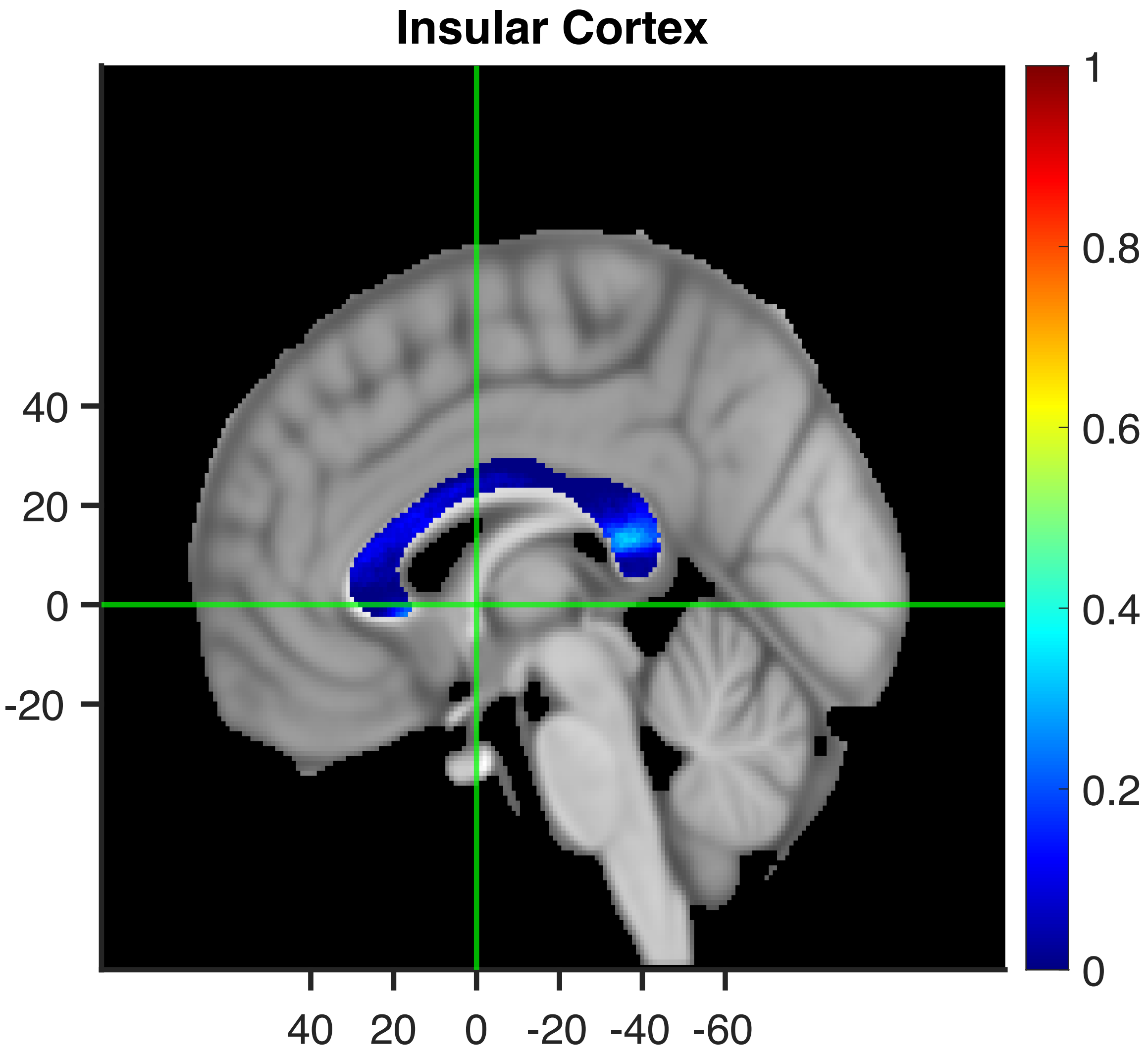
Insular Cortex
40
20
0
-20
40 20 0 -20 -40 -60
1
0.8
0.6
0.4
0.2
0

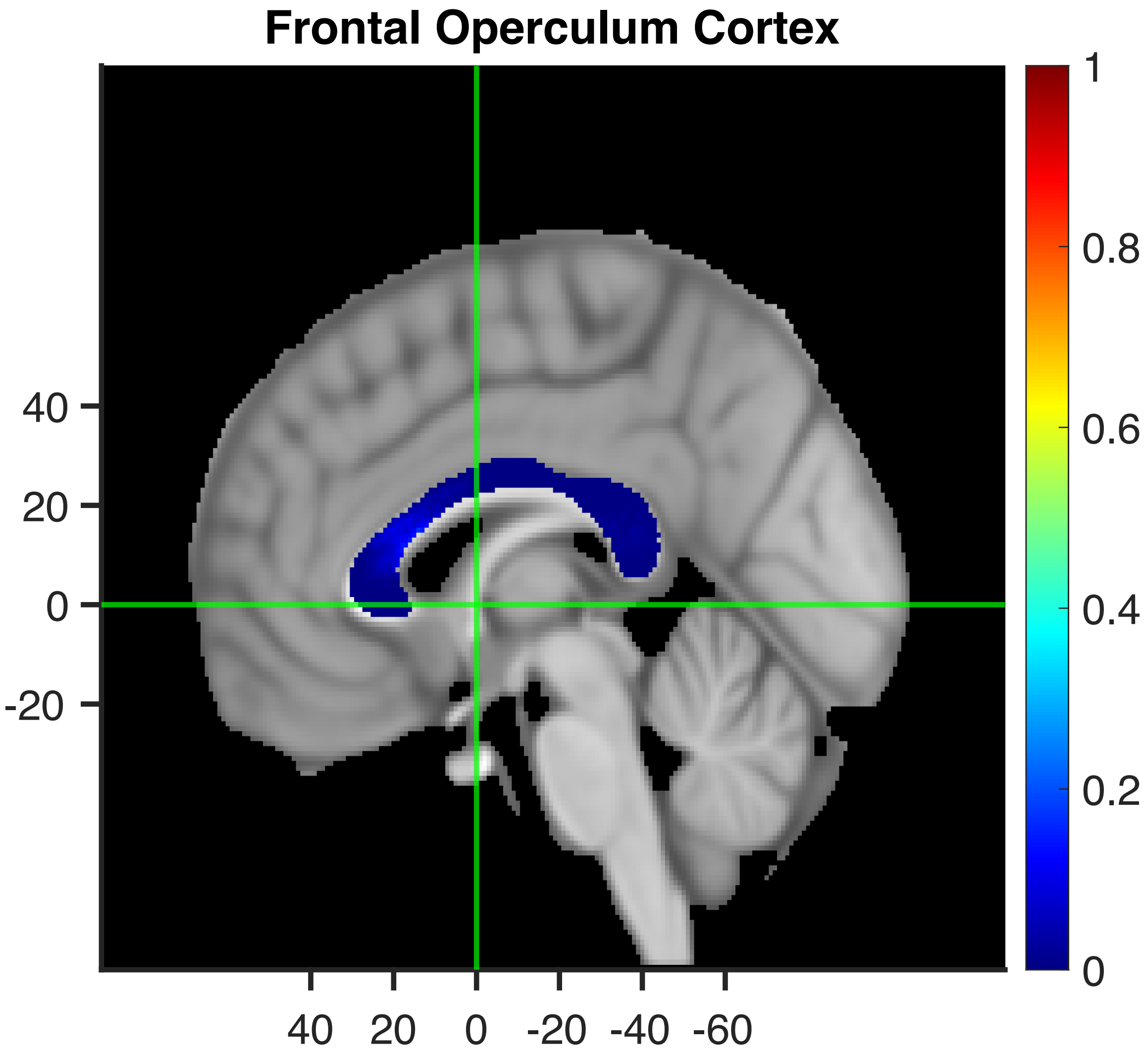
Frontal Operculum Cortex
40
20
0
-20
40 20 0 -20 -40 -60
1
0.8
0.6
0.4
0.2
0

Inferior Frontal Gyrus, pars triangularis

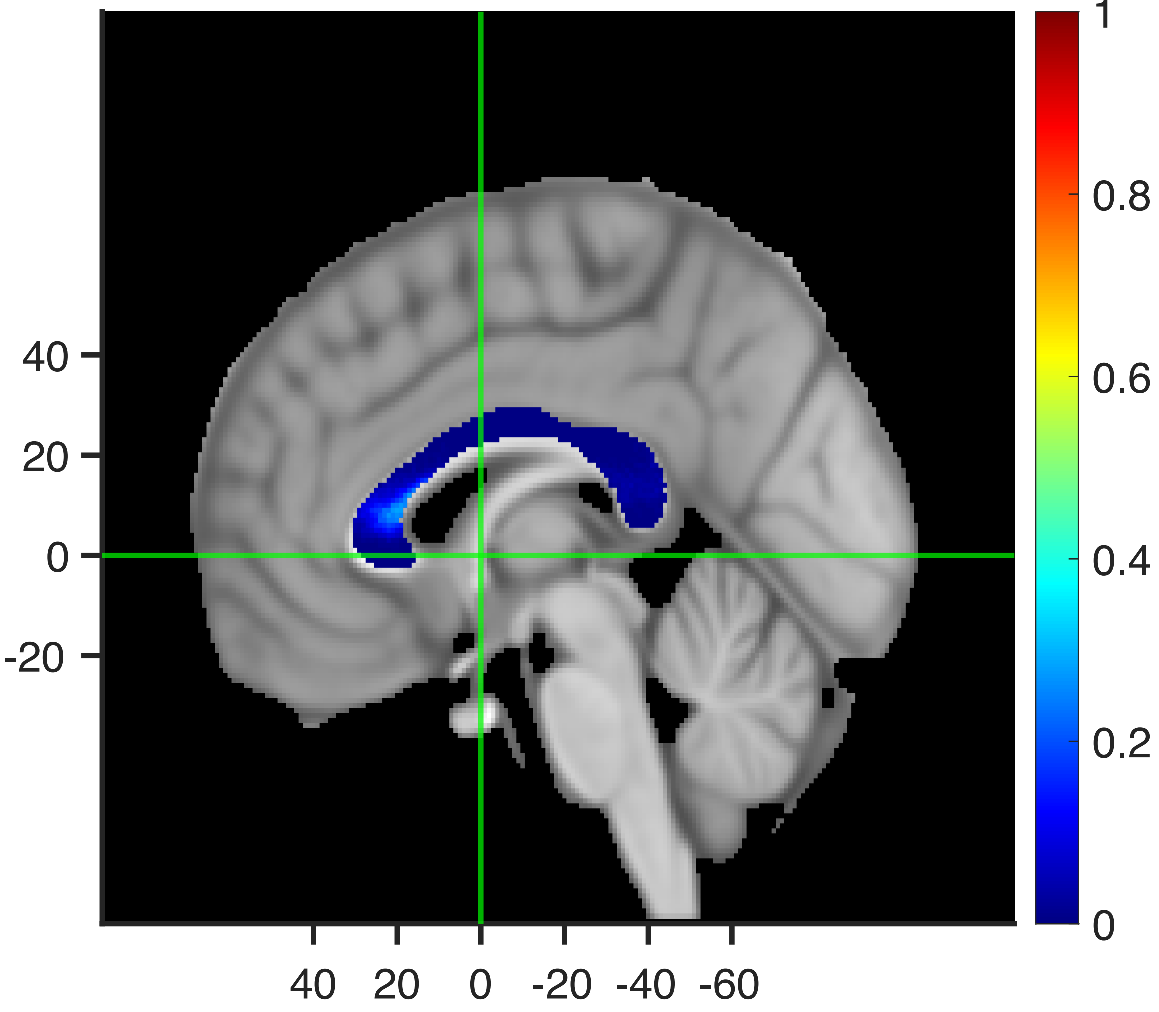

Inferior Frontal Gyrus, pars opercularis

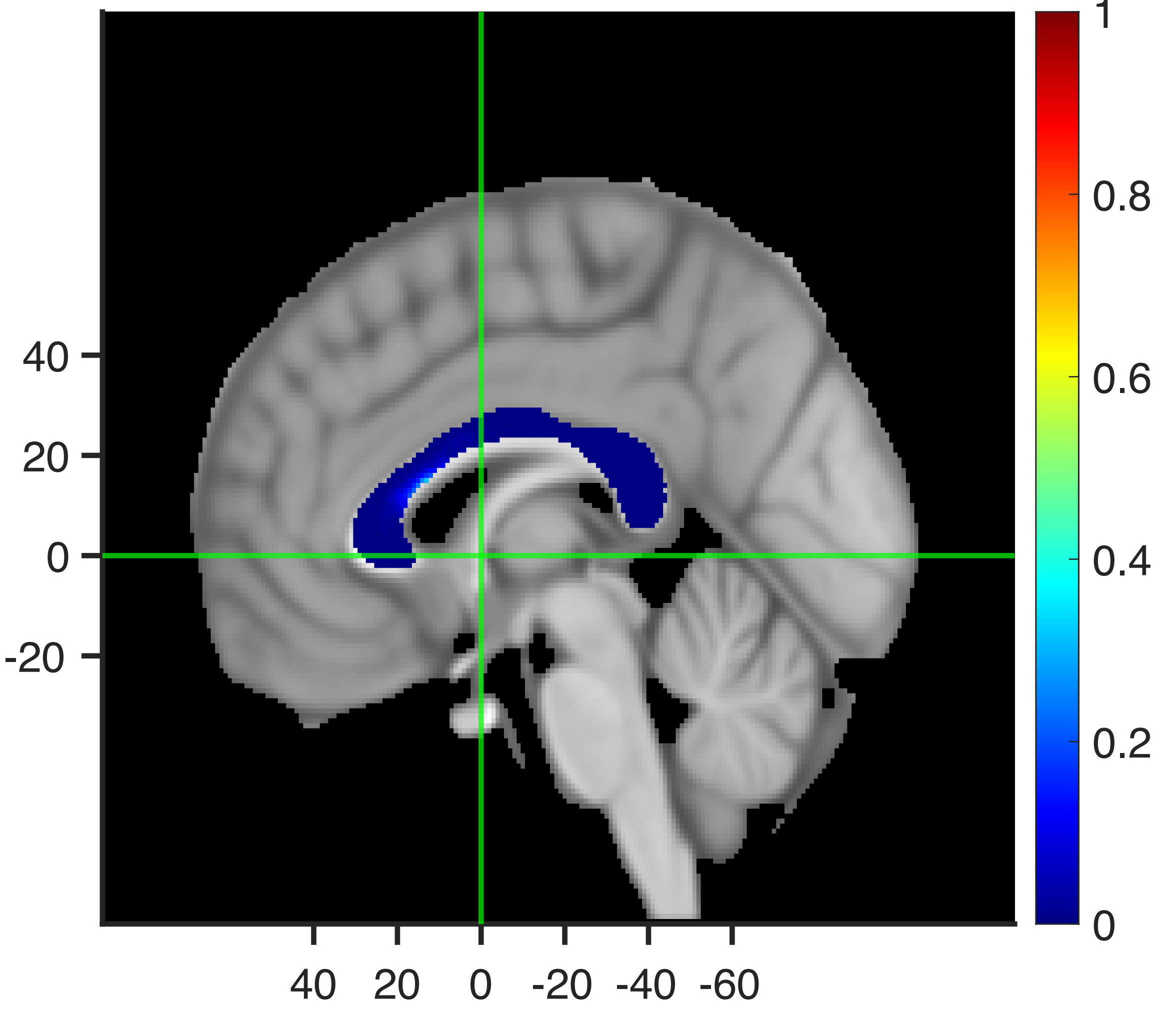

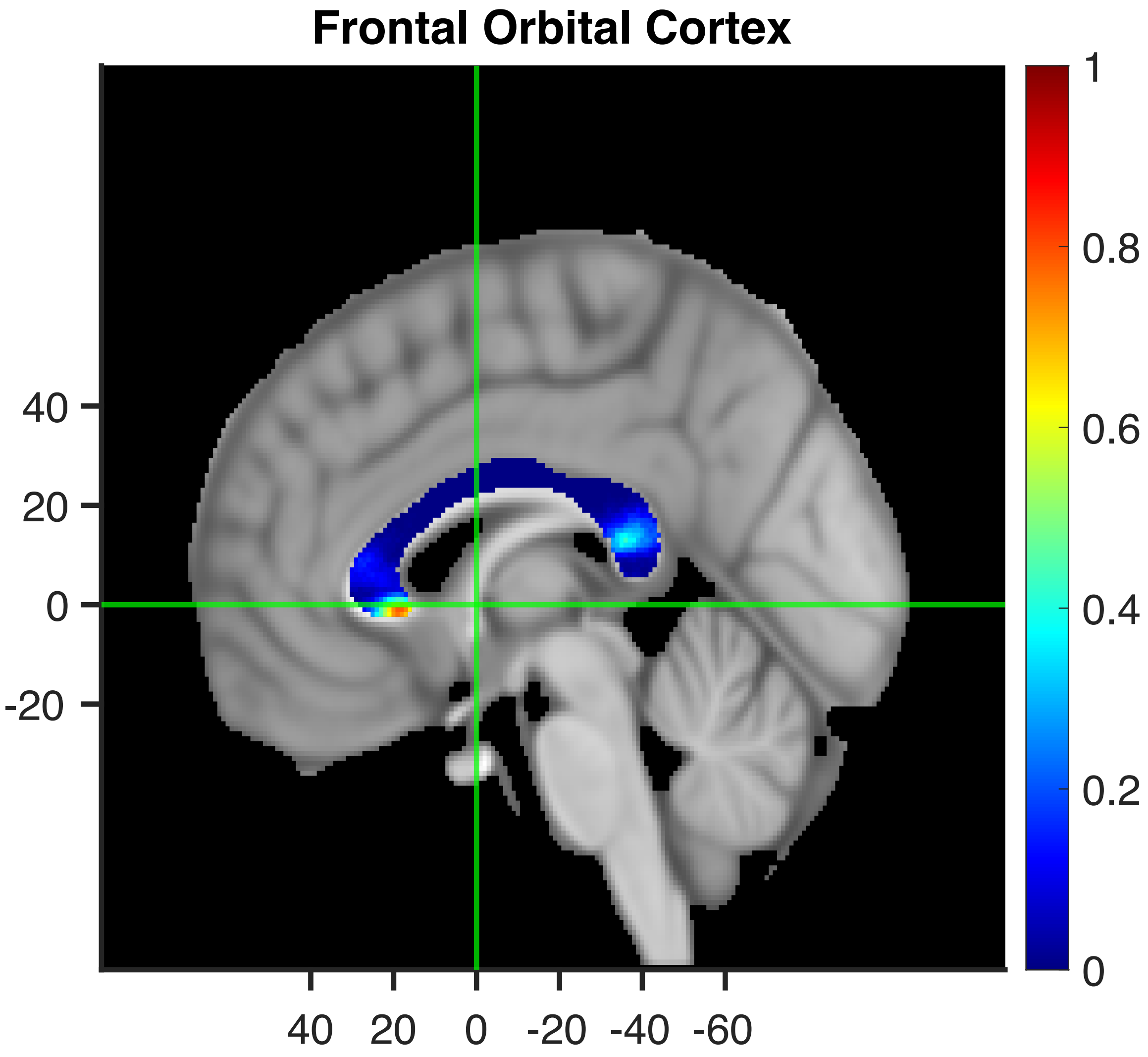
Frontal Orbital Cortex
40
20
0
-20
40 20 0 -20 -40 -60
1
0.8
0.6
0.4
0.2
0

#### Subcallosal Cortex

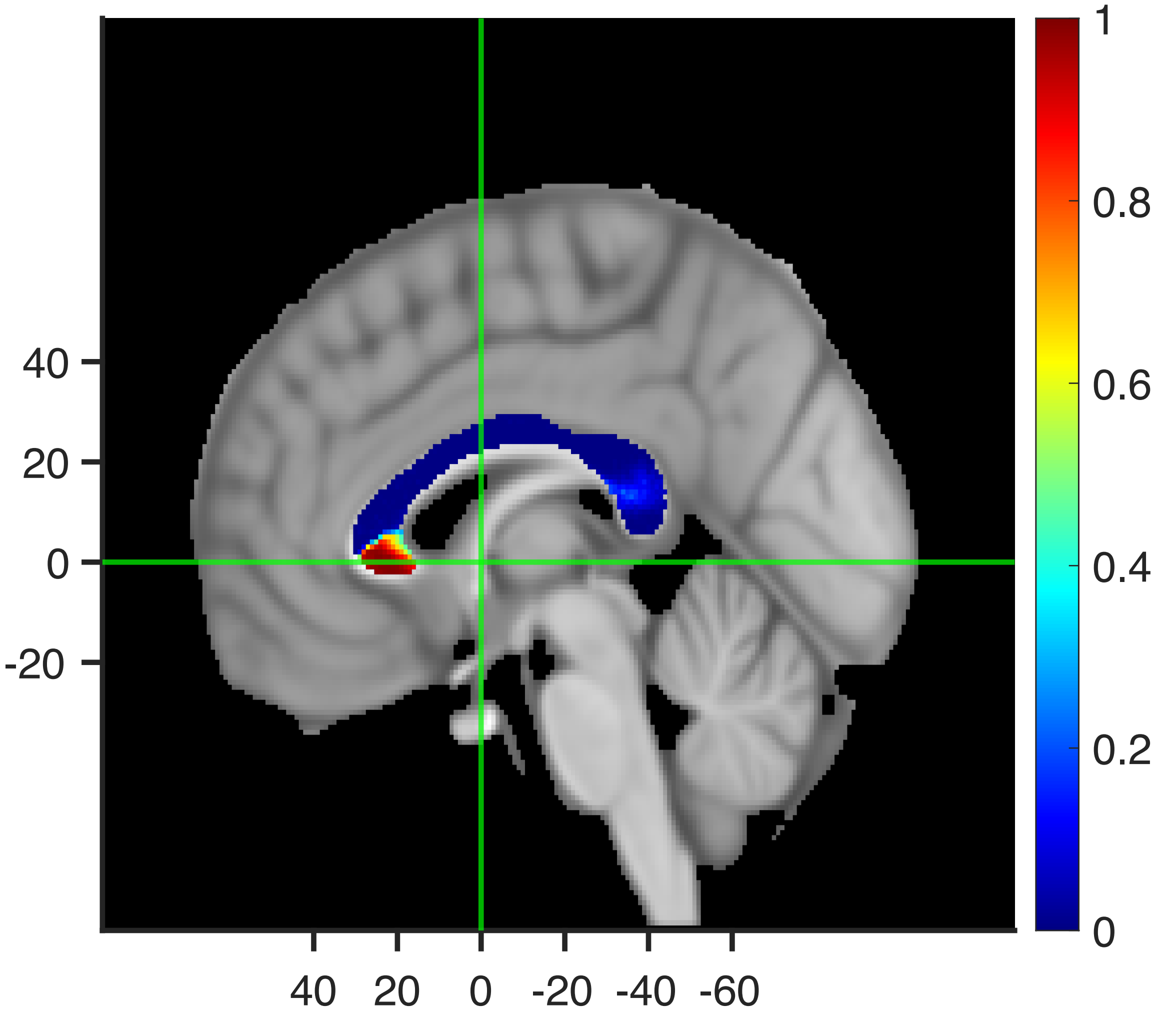

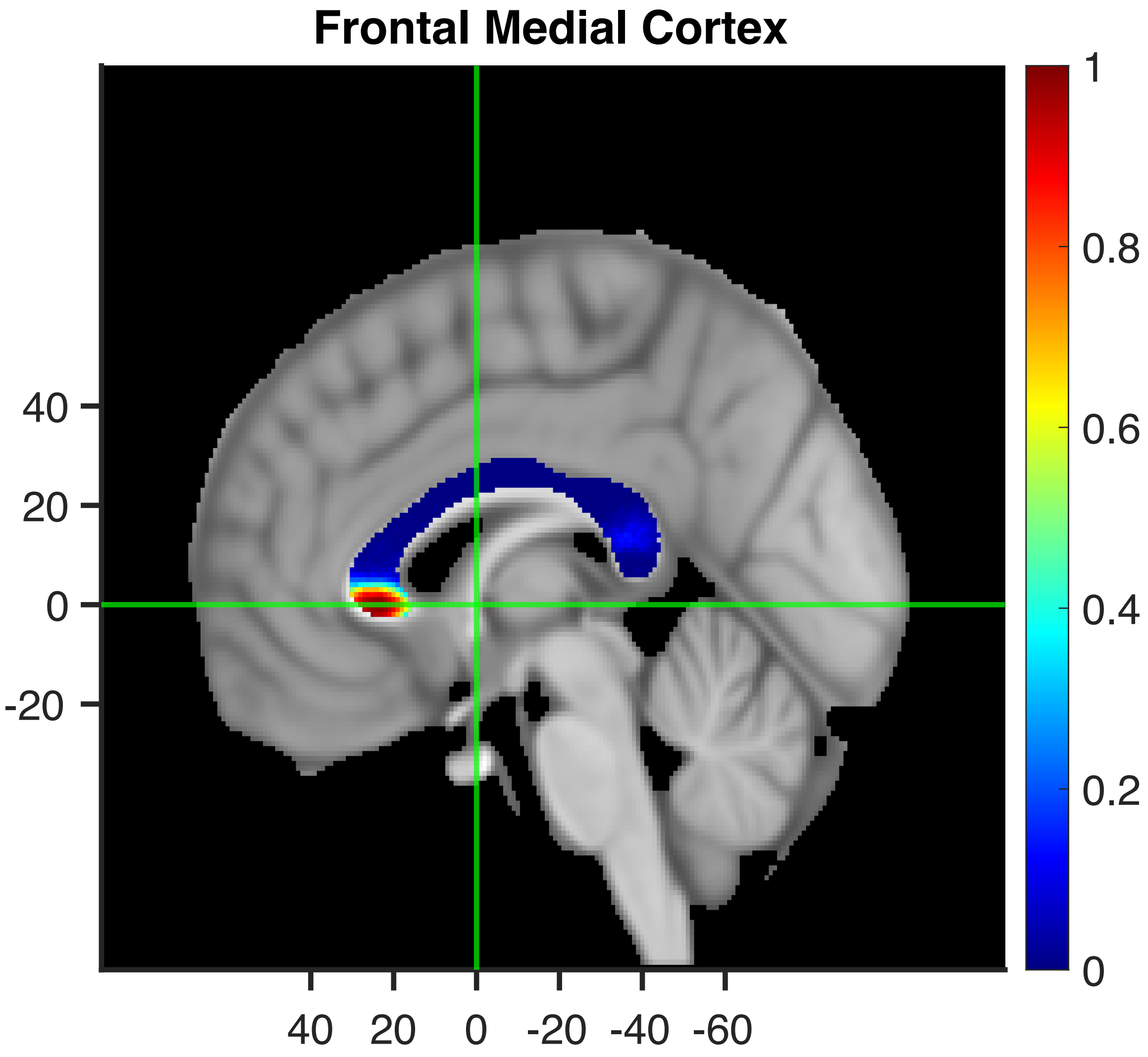

Frontal Medial Cortex
40
20
0
-20
40
20
0
-20
-40
-60
1
0.8
0.6
0.4
0.2
0

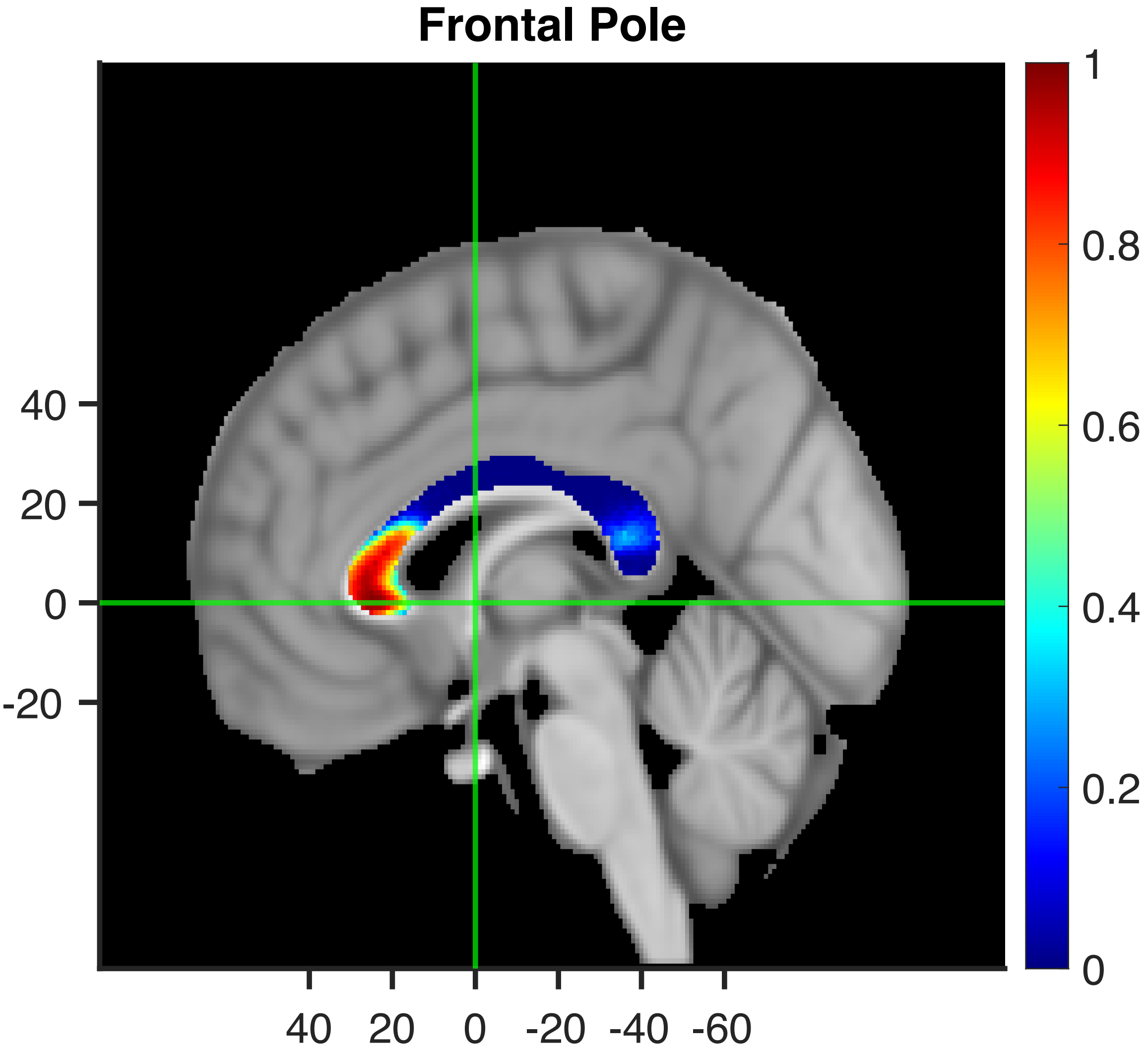
Frontal Pole
40
20
0
-20
40 20 0 -20 -40 -60
1
0.8
0.6
0.4
0.2
0

#### Cingulate Gyrus, anterior division

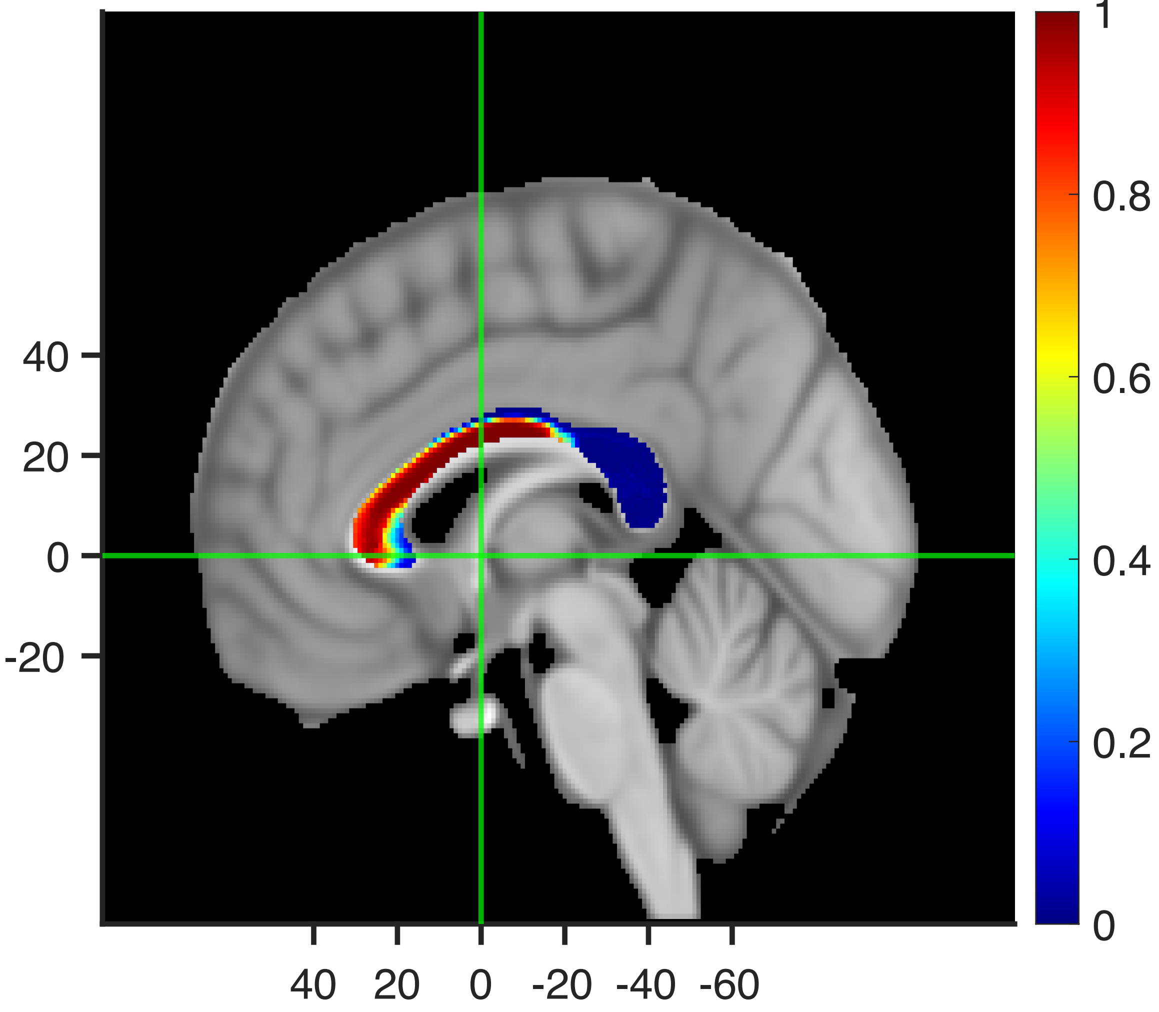

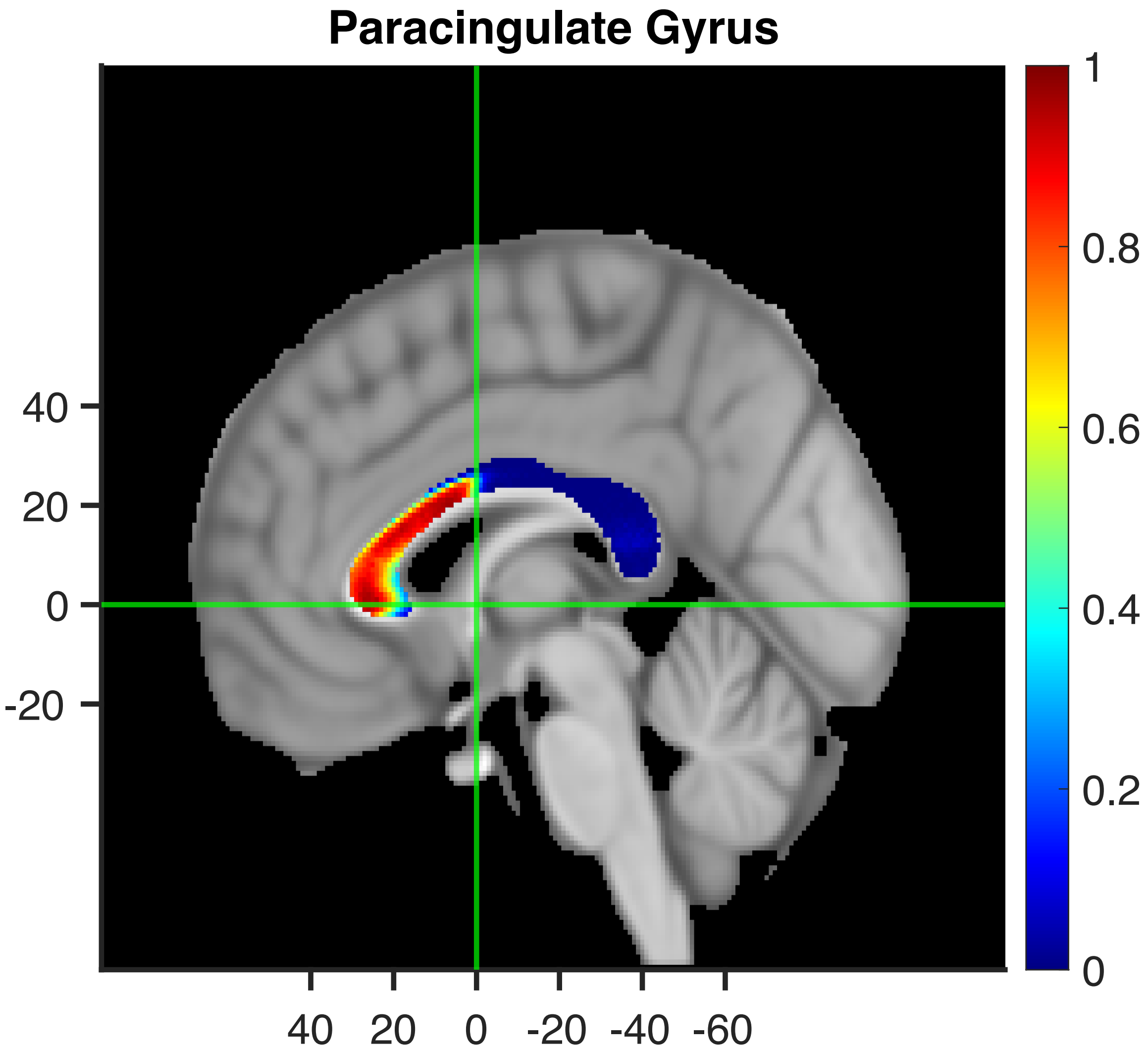
Paracingulate Gyrus
40
20
0
-20
40 20 0 -20 -40 -60
1
0.8
0.6
0.4
0.2
0

#### Middle Frontal Gyrus

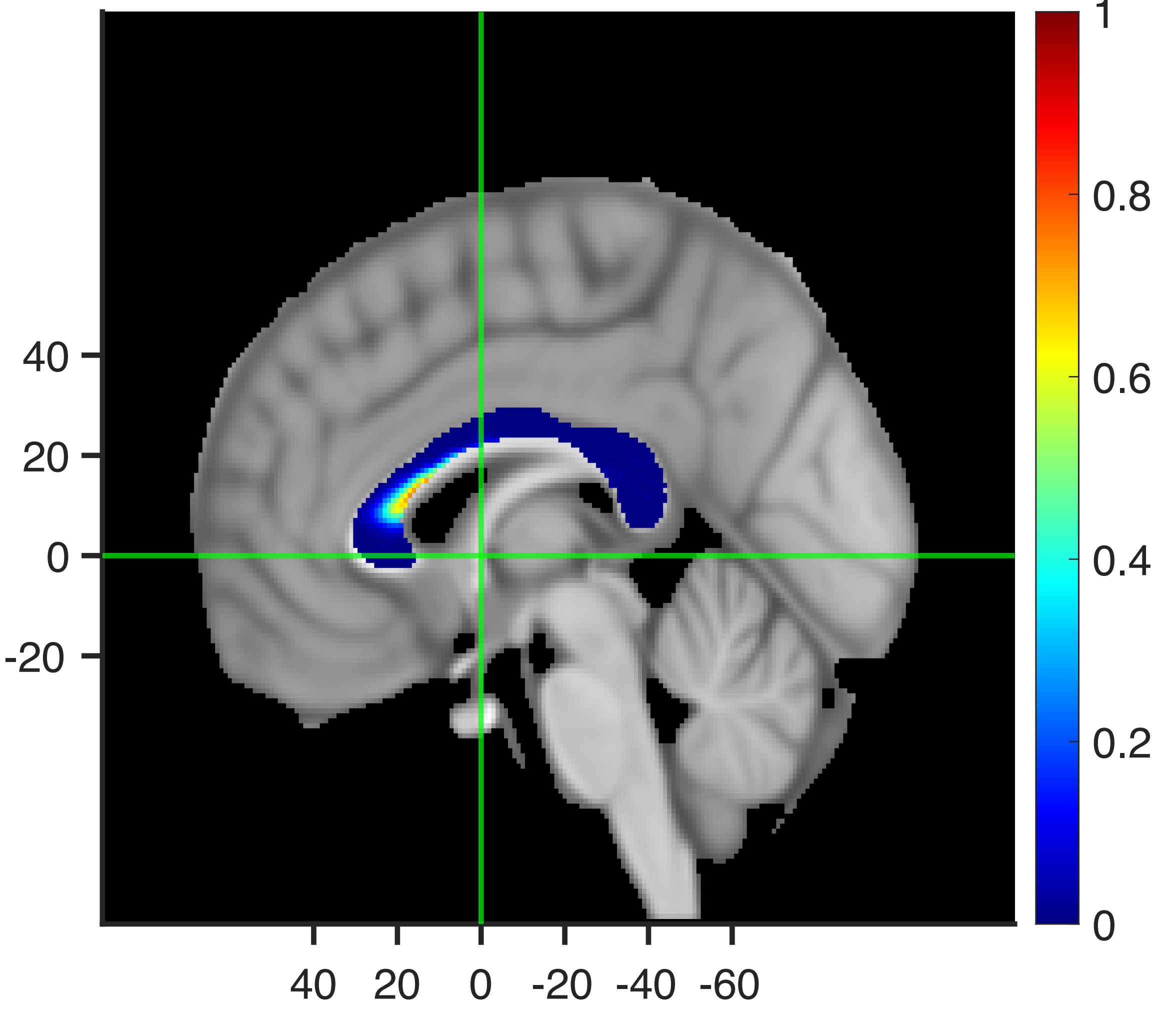

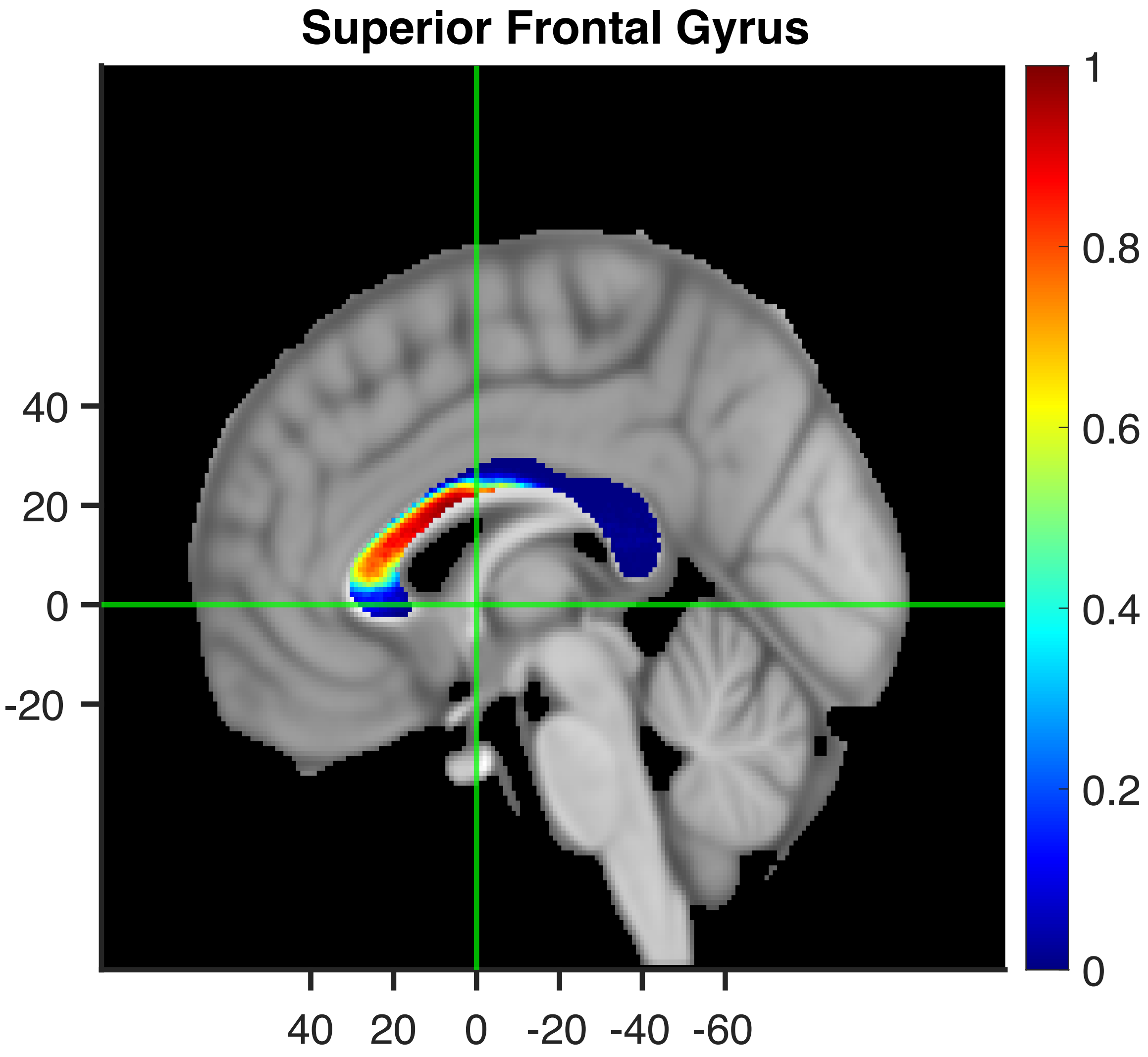

Superior Frontal Gyrus
40
20
0
-20
40
20
0
-20
-40
-60
1
0.8
0.6
0.4
0.2
0

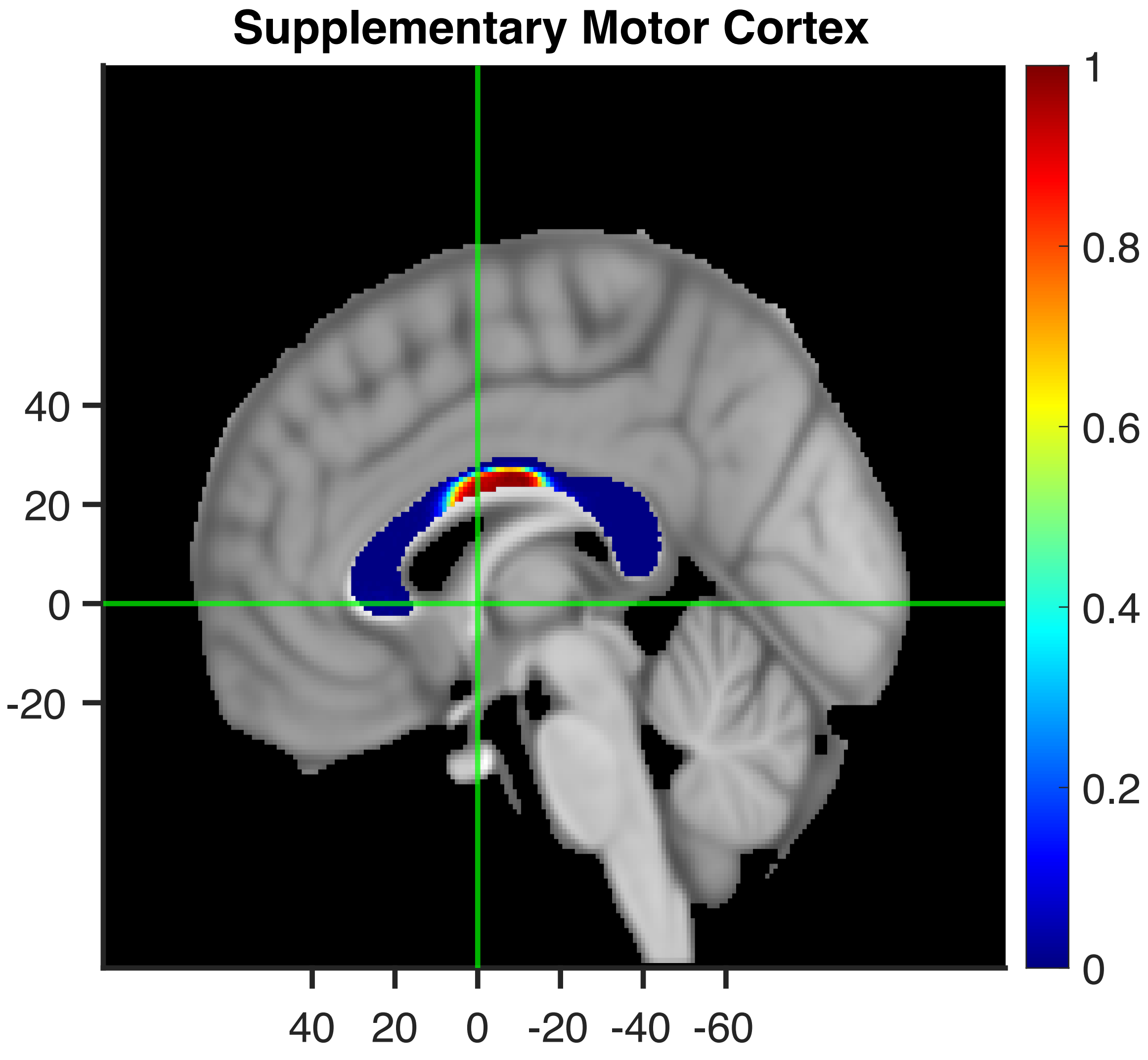
Supplementary Motor Cortex
40
20
0
-20
40
20
0
-20
-40
-60
1
0.8
0.6
0.4
0.2
0

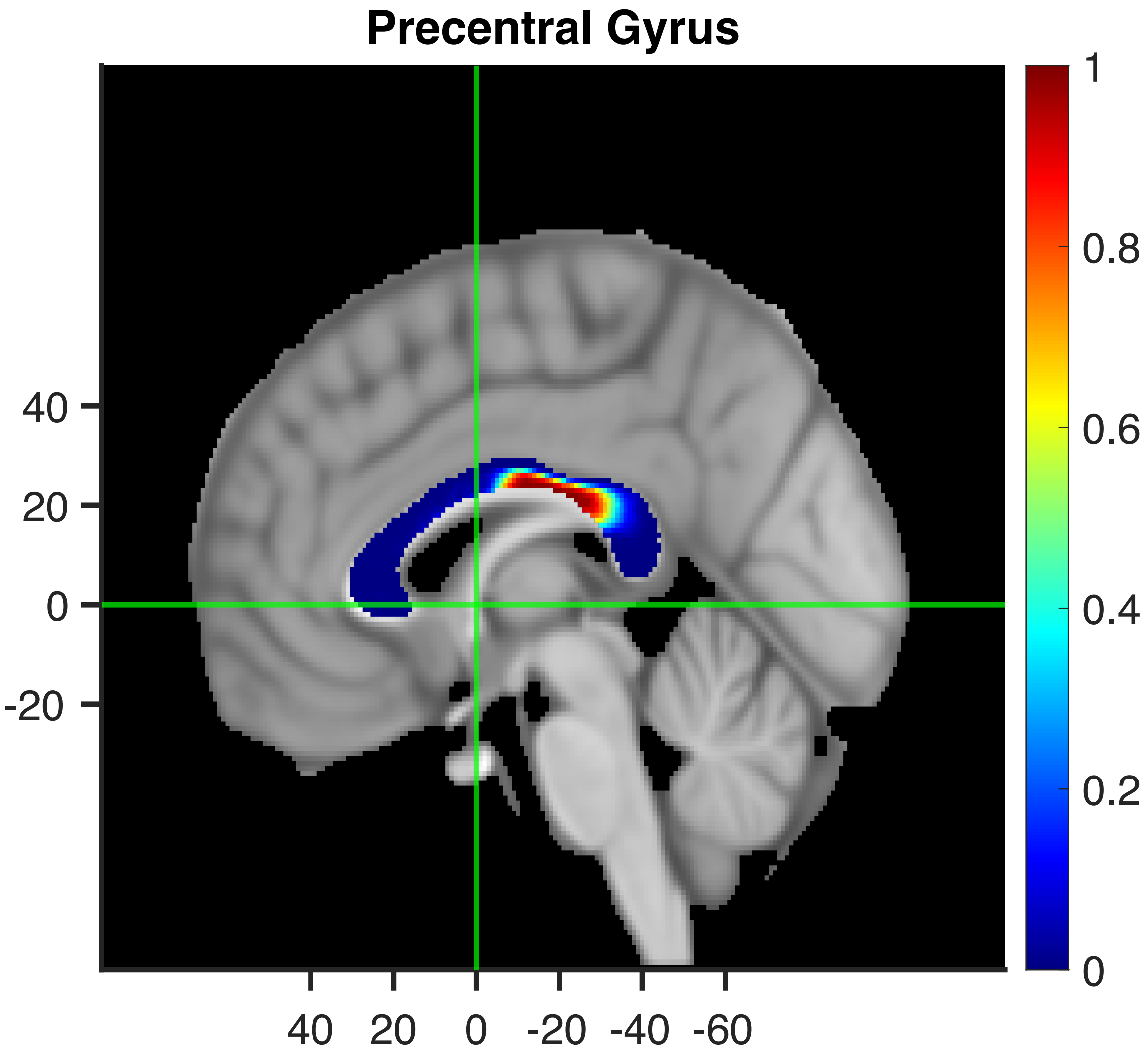

Precentral Gyrus
40
20
0
-20
40
20
0
-20
-40
-60
1
0.8
0.6
0.4
0.2
0

### Central Opercular Cortex

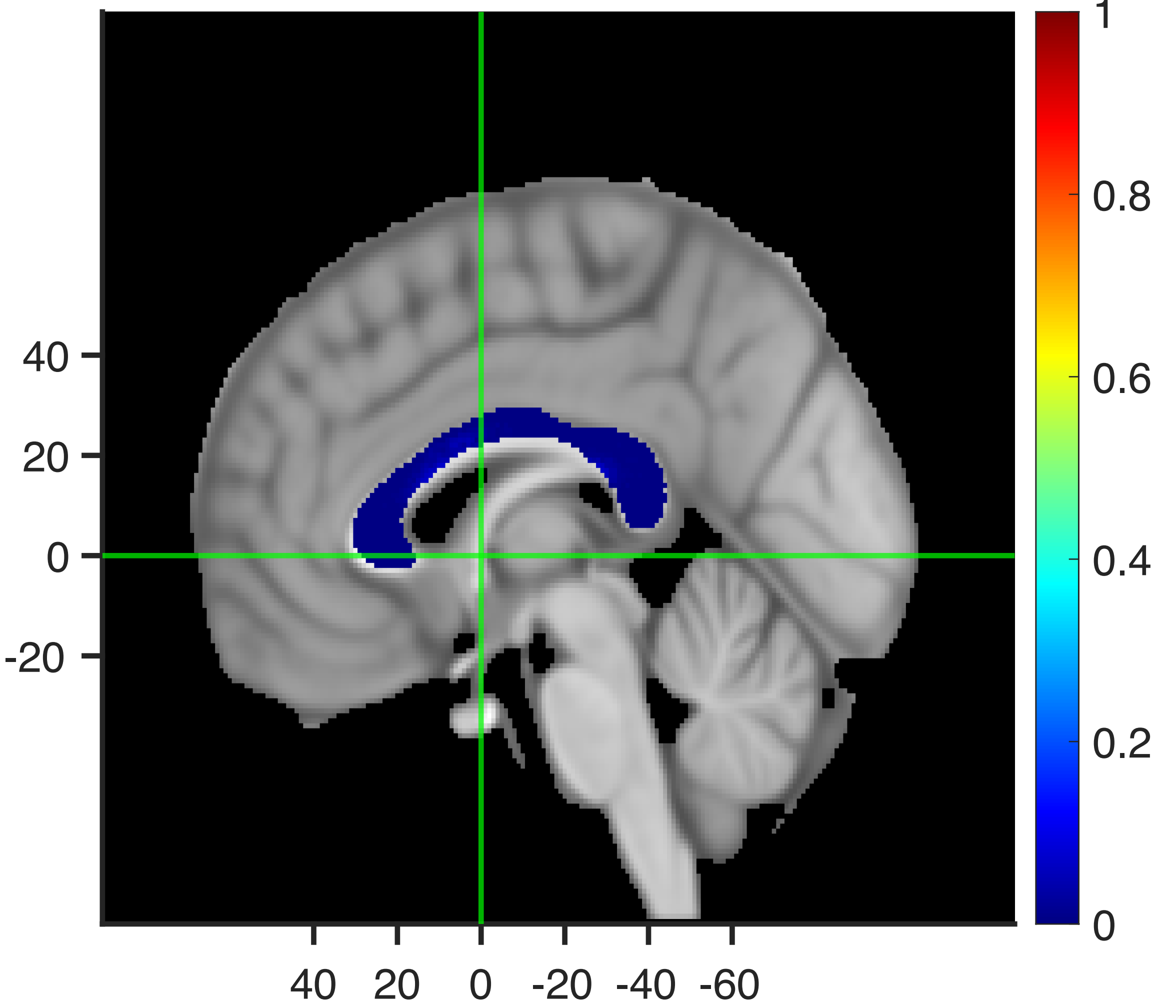

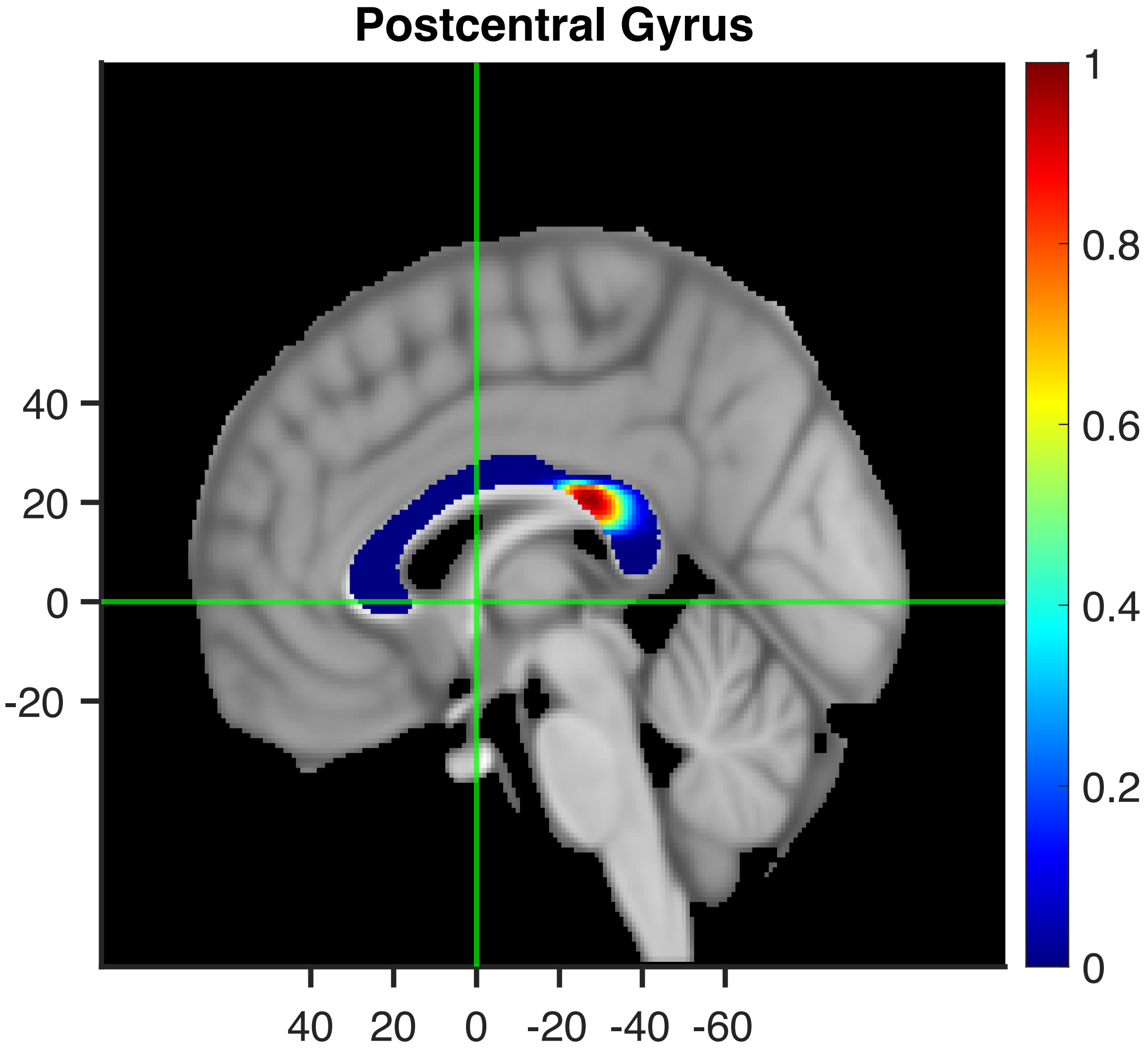
Postcentral Gyrus
40
20
0
-20
40
20
0
-20
-40
-60
1
0.8
0.6
0.4
0.2
0

#### Superior Parietal Lobule

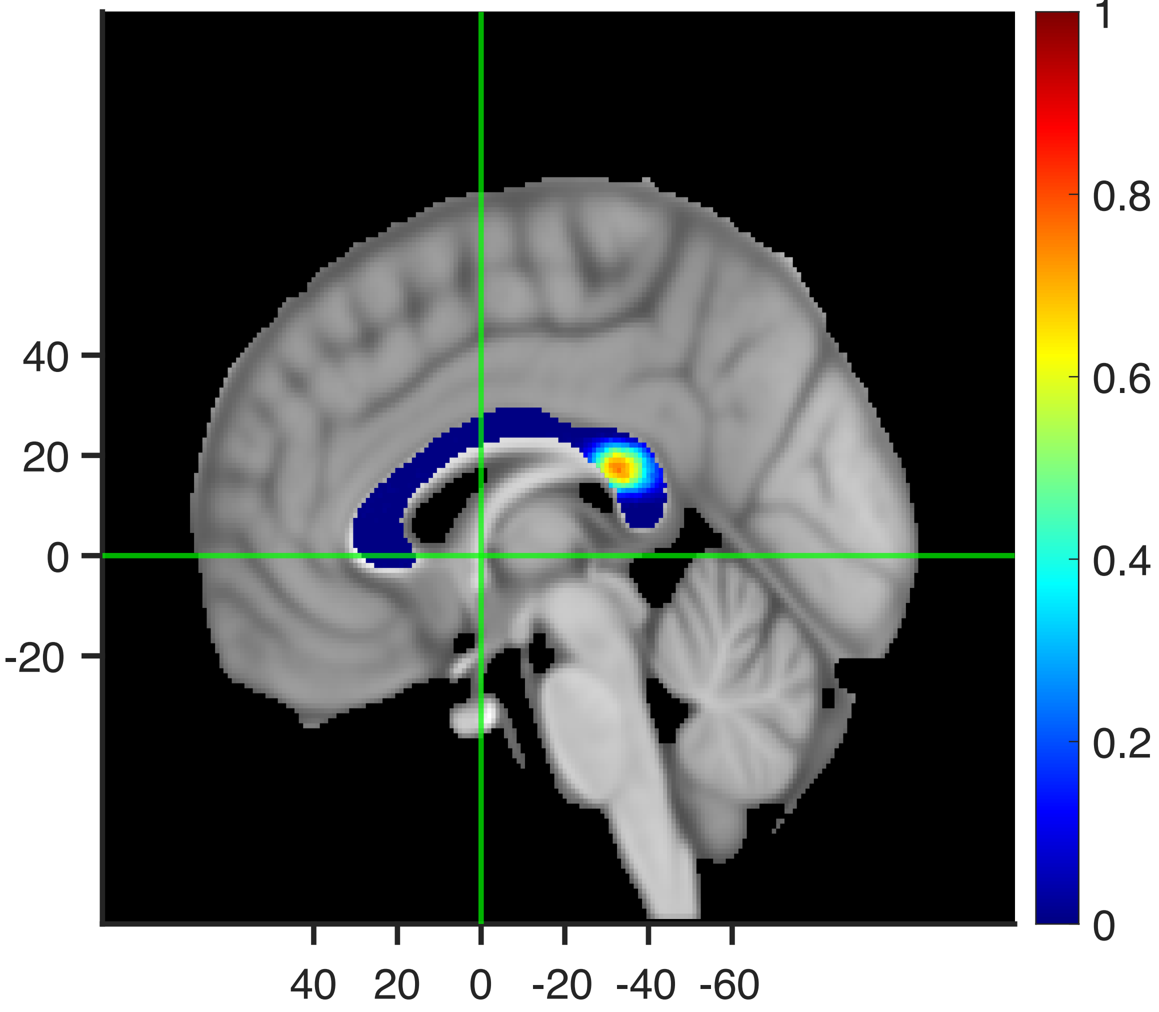

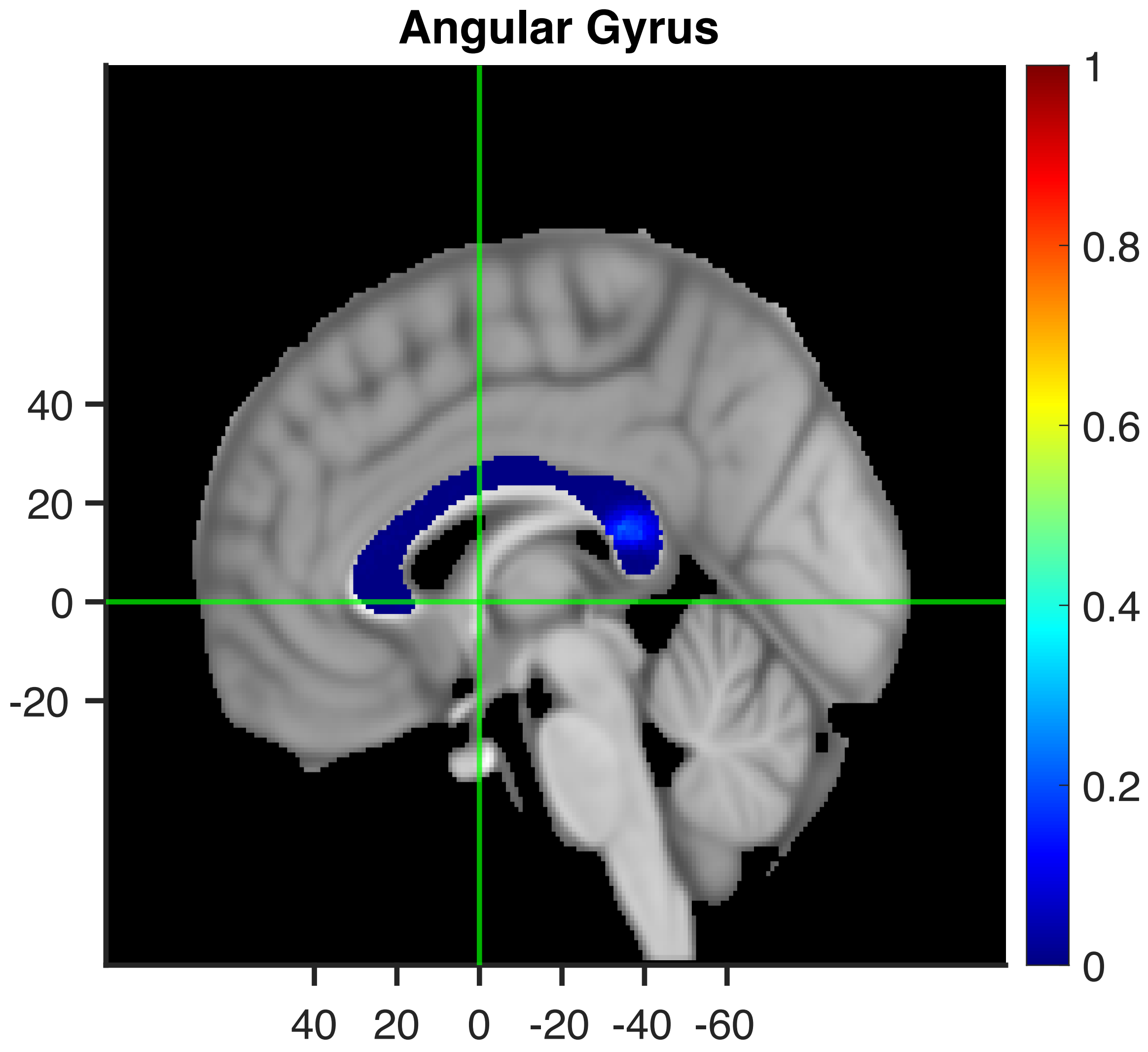

Angular Gyrus
40
20
0
-20
40 20 0 -20 -40 -60
1
0.8
0.6
0.4
0.2
0

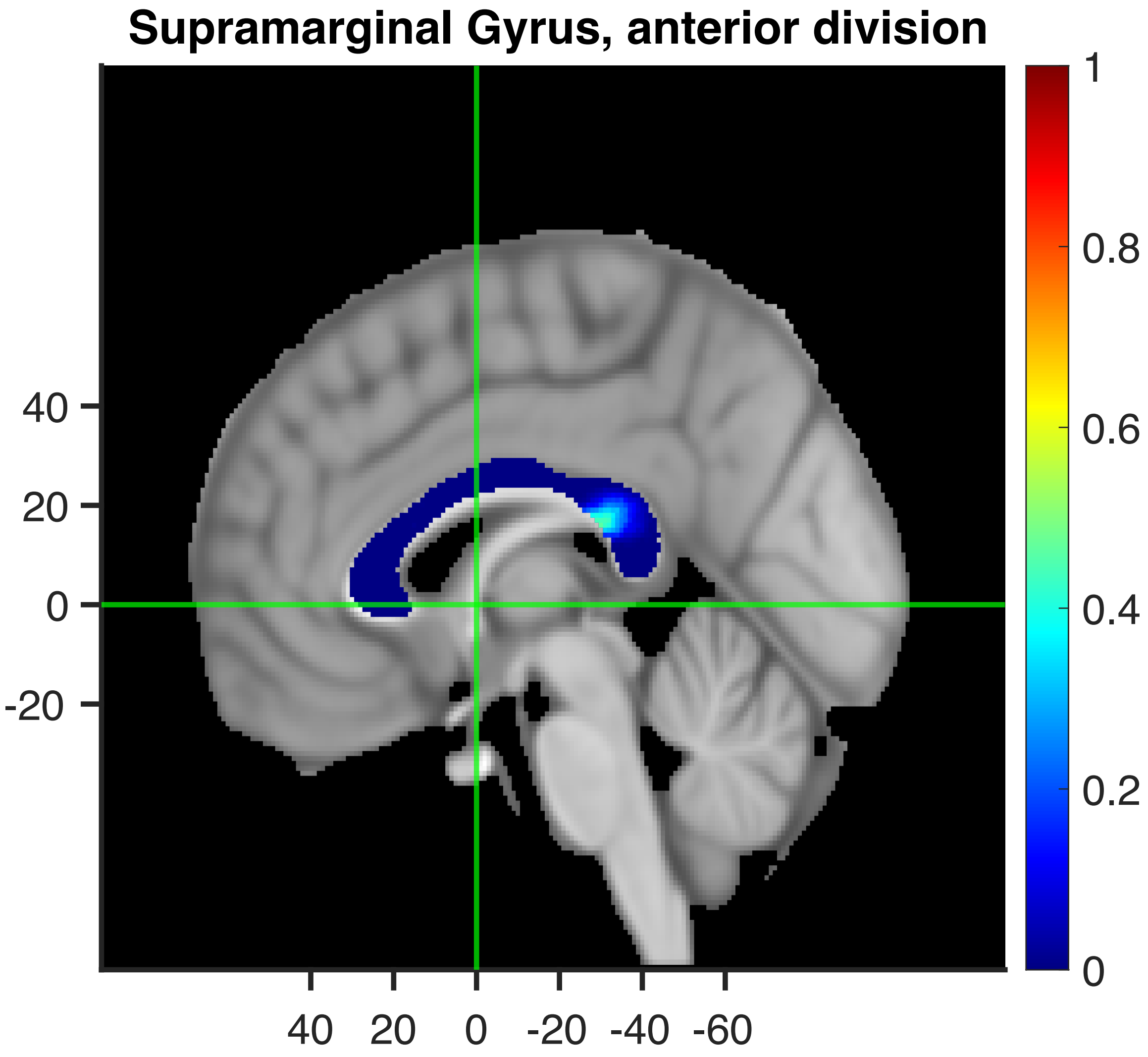
Supramarginal Gyrus, anterior division
40
20
0
-20
40
20
0
-20
-40
-60
1
0.8
0.6
0.4
0.2
0

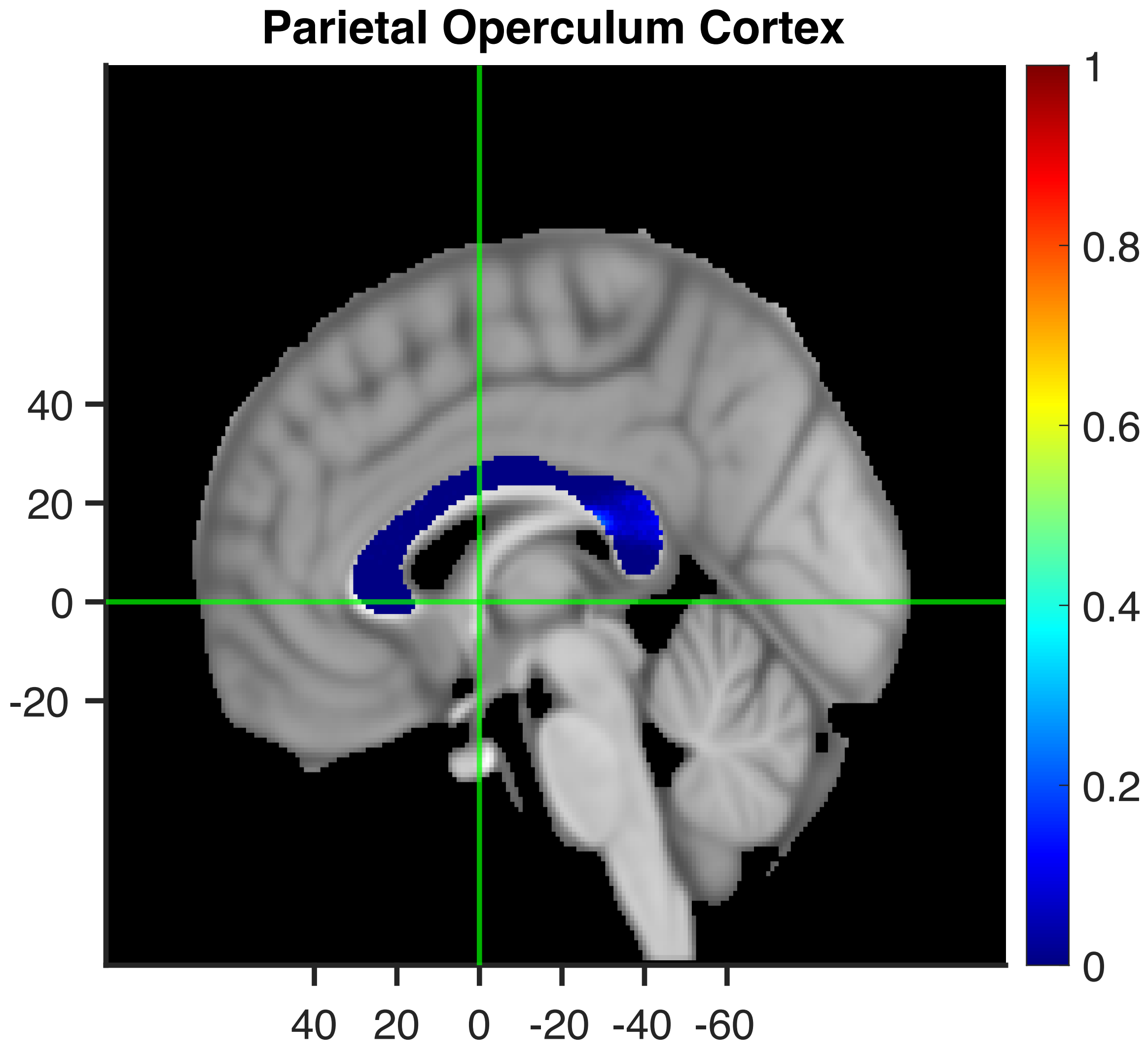

Parietal Operculum Cortex
40
20
0
-20
40 20 0 -20 -40 -60
1
0.8
0.6
0.4
0.2
0

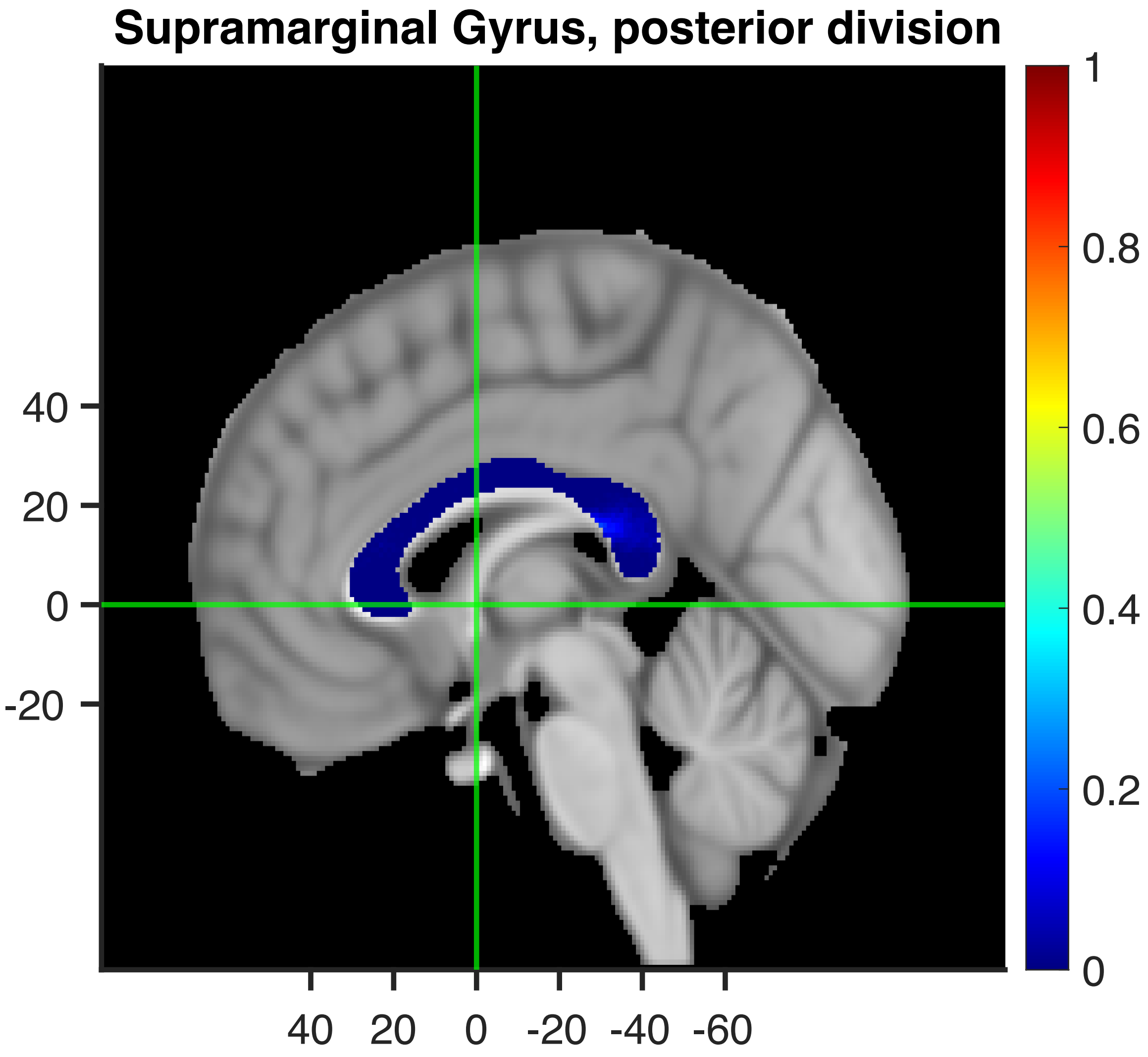
Supramarginal Gyrus, posterior division
40
20
0
-20
40
20
0
-20
-40
-60
1
0.8
0.6
0.4
0.2
0

**Cingulate Gyrus, posterior division**

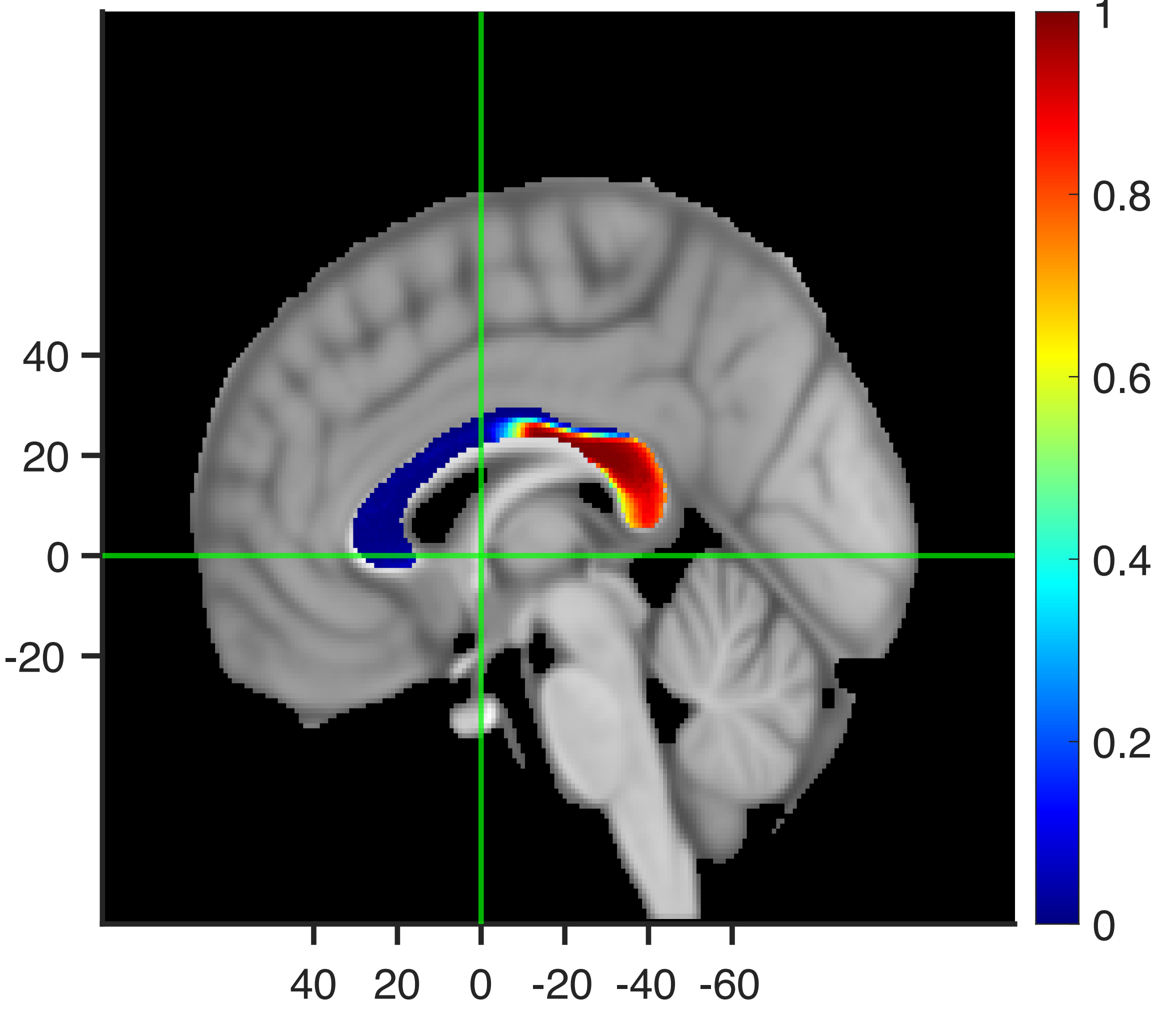

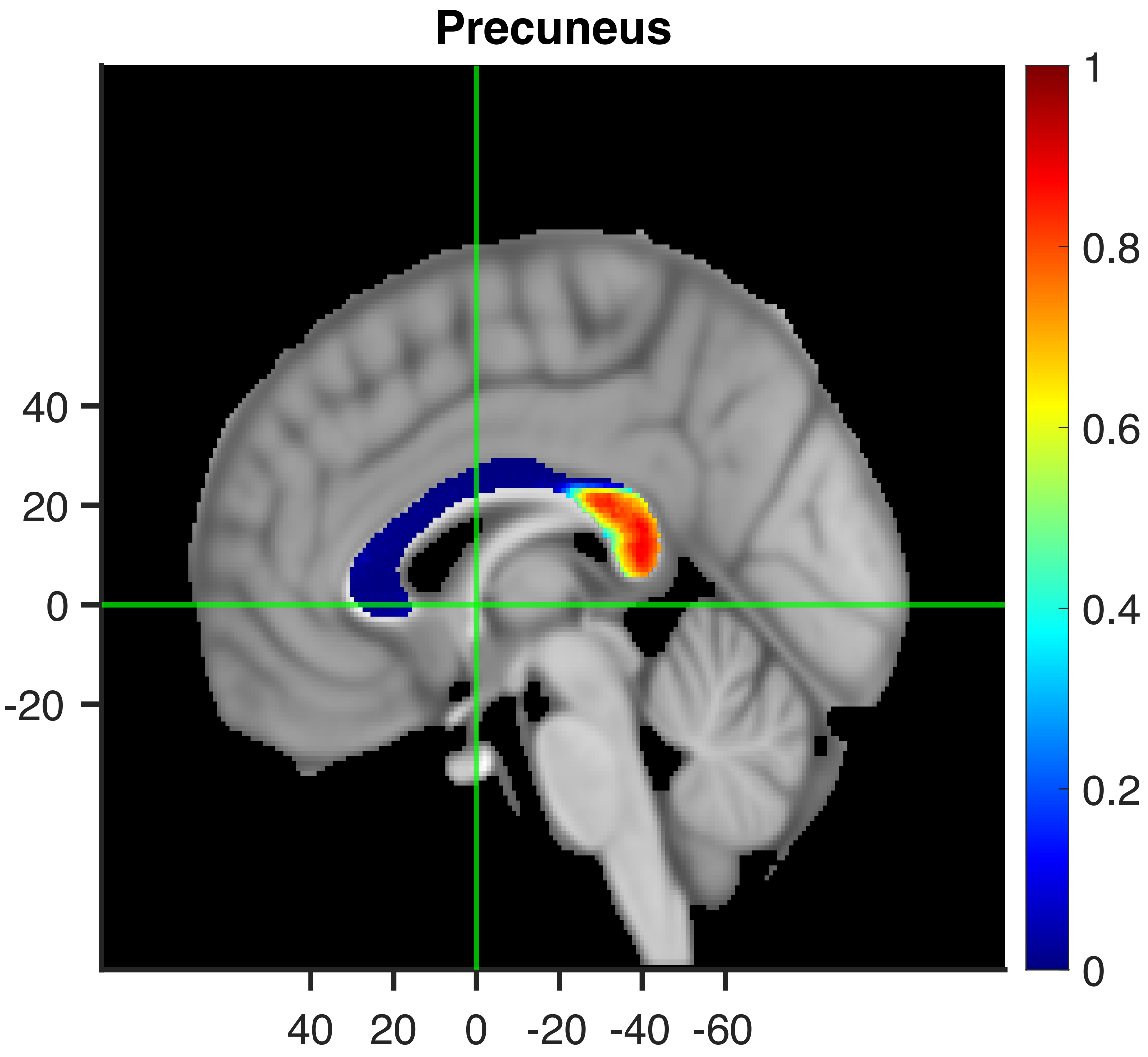
Precuneus
40
20
0
-20
40 20 0 -20 -40 -60
1
0.8
0.6
0.4
0.2
0

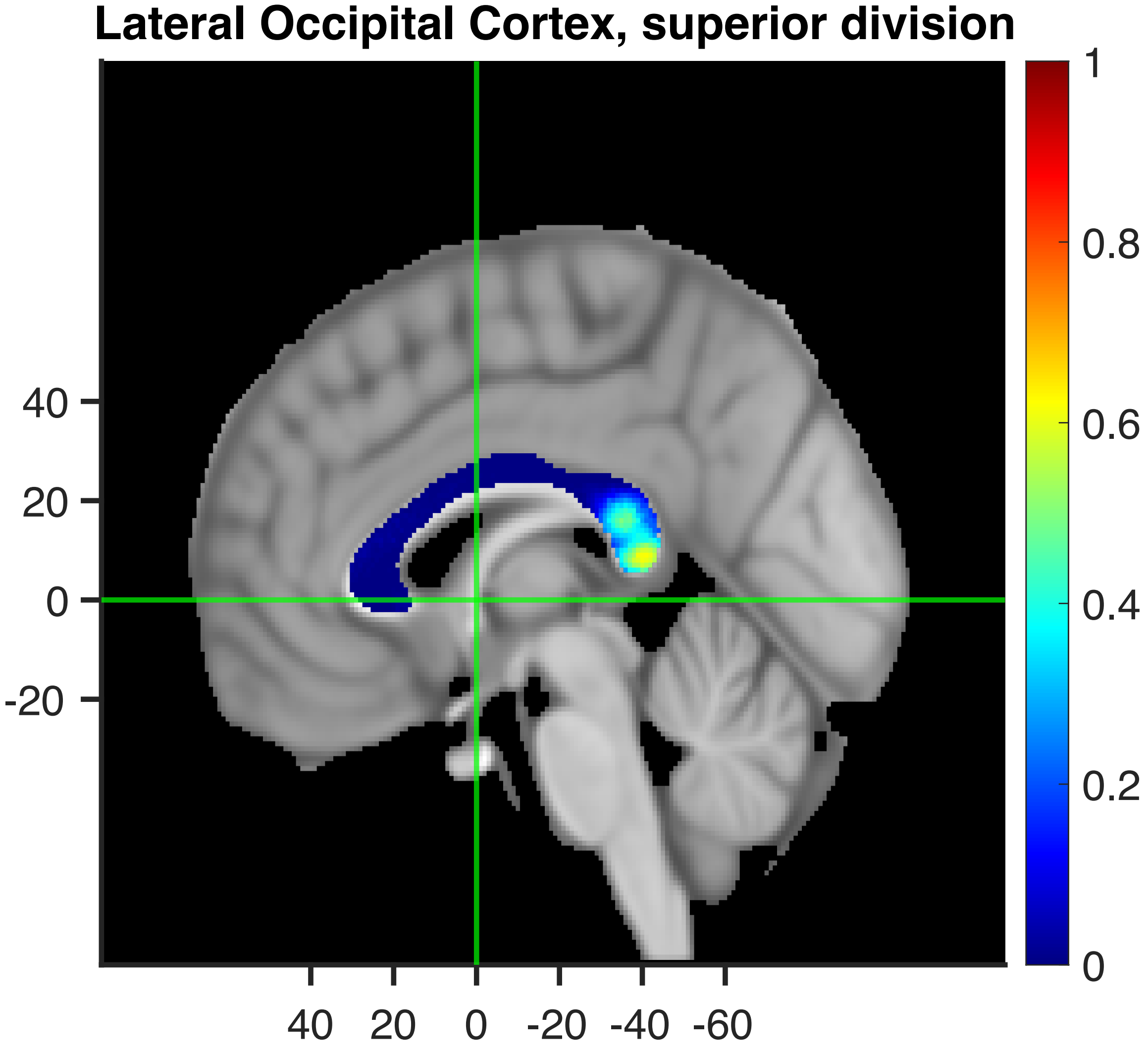
Lateral Occipital Cortex, superior division
40
20
0
-20
40 20 0 -20 -40 -60
1
0.8
0.6
0.4
0.2
0

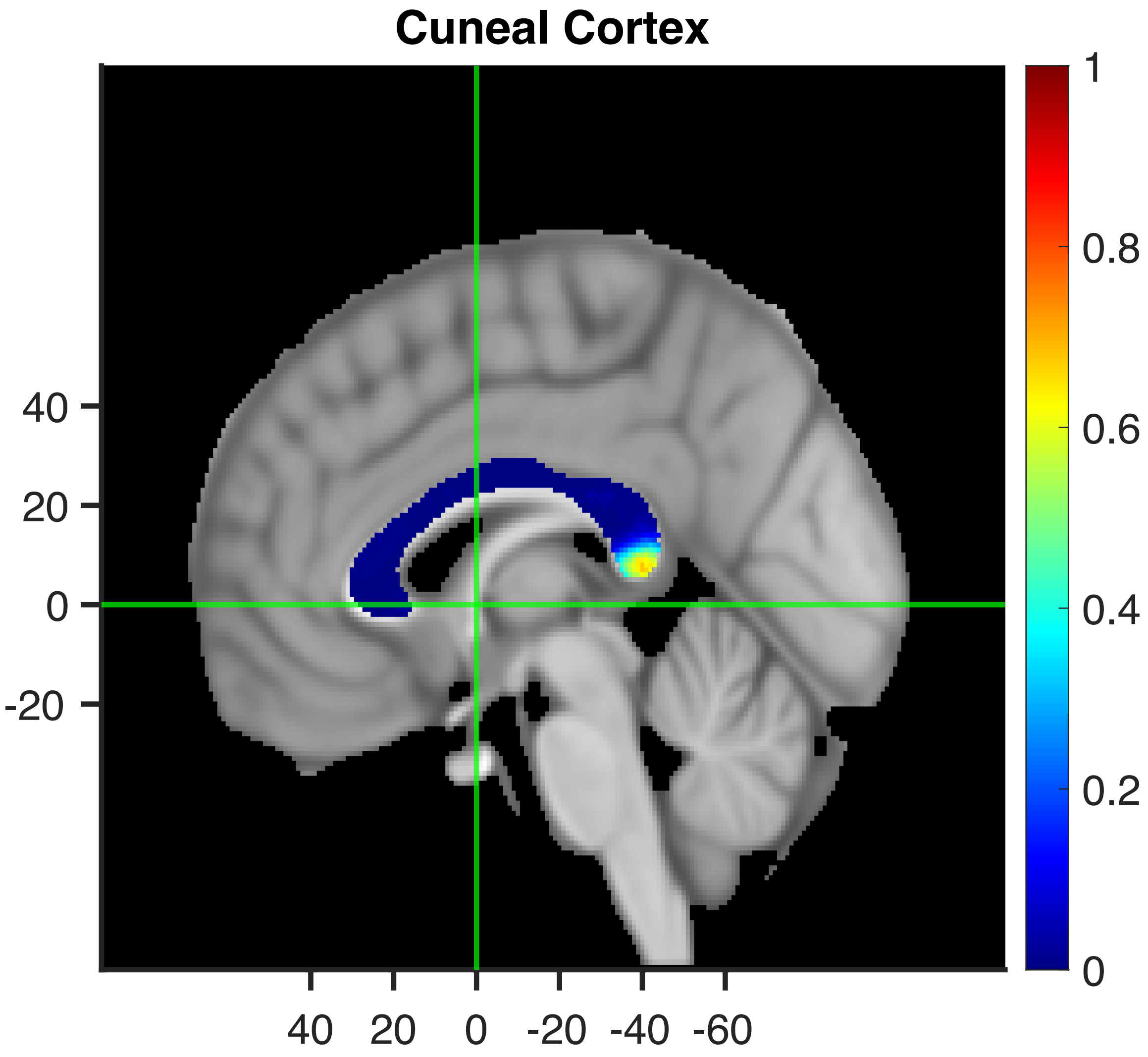

Cuneal Cortex
40
20
0
-20
40
20
0
-20
-40
-60
1
0.8
0.6
0.4
0.2
0

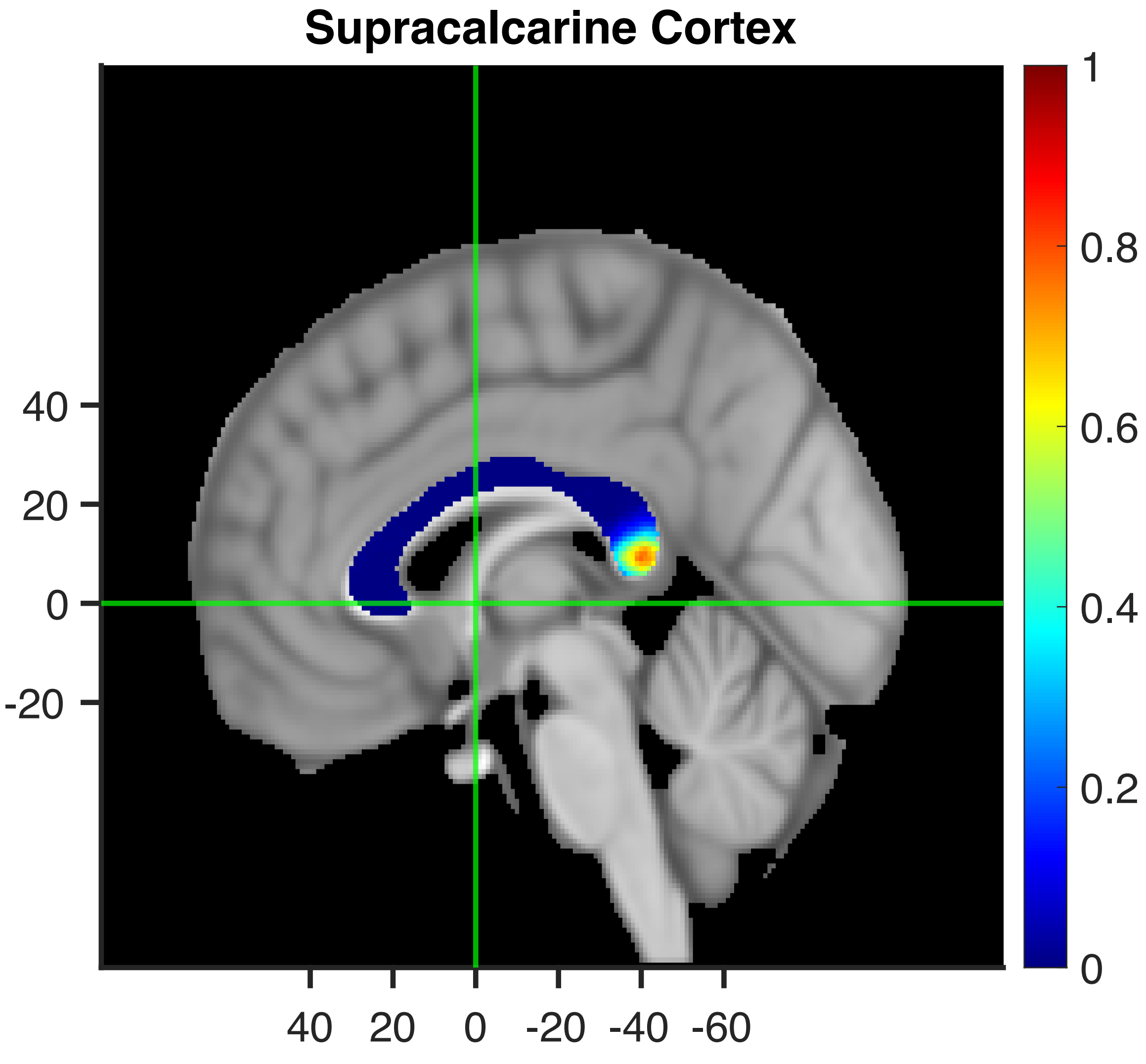

Supracalcarine Cortex
40
20
0
-20
40
20
0
-20
-40
-60
1
0.8
0.6
0.4
0.2
0

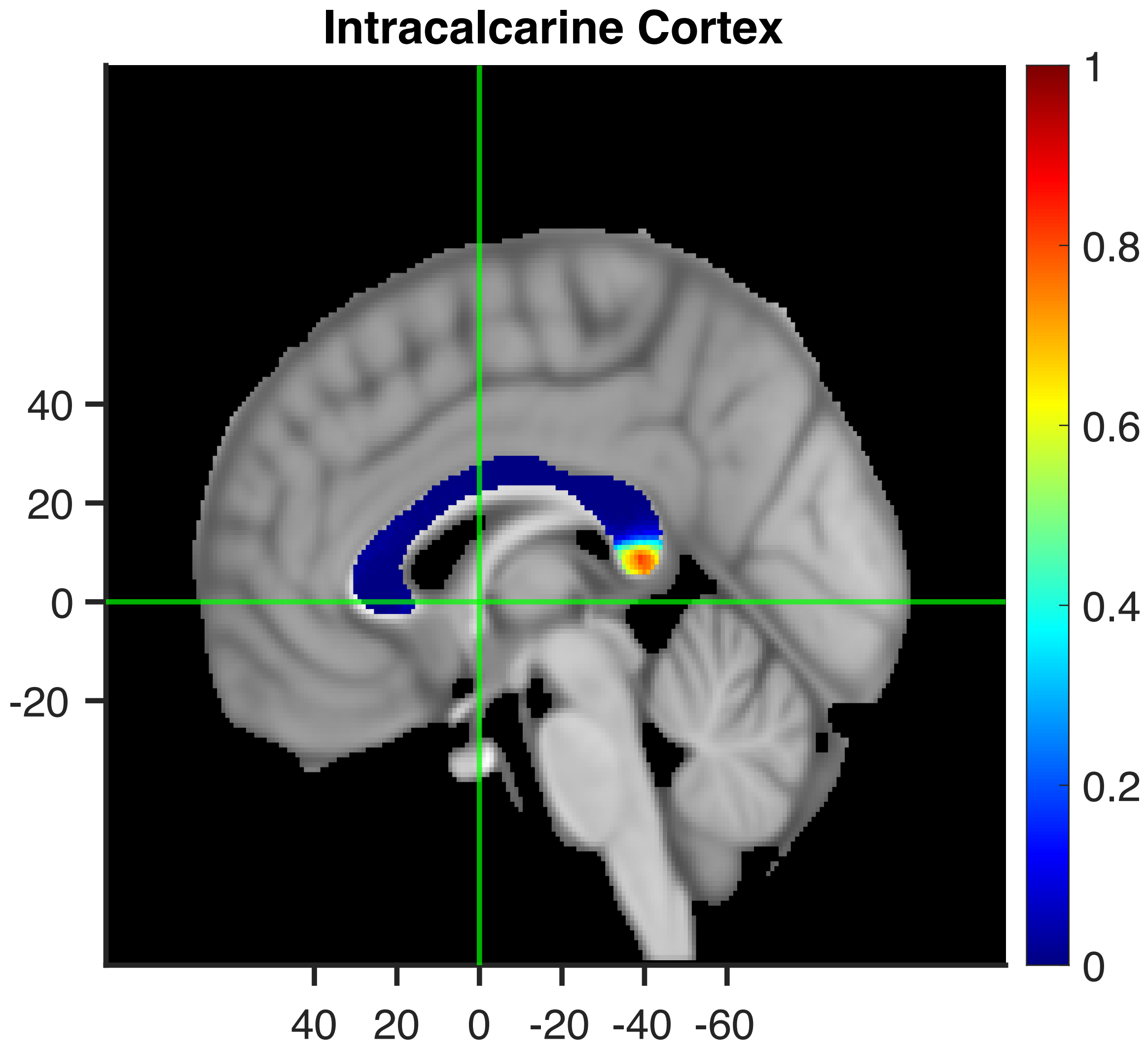
Intracalcarine Cortex
40
20
0
-20
40 20 0 -20 -40 -60
1
0.8
0.6
0.4
0.2
0

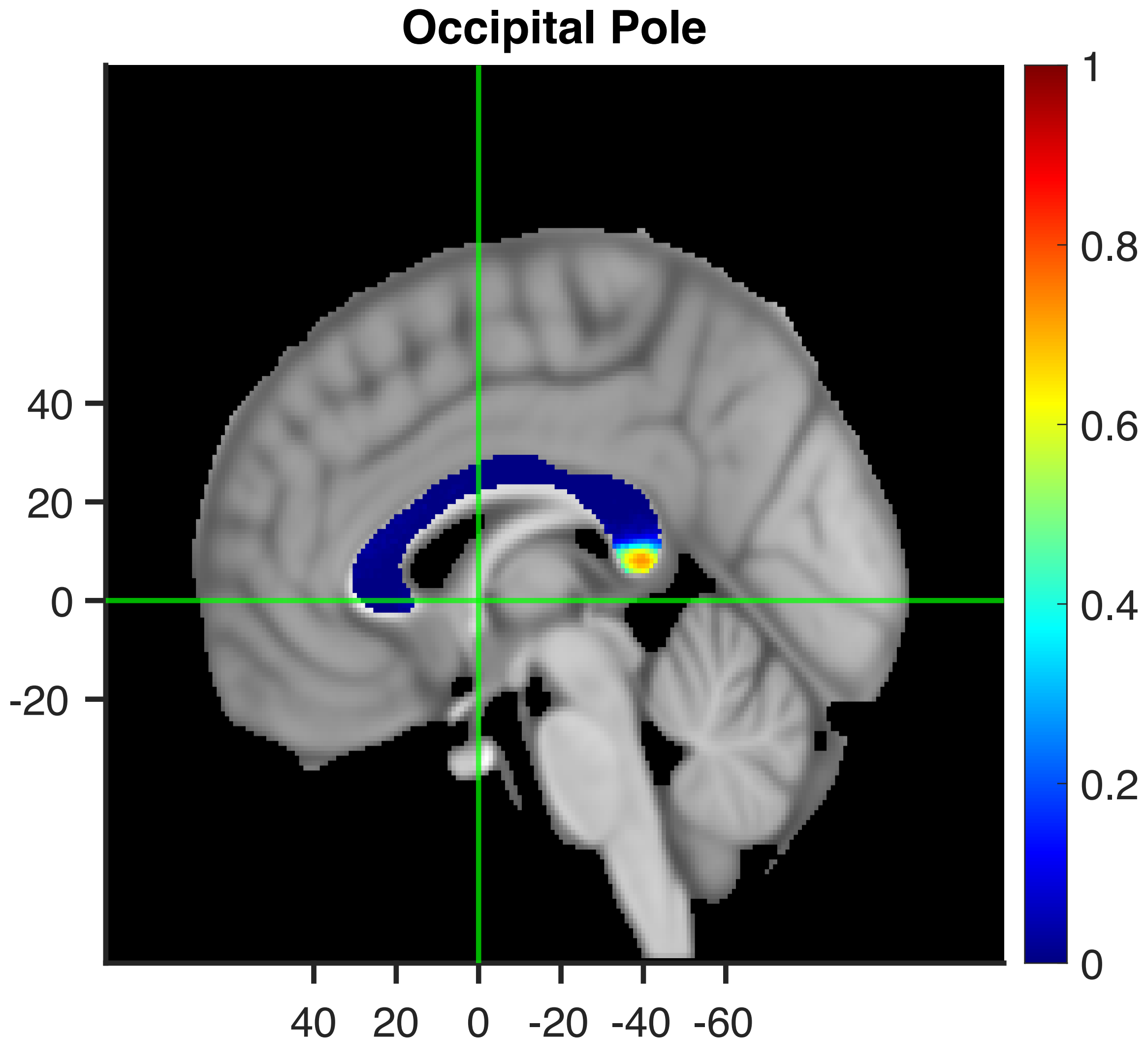
Occipital Pole
40
20
0
-20
40 20 0 -20 -40 -60
1
0.8
0.6
0.4
0.2
0

#### Lingual Gyrus

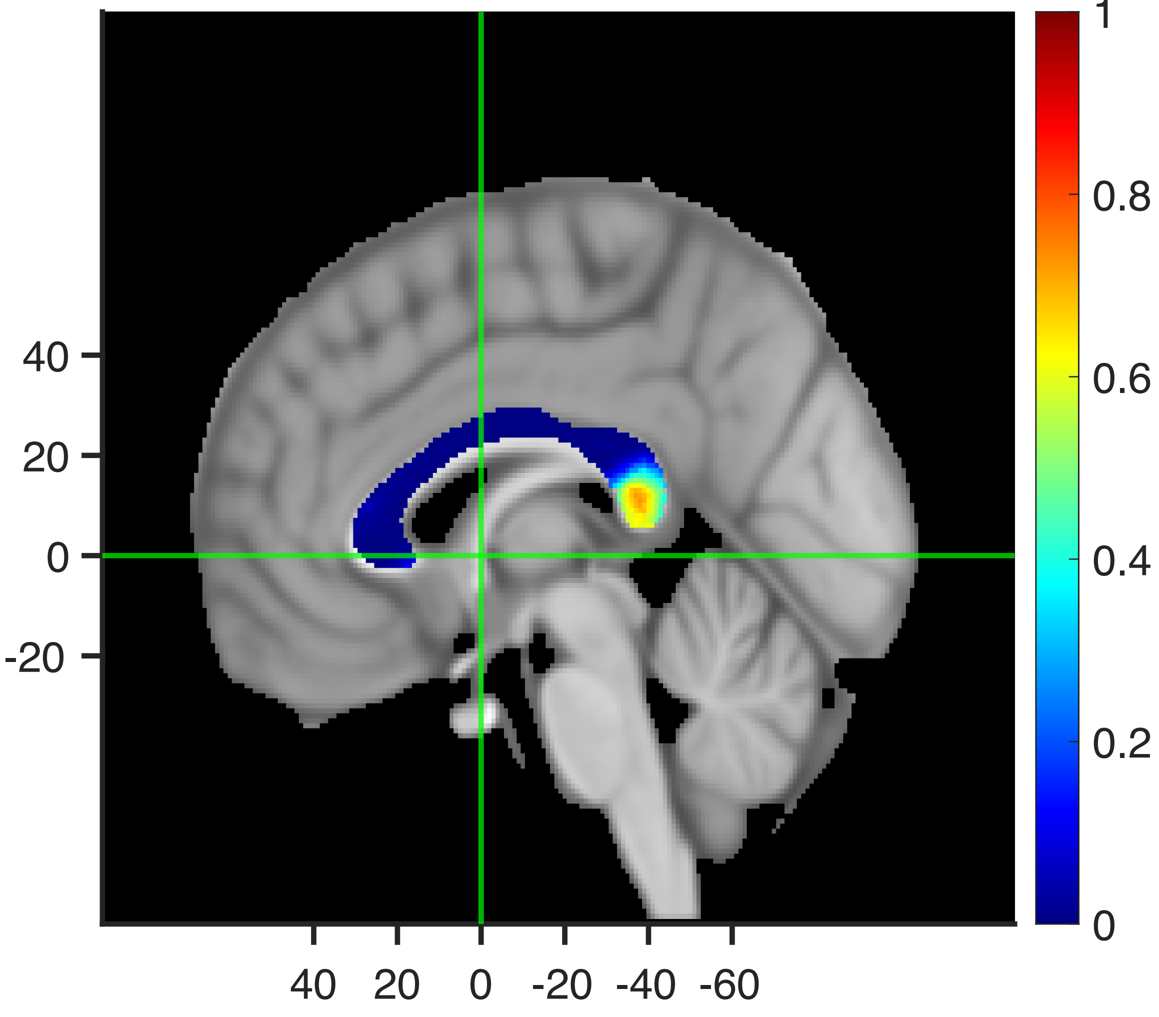

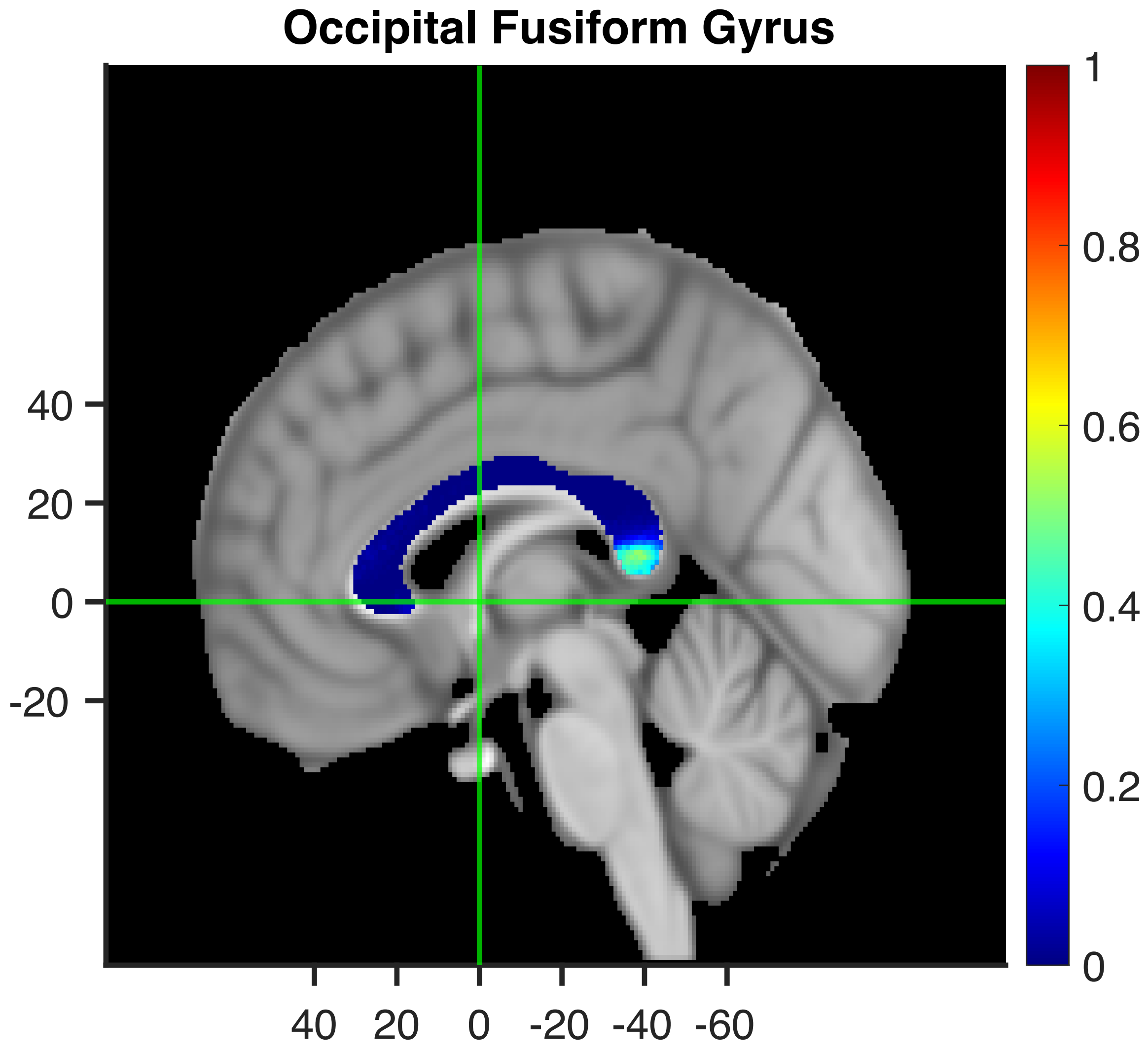
Occipital Fusiform Gyrus
40
20
0
-20
40
20
0
-20
-40
-60
1
0.8
0.6
0.4
0.2
0

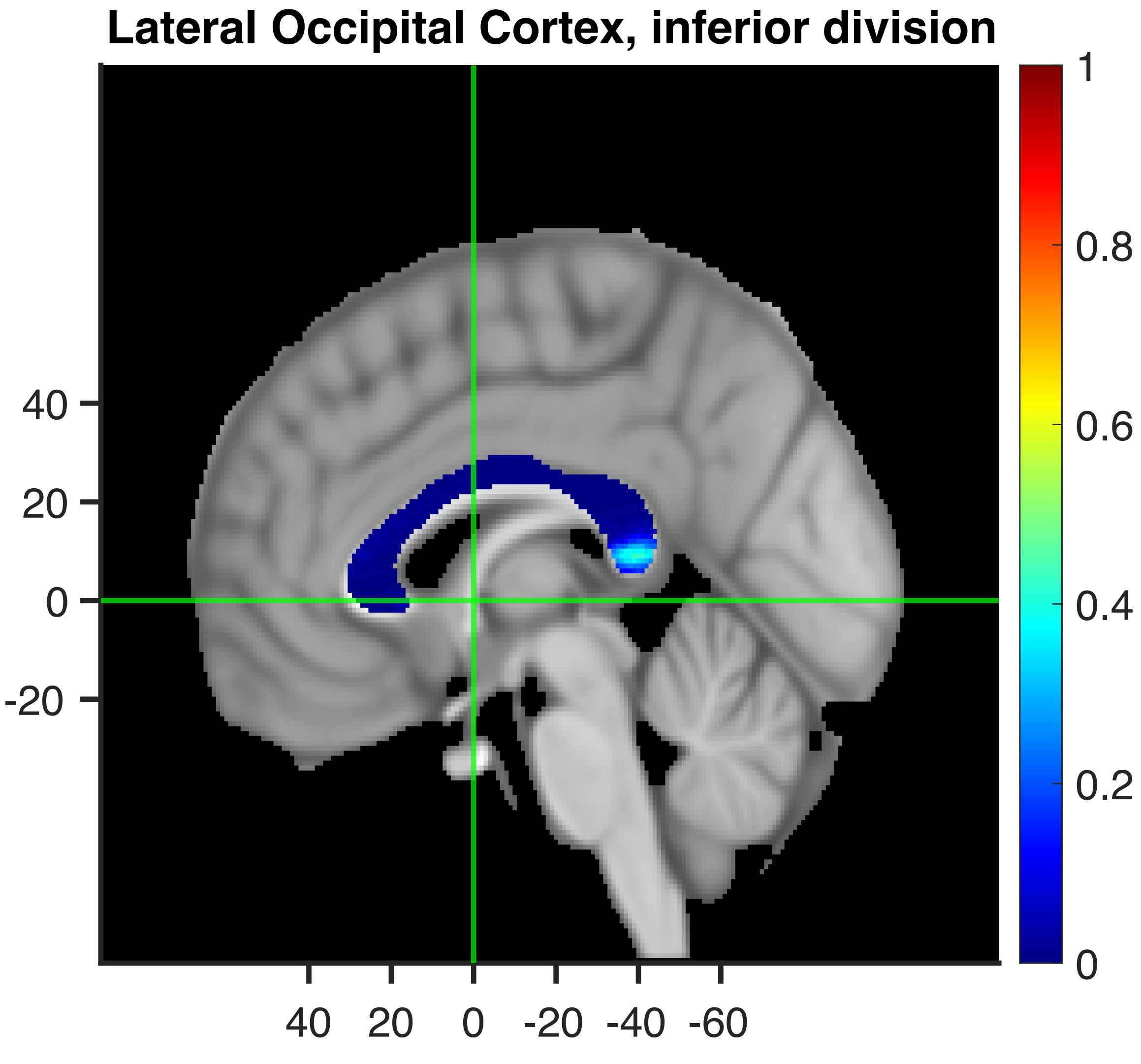

Lateral Occipital Cortex, inferior division
40
20
0
-20
40 20 0 -20 -40 -60
1
0.8
0.6
0.4
0.2
0

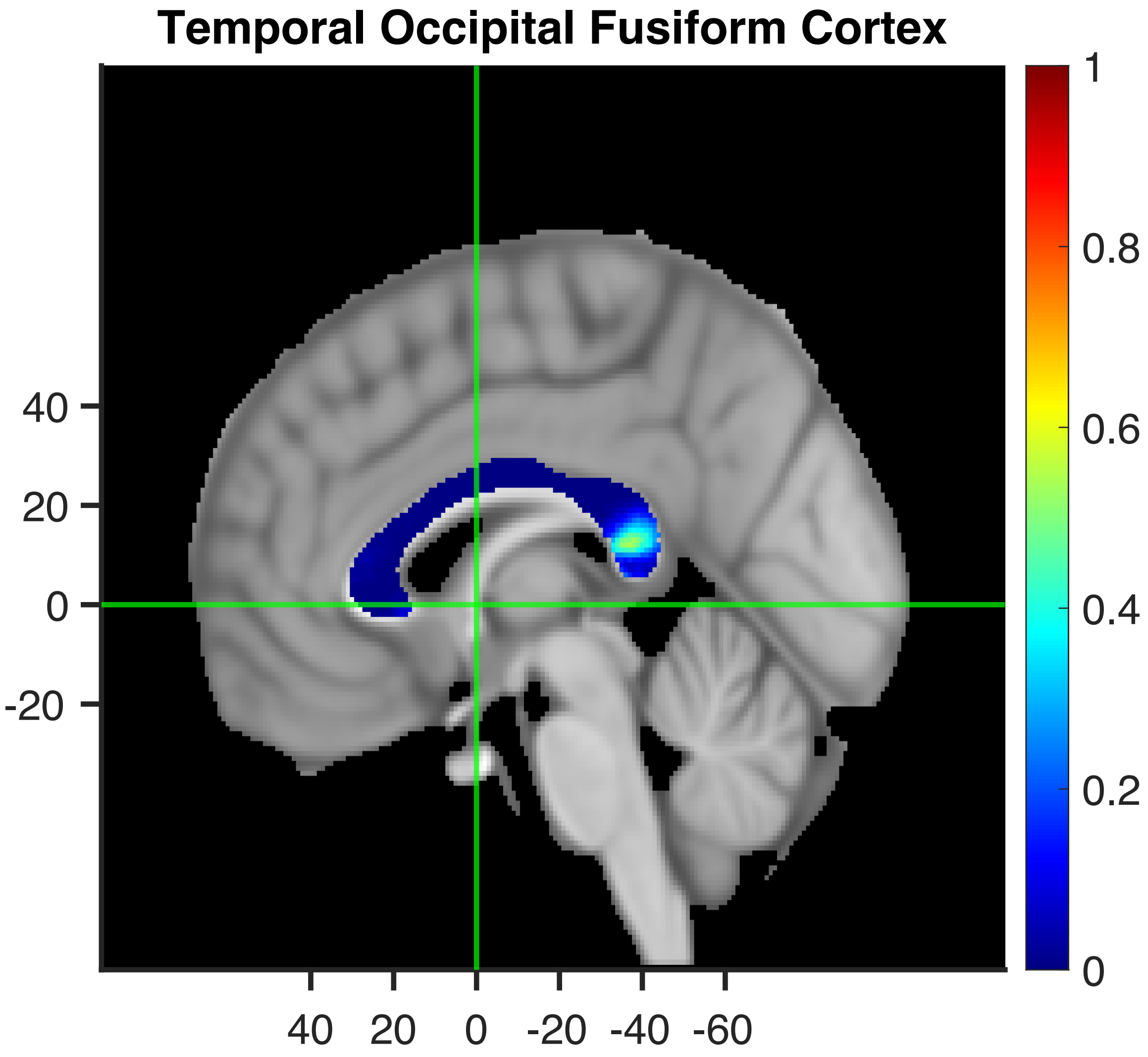

Temporal Occipital Fusiform Cortex
40
20
0
-20
40 20 0 -20 -40 -60
1
0.8
0.6
0.4
0.2
0

#### Inferior Temporal Gyrus, temporooccipital part

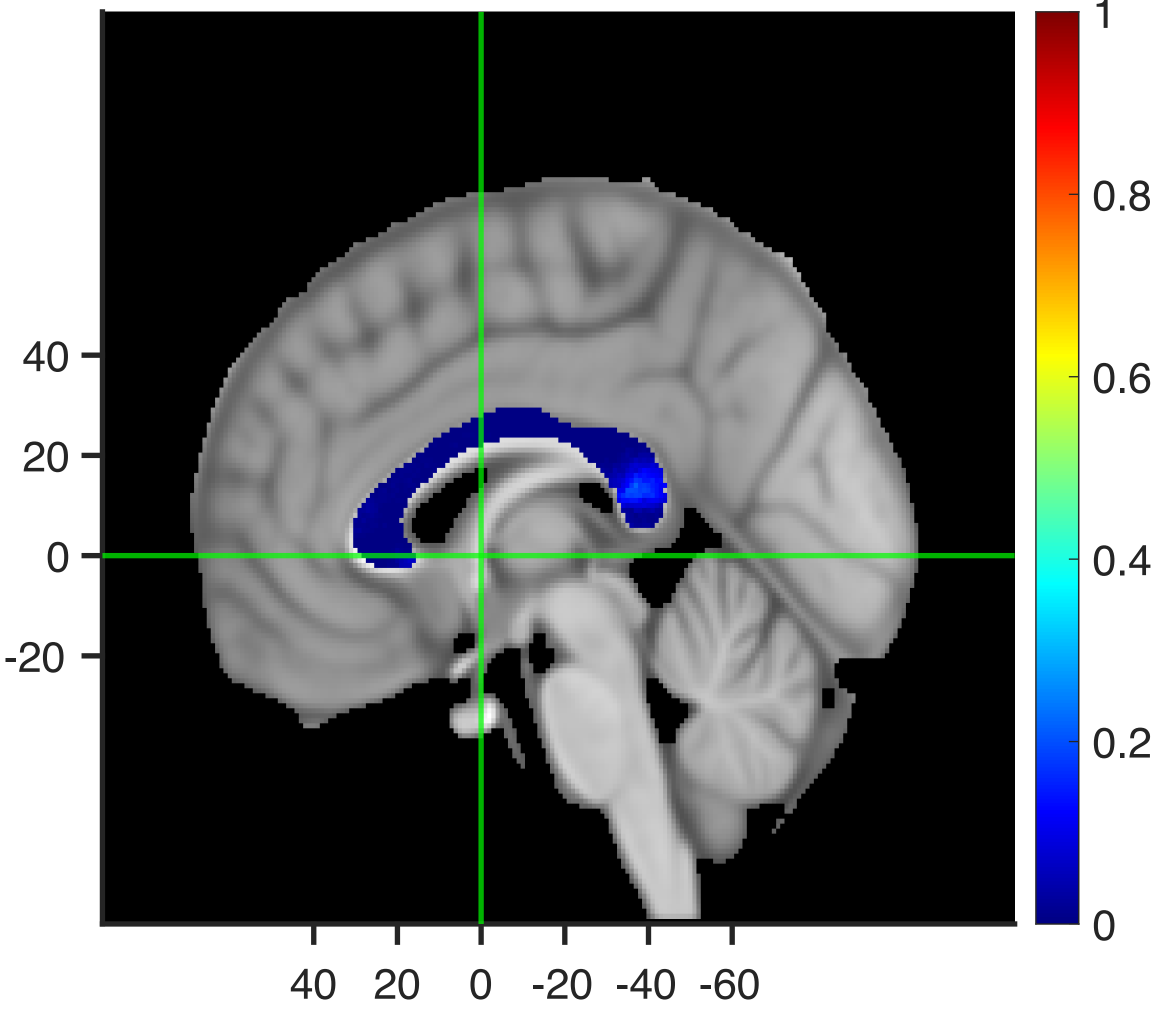

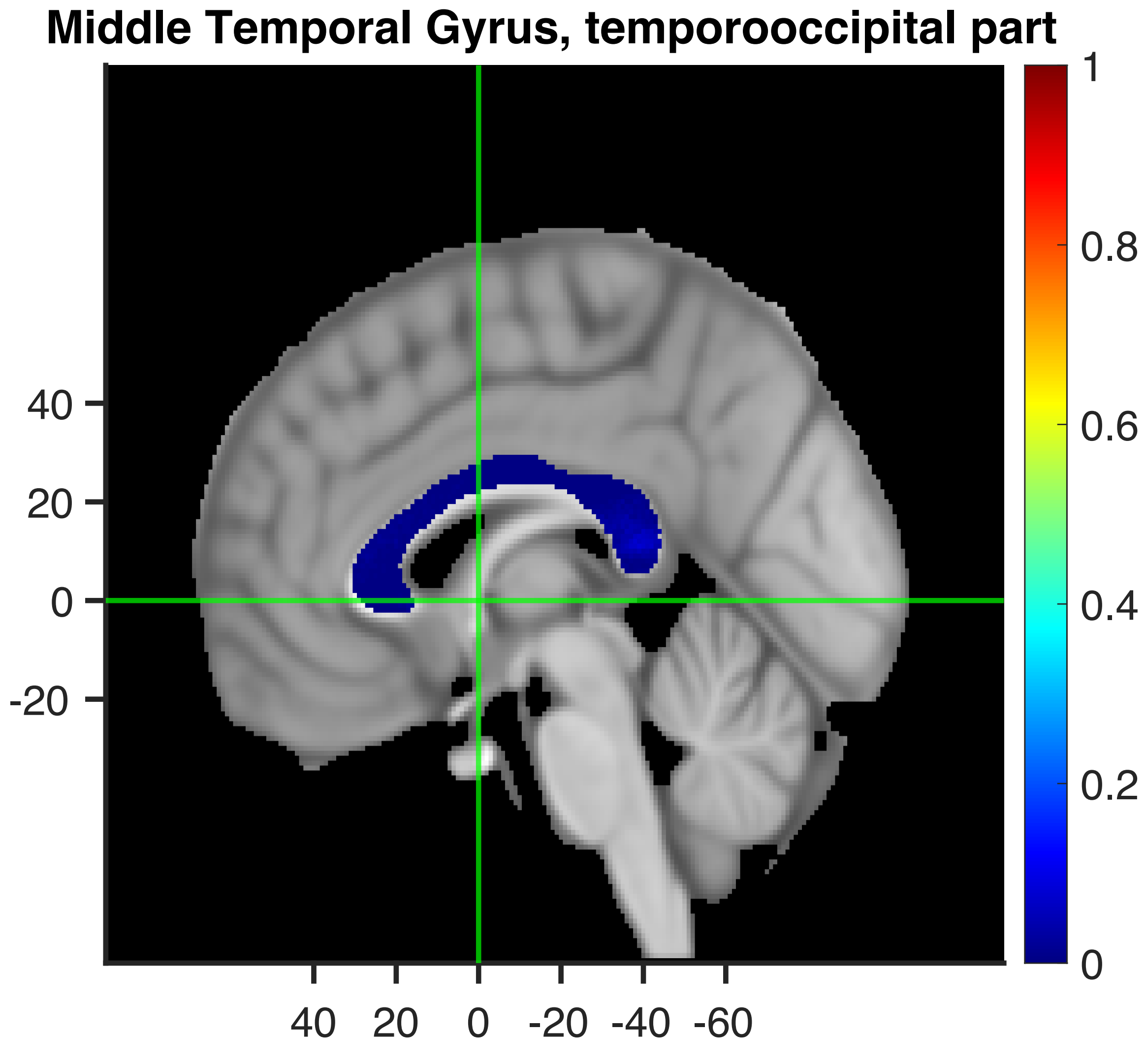

Middle Temporal Gyrus, temporooccipital part
40
20
0
-20
40
20
0
-20
-40
-60
1
0.8
0.6
0.4
0.2
0

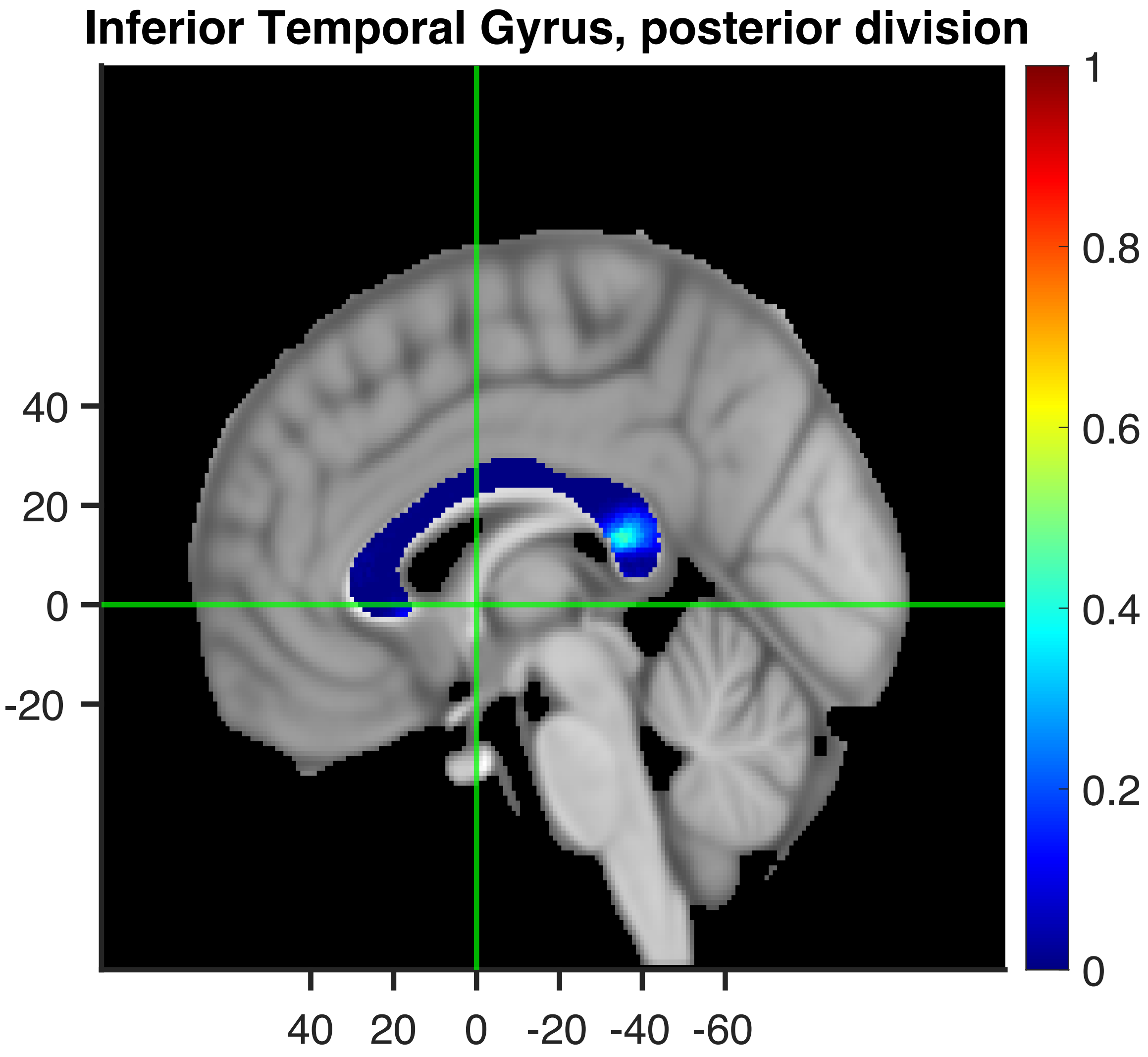

Inferior Temporal Gyrus, posterior division
40
20
0
-20
40 20 0 -20 -40 -60
1
0.8
0.6
0.4
0.2
0

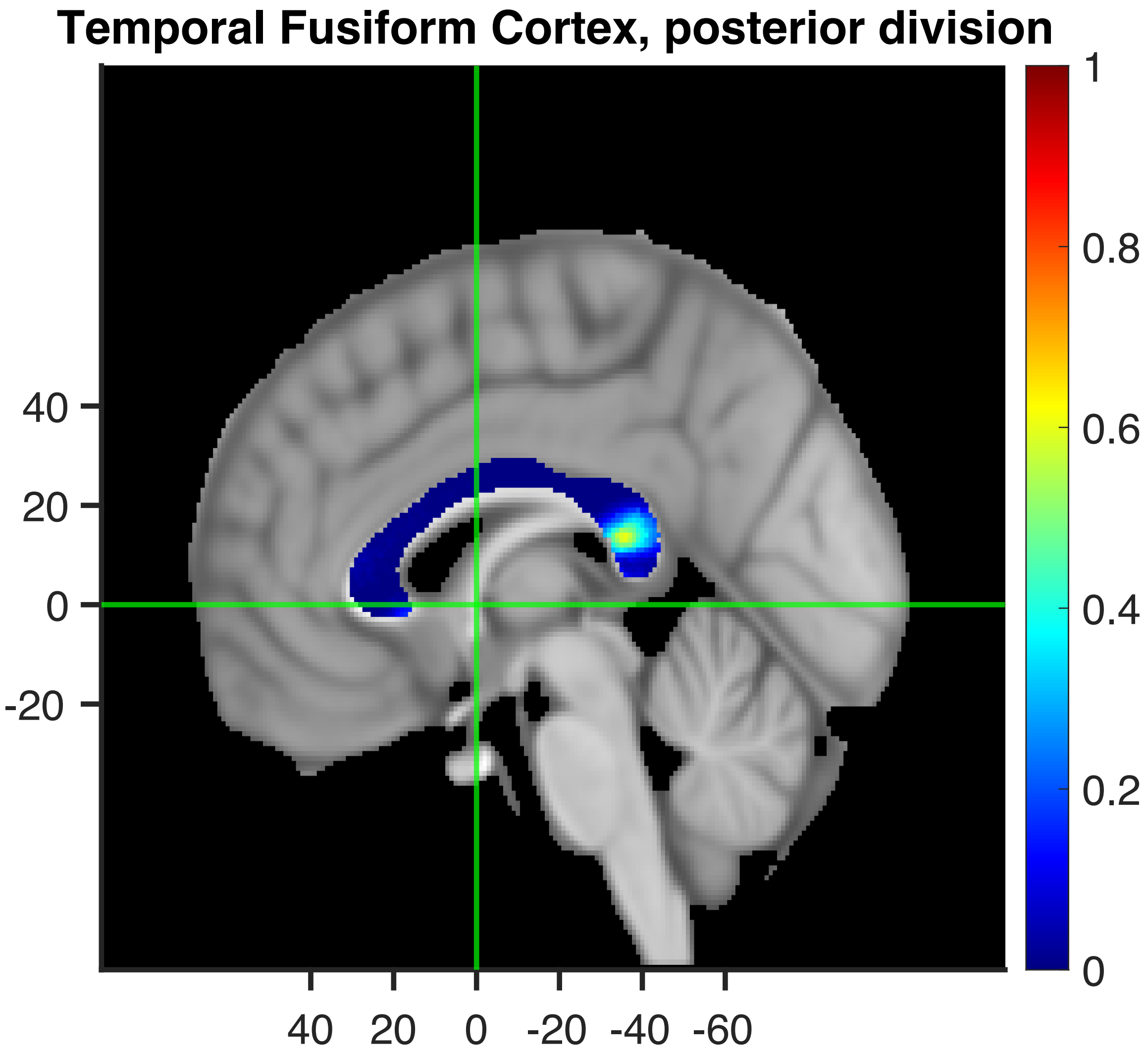

Temporal Fusiform Cortex, posterior division
40
20
0
-20
40
20
0
-20
-40
-60
1
0.8
0.6
0.4
0.2
0

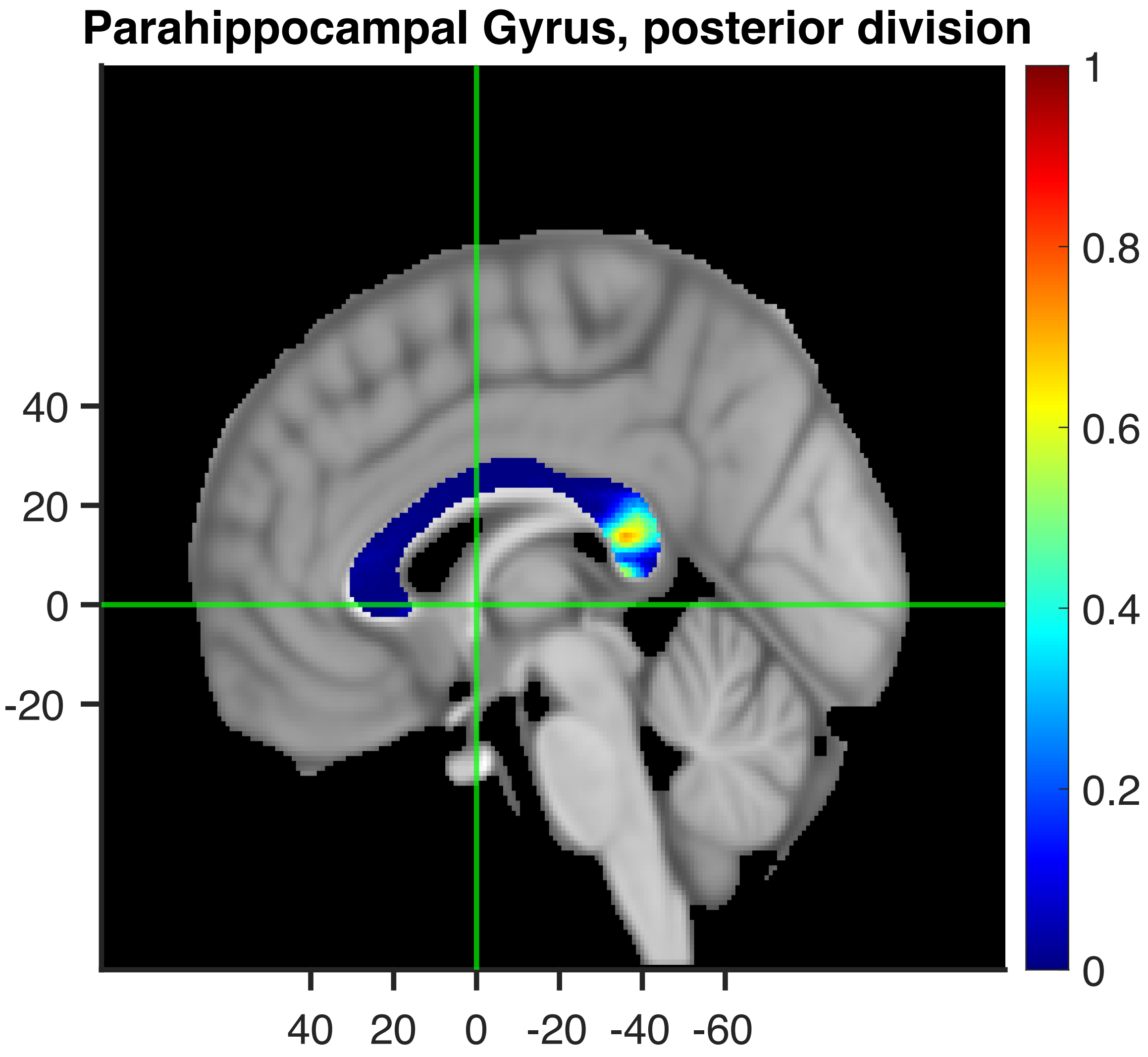

Parahippocampal Gyrus, posterior division
40
20
0
-20
40
20
0
-20
-40
-60
1
0.8
0.6
0.4
0.2
0

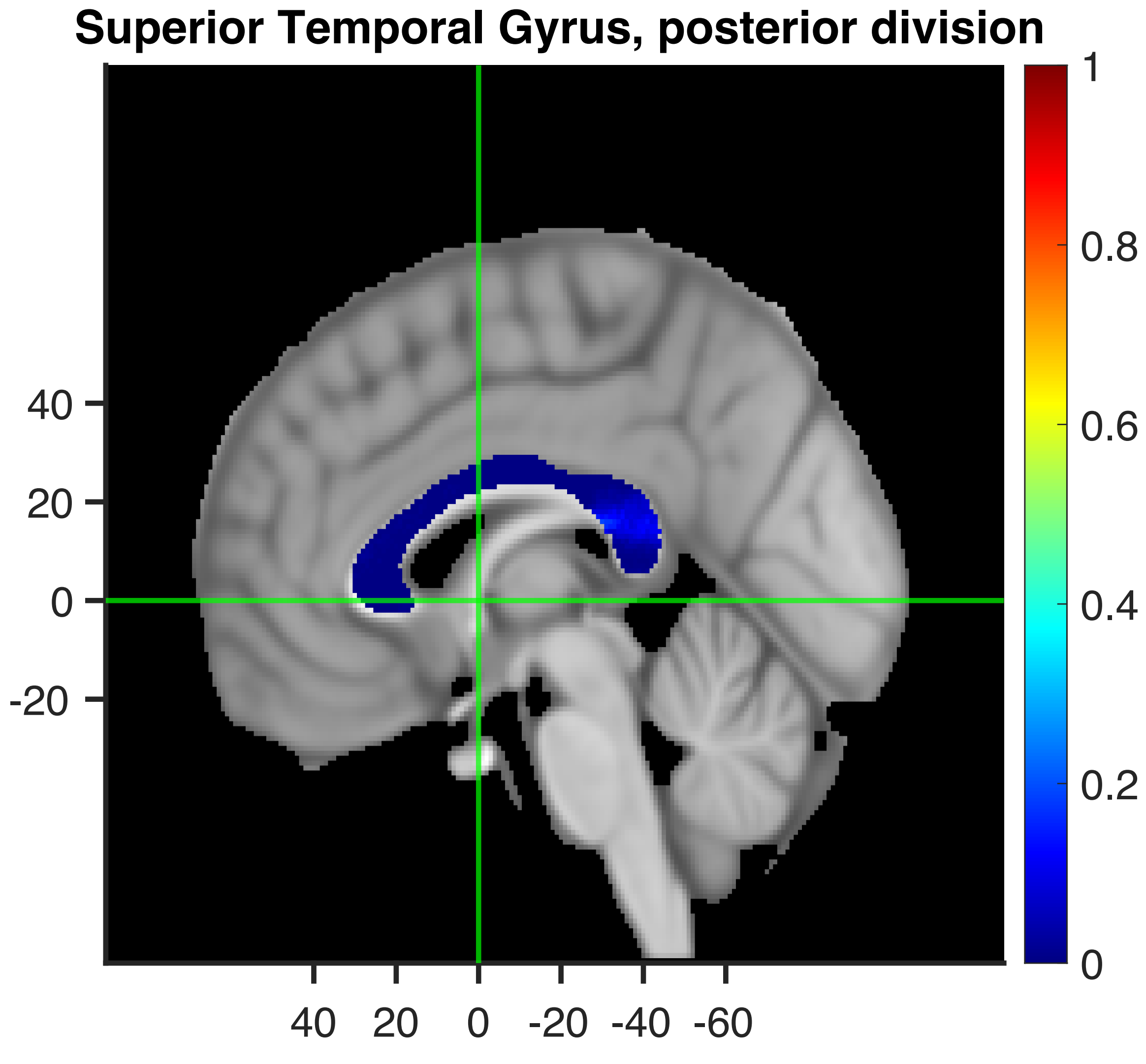

Superior Temporal Gyrus, posterior division
40
20
0
-20
40
20
0
-20
-40
-60
1
0.8
0.6
0.4
0.2
0

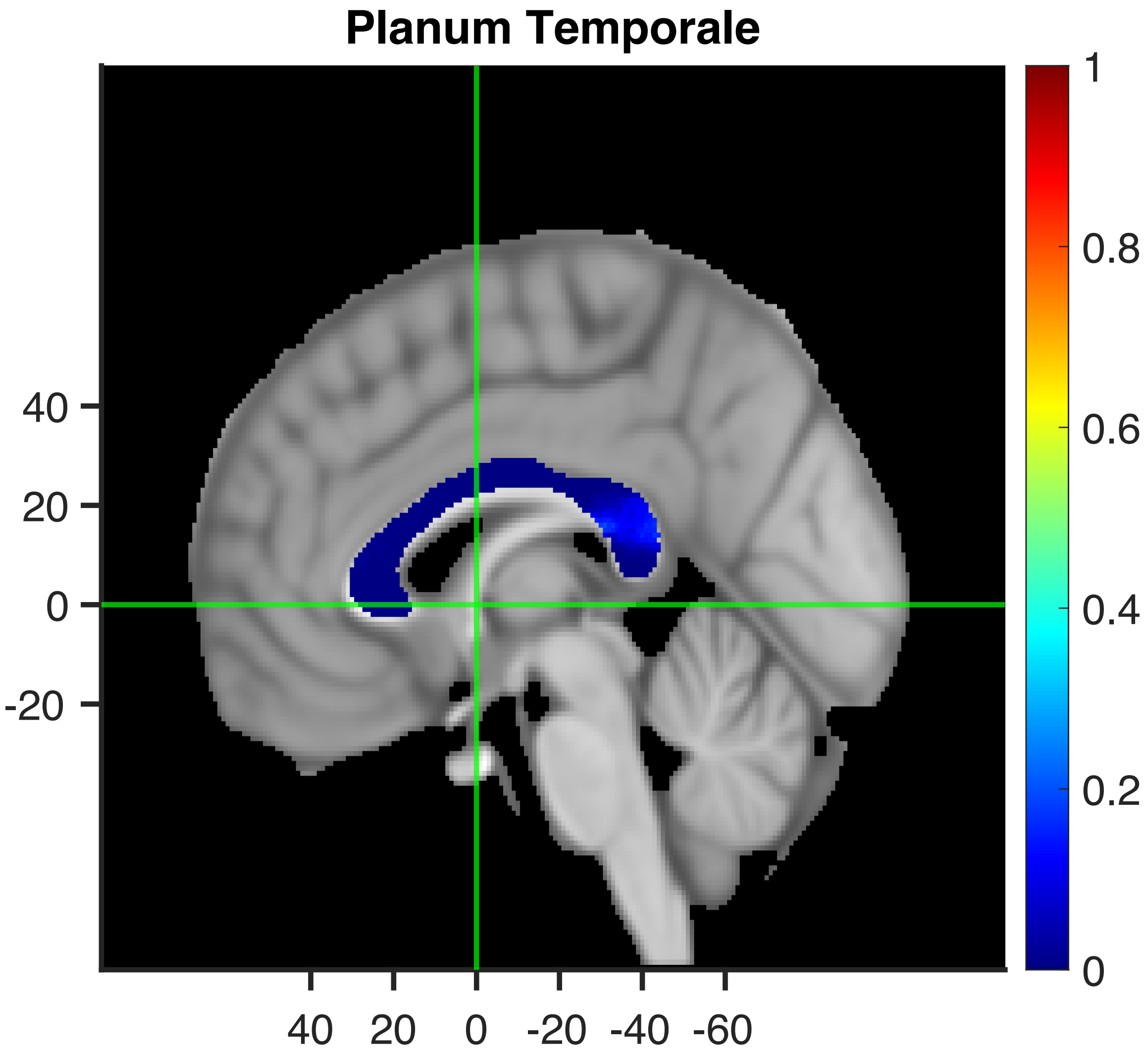

Planum Temporale
40
20
0
-20
40
20
0
-20
-40
-60
1
0.8
0.6
0.4
0.2
0

Heschl's Gyrus (includes H1 and H2)

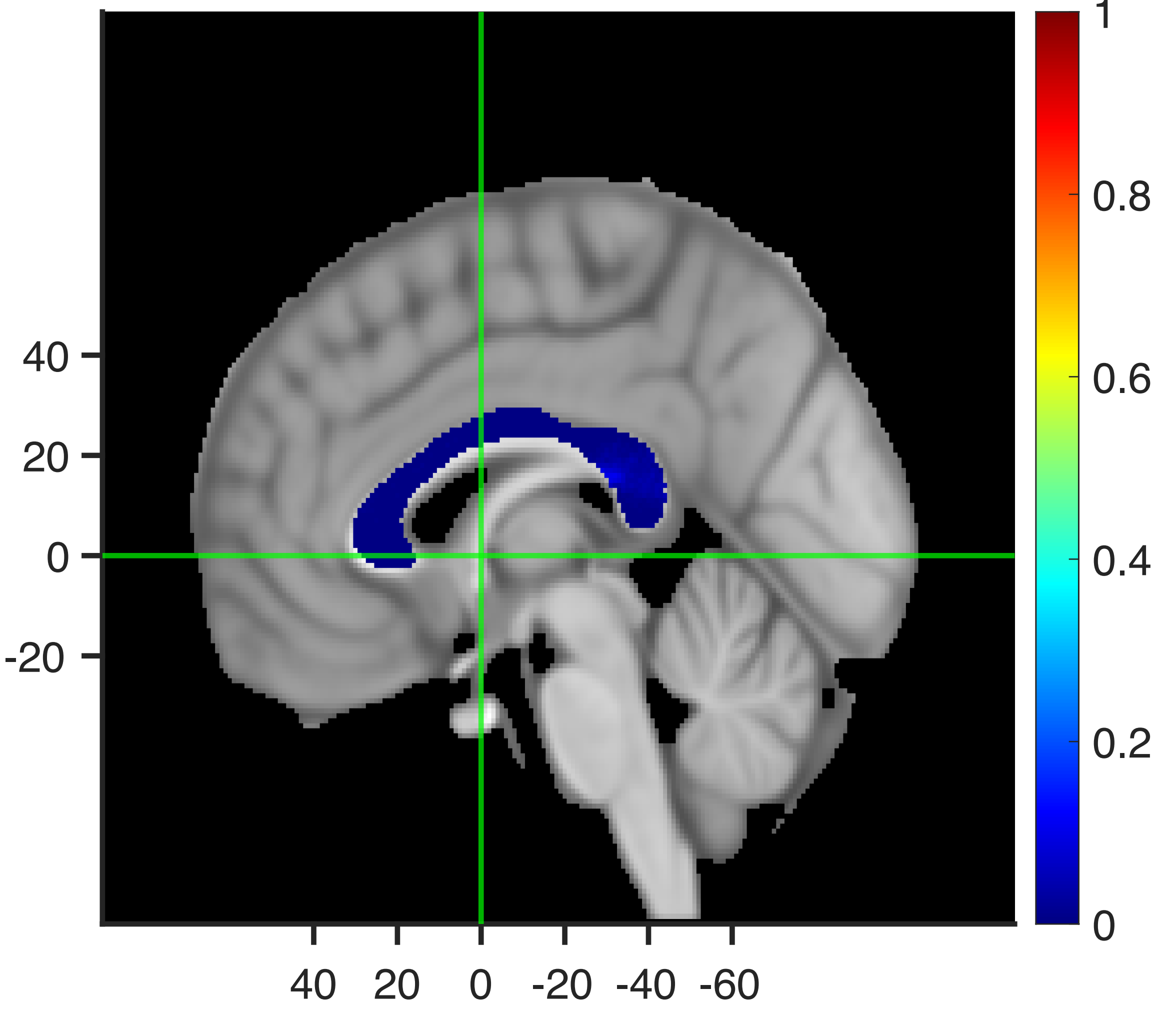

#### Middle Temporal Gyrus, posterior division

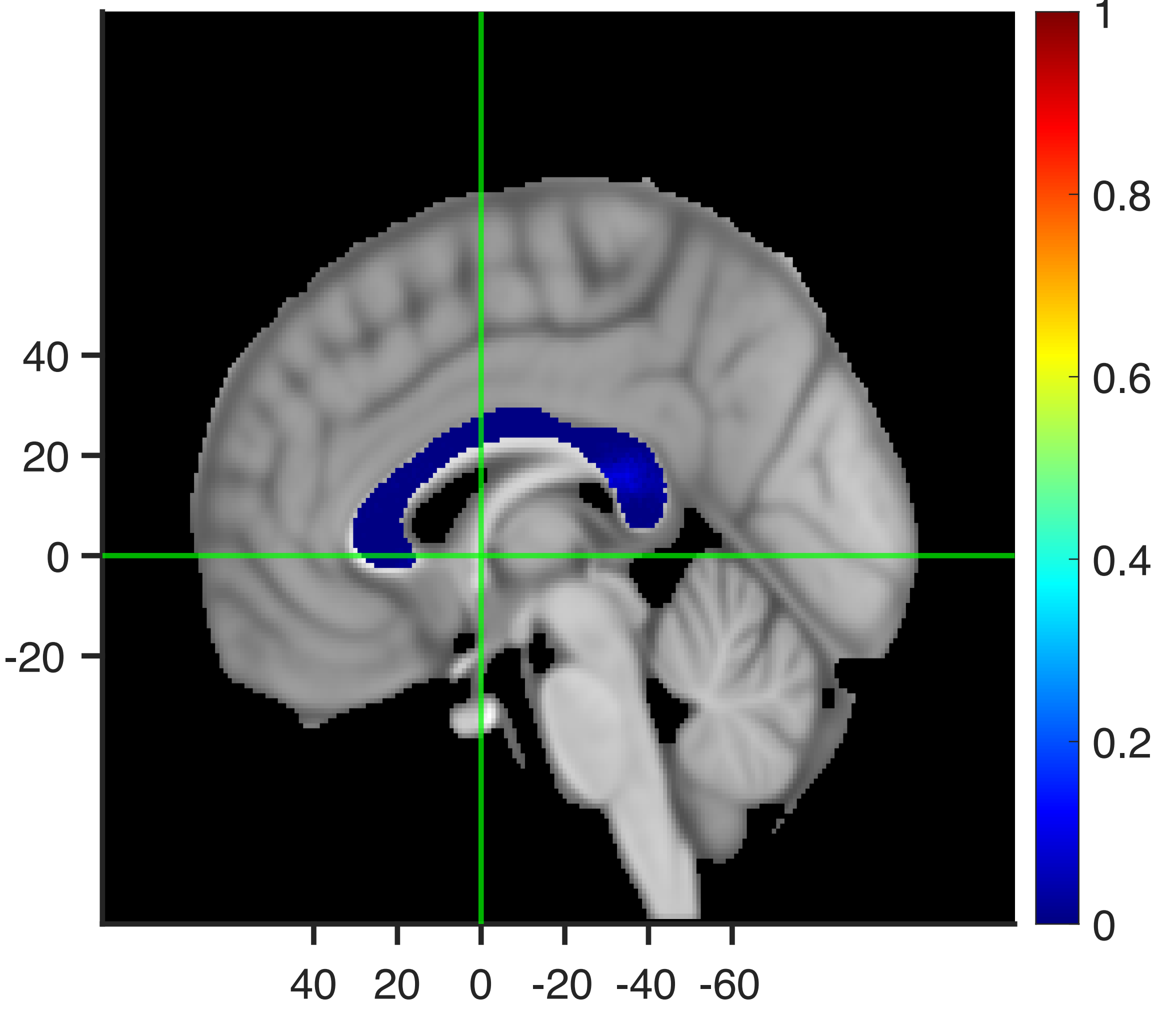

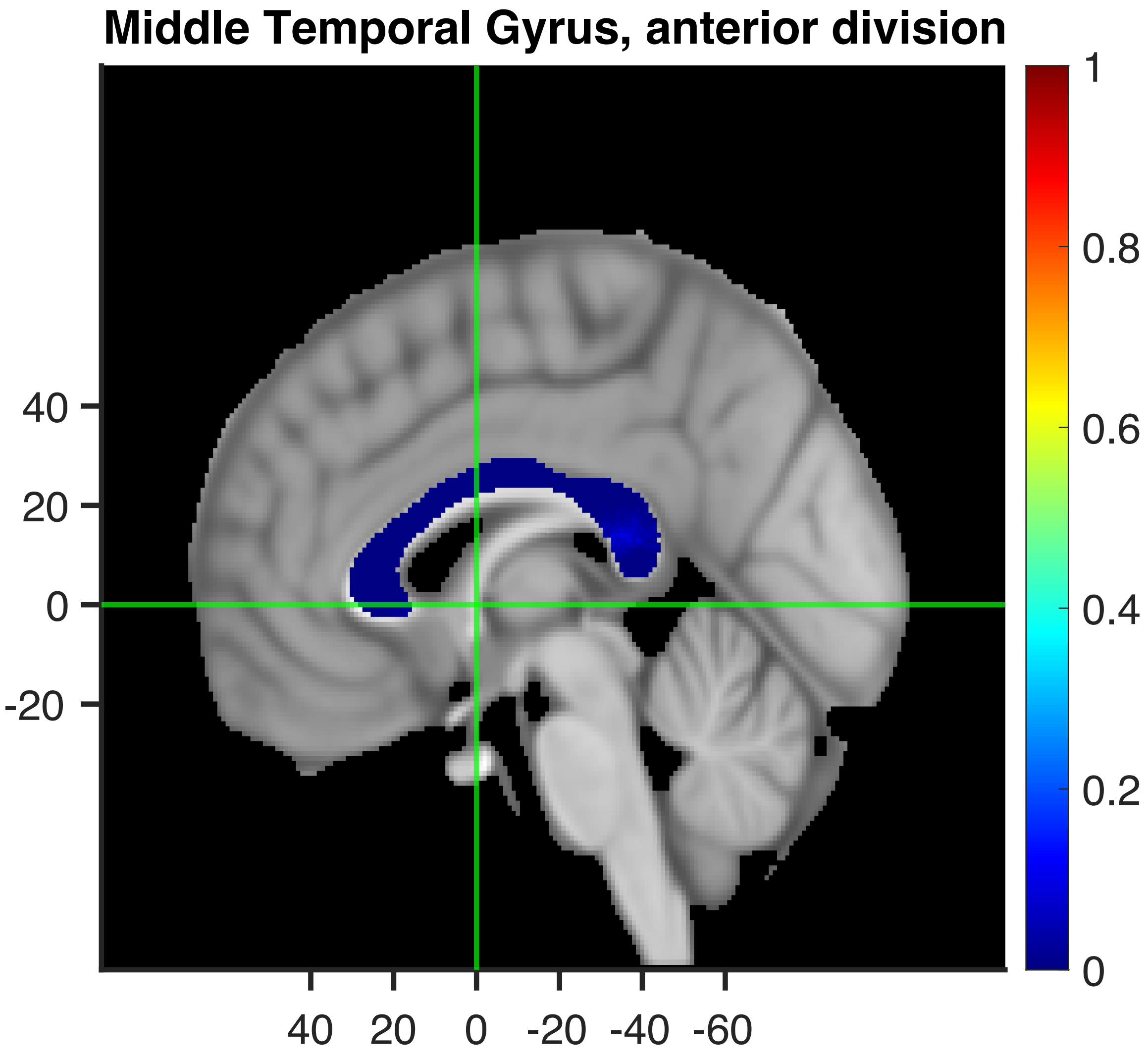

Middle Temporal Gyrus, anterior division
40
20
0
-20
40
20
0
-20
-40
-60
1
0.8
0.6
0.4
0.2
0

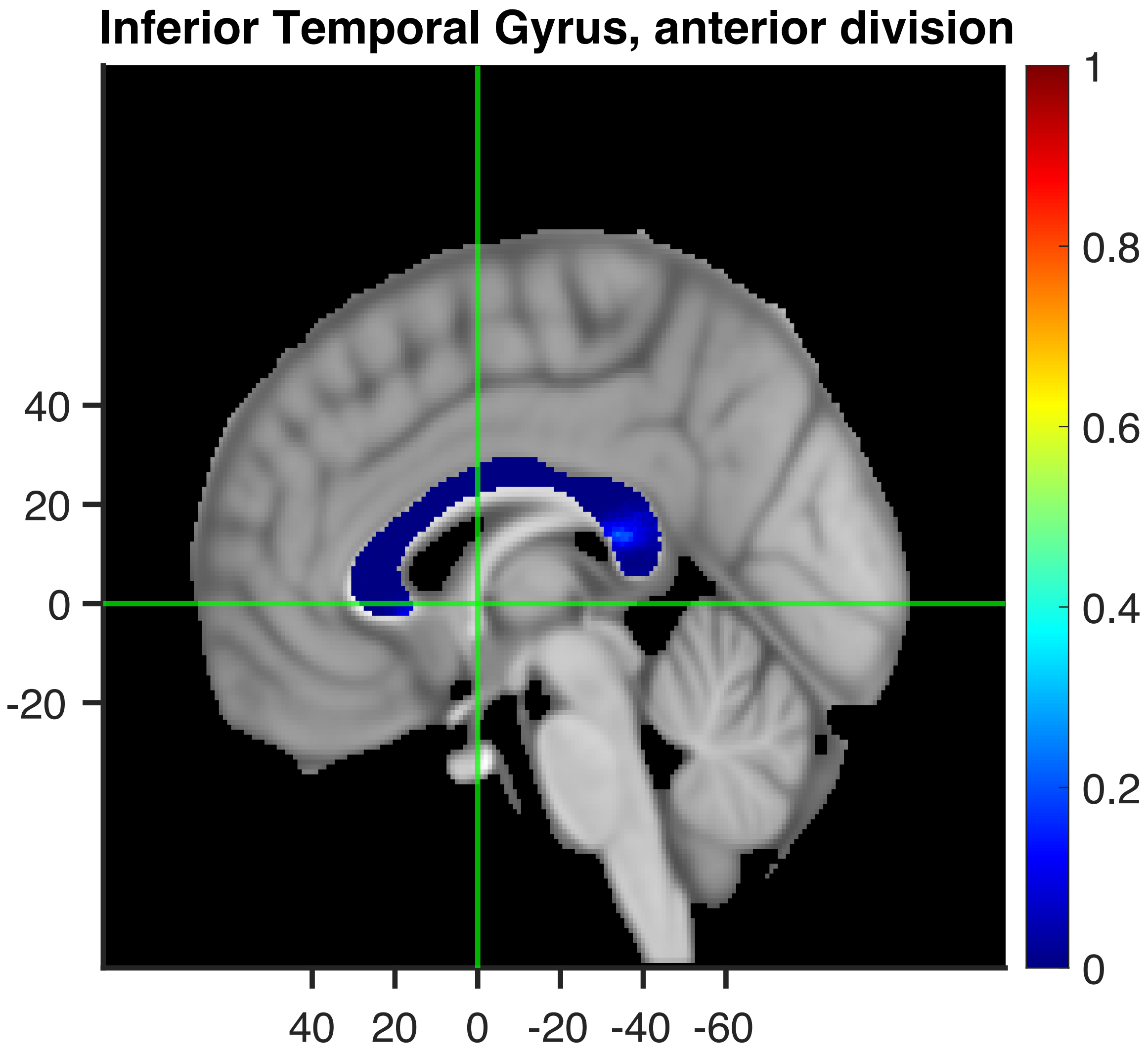
Inferior Temporal Gyrus, anterior division
40
20
0
-20
40 20 0 -20 -40 -60
1
0.8
0.6
0.4
0.2
0

**Temporal Fusiform Cortex, anterior division**

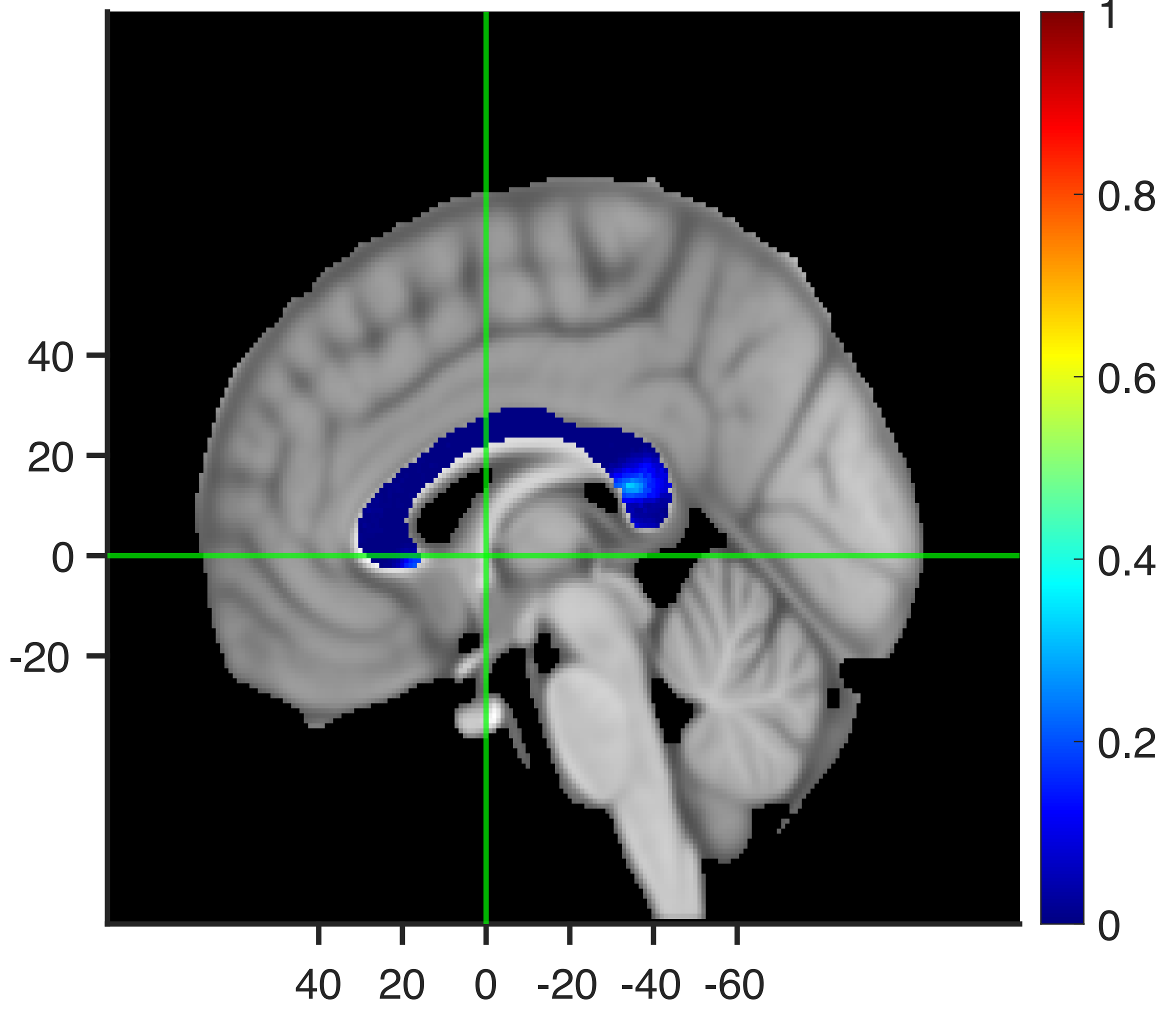

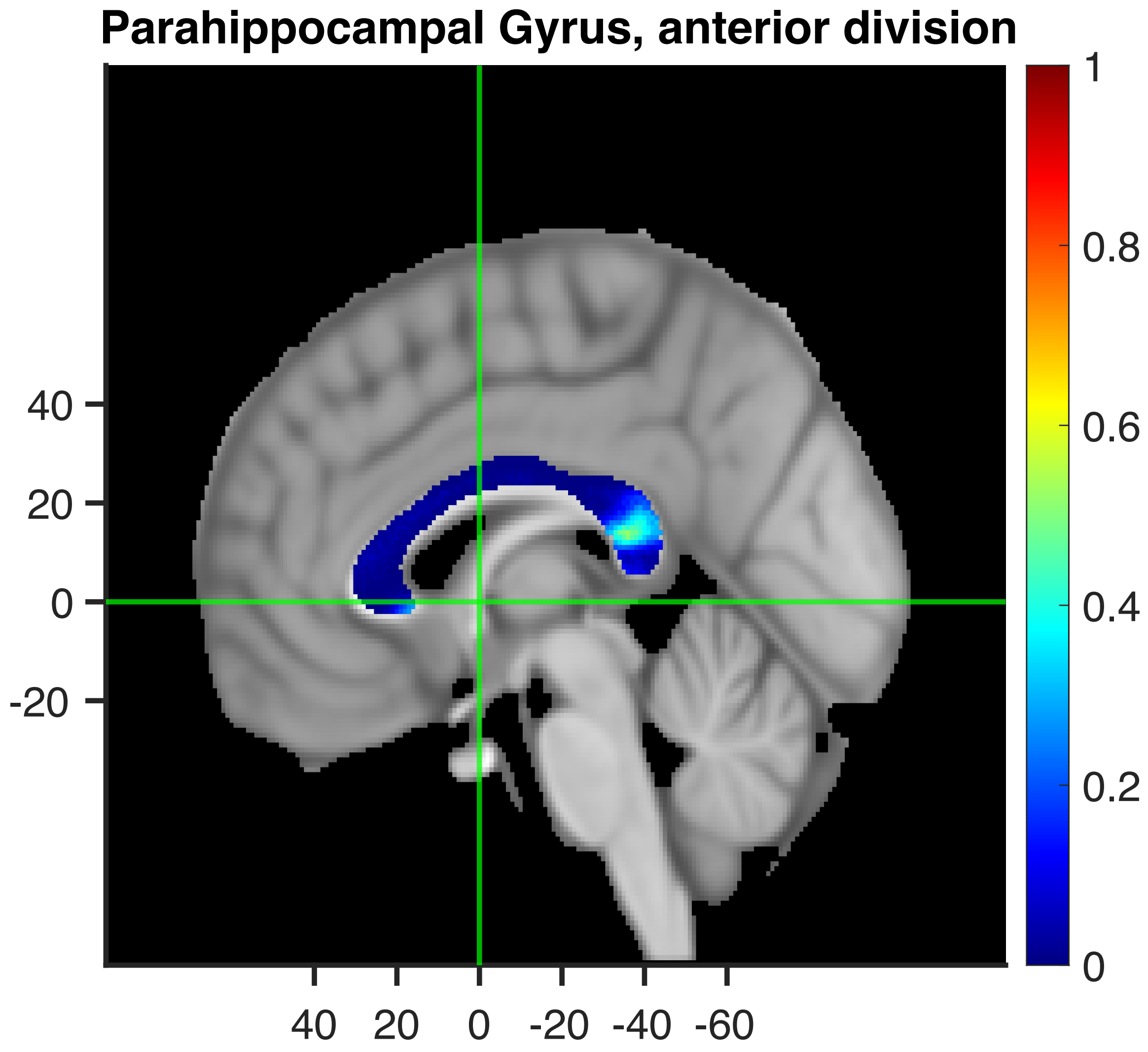
Parahippocampal Gyrus, anterior division
40
20
0
-20
40 20 0 -20 -40 -60
1
0.8
0.6
0.4
0.2
0

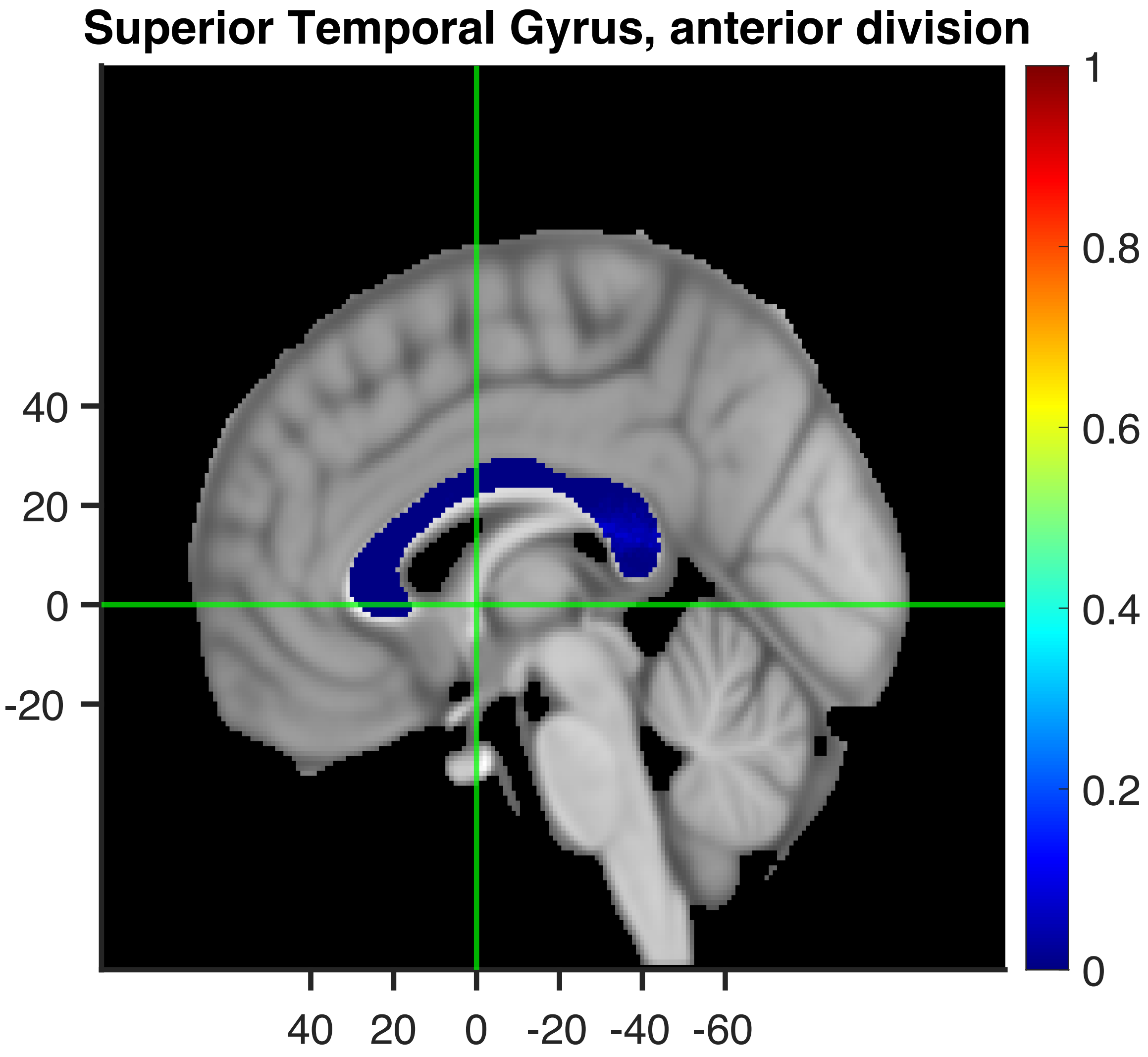
Superior Temporal Gyrus, anterior division
40
20
0
-20
40 20 0 -20 -40 -60
1
0.8
0.6
0.4
0.2
0

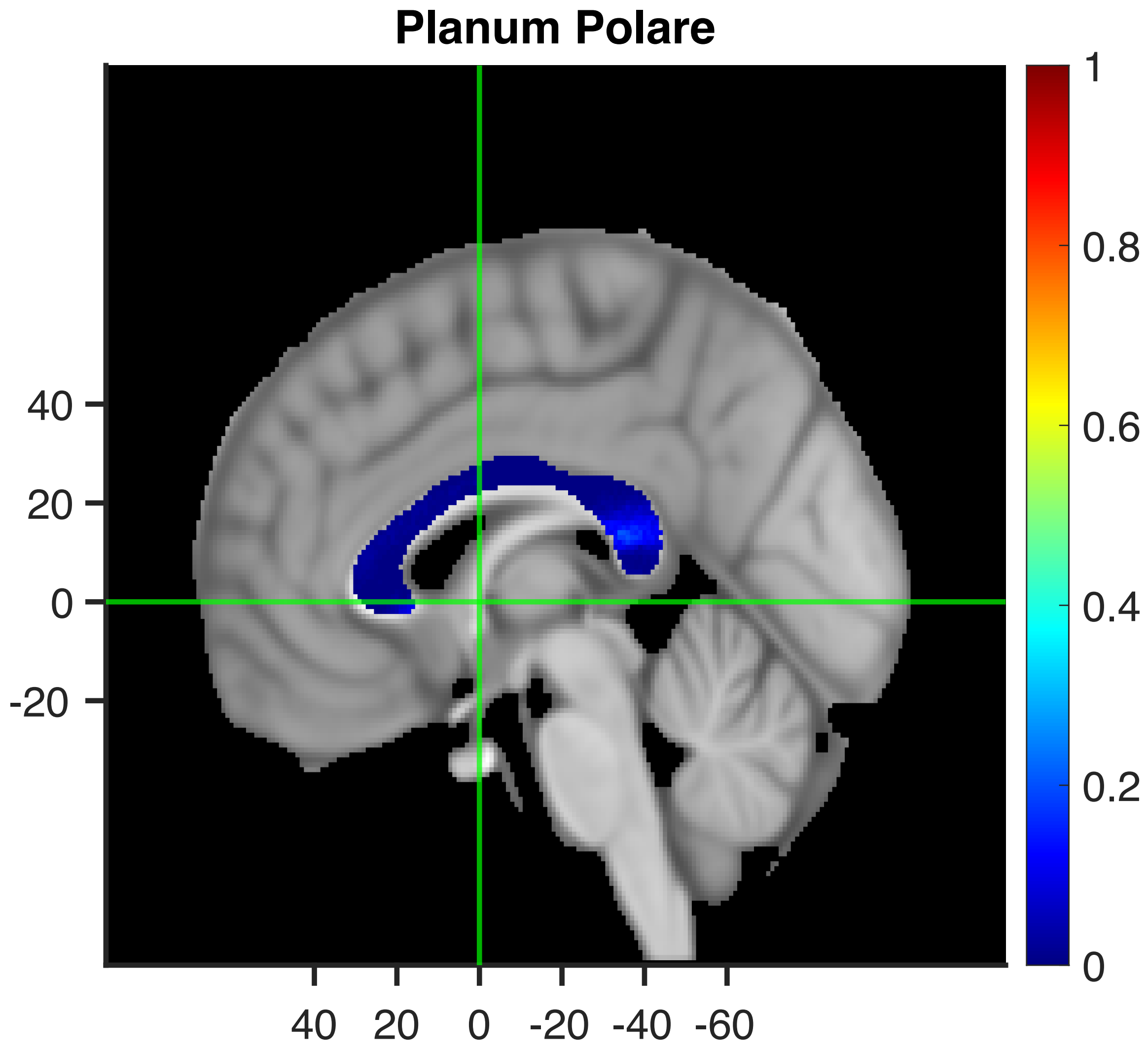
Planum Polare
40
20
0
-20
40 20 0 -20 -40 -60
1
0.8
0.6
0.4
0.2
0

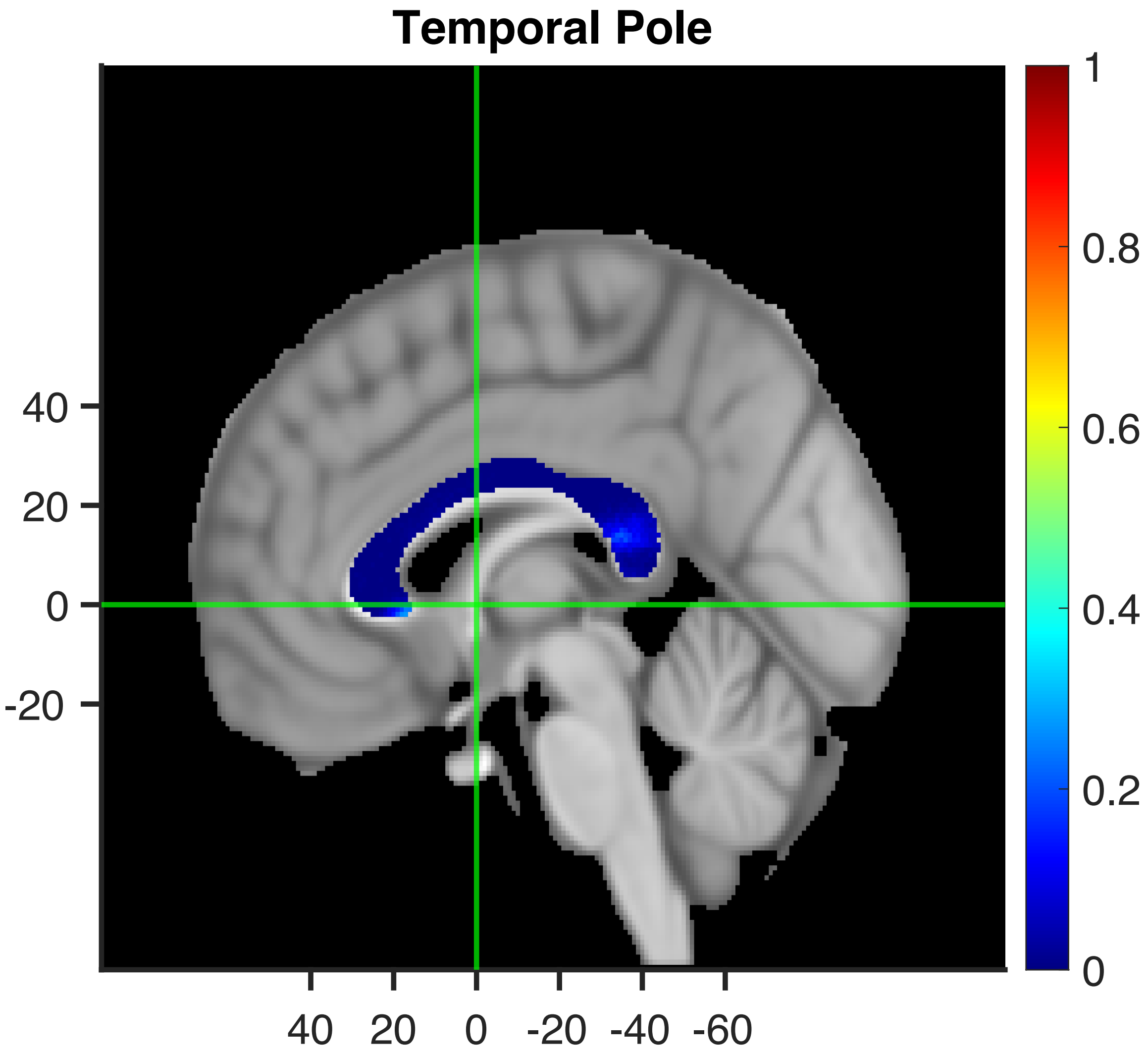
Temporal Pole
40
20
0
-20
40 20 0 -20 -40 -60
1
0.8
0.6
0.4
0.2
0

### Functional projection of cognitive functions through the human corpus callosum

Rolando Bonandrini*, Claudio Luzzatti, Marco Tettamanti

Department of Psychology, University of Milano-Bicocca and Milan Center for Neuroscience, Milan, Italy.

Supplementary Material:

- Neuropsychological validation
- Confirmatory analysis with the Schaefer et al. (2018) reference atlas

***Correspondence:**
Rolando Bonandrini

#### Neuropsychological validation

We sought to conduct a proof-of-concept analysis using lesion data. We referred to an atlas of the brain disconnections associated with neuropsychological test scores, derived from a population of stroke patients (Talozzi et al., 2023; https://neurovault.org/collections/11260/).This atlas maps, separately for 87 neuropsychological measures (across the domains of Motion, Language, Visuospatial Attention, Visuospatial Memory, Verbal Memory, and Pain) the association between structural disconnection in each voxel and performance in a neuropsychological test. The logic underlying our tentative neuropsychological validation analyses was that, among the voxels of the medial sagittal slice of the corpus callosum, those that represent the putative projection of a given cognitive function should have (relative to the voxels that are not the putative callosal projection of that function) greater association with a test measuring the same cognitive function in neuropsychological patients.

Given that the domains taken into account by Talozzi et al. (2023) do not completely overlap with those we proposed, neuropsychological validation was carried out for the callosal projections of the keywords "Motor", "Attention", "Episodic", "Language Comprehension", and "Speech Production".

More specifically, for each functional projection taken into account, we defined one or more neuropsychological tests from Talozzi et al. (2023) that tapped into the same construct (i.e., for "Motor", Action Research Arm test: Lyle, 1981; Jamar Dynamometer grip strength assessment: Fess, 1981; 9-Hole Peg test: Kellor et al., 1971; The shoulder flexion and wrist extension assessments: Dreeben, 2008; Combined walking index: Corbetta et al., 2015. http://165.232.73.88/motor_page_info/; for "Attention", Star cancellation (Behavioral Inattention Test): Wilson, Cockburn & Halligan, 1987; Mesulam Unstructured Symbol Cancellation Test: Mesulam, 1985; Posner orienting task: Posner & Cohen, 1984 http://165.232.73.88/visuospatial_page_info/, for "Episodic"; The Hopkins Verbal Learning Test; Brandt, 1991 http://165.232.73.88/verbal_memory_page_info/; for "Language Comprehension", Performing listened commands (BDAE); Comprehension of read sentence (BDAE); Comprehension of listened word (BDAE) Goodglass & Kaplan, 1983 http://165.232.73.88/language_page_info/, for "Speech Production", Semantic verbal fluency (animals): Whiteside et al., 2016; Sentence reading (BDAE) and Nonword repetition (BDAE): Goodglass & Kaplan, 1983 http://165.232.73.88/language_page_info/ ). Then, for each test, we considered the corresponding image in Talozzi et al.'s atlas relating the lesion-driven structural disconnection profile and the scores at that test in a cohort of stroke patients. We focused on the medial sagittal section, and in particular the portion included in the JHU mask of the corpus callosum. We then carried out a Wilcoxon test (non-parametric version of a two-sample *t* test) comparing the associations with the neuropsychological score in the voxels that are part of the putative callosal projection of the cognitive function of interest with those that are not. Voxels corresponding to the functional callosal projection were determined by using the same binary maps used for functional validation back-projection analyses (containing 1 in voxels whose estimated functional projection was greater than the midpoint of the range and 0 otherwise). Such binary masks and the JHU callosal ROI were down-sampled from a resolution of 1x1x1mm to 2x2x2mm to match the resolution of the atlas by Talozzi et al. (2023). In each domain corresponding to our functional projections through the callosum, if multiple Wilcoxon tests were carried out (due to multiple tests from the Talozzi et al., 2023 data set being available for that function), Bonferroni-corrected p values were used for further data interpretation. Supplementary Table S1 reports the results of the neuropsychological validation.

Within the motor domain, the neuropsychological validation was successful (with the only exception of the grip task) for tests gauging deficits in left hand movements, which is consistent with left hand apraxic symptoms following lesions to the body of the corpus callosum. For what concerns the attention domain, the neuropsychological validation showed an inconsistent pattern of results: in some instances, associations were found with performance in attention tests in the left side of the space, in others in the right side of the space, and in other with the full test with left and right halves of the space considered together. As for the episodic memory domain, validation was successful for all but two tests. In both the Speech Production and Language Comprehension domains validation was significant for all tests (with the only exception of the fluency test, for which the association barely missed formal significance).

The results of this validation procedure (Supplementary Table S1) are overall in line with those of our back-projection approach: results are clear-cut for what concerns Episodic Memory and the Motor domains (once the left bias related to callosal apraxia is considered), while inconsistent effects in the domain of Attention is coherent with the null effect we detected using the back-projection procedure. As for the language keywords (Language Comprehension and Speech Production), the neuropsychological validation yielded significant results, which are inconsistent with those we obtained through our back-projection procedure. However, we must consider that the tests taken into account involve a large set of linguistic subdomains, which makes the comparison with the neural correlates of these constructs measured through functional imaging (form which functional projections are derived) potentially hard to evaluate. In this regard, we prefer to embrace the more cautious standpoint suggested by our back-projection approach and consider validation in these two domains as unsuccessful (which is also in line with a lack of substantial evidence relating aphasic symptoms with a callosal lesion).

**Supplementary Table S1:** results of the neuropsychological validation. Domain: functional Keyword; DisconnectomeAssociation: neuropsychological test whose association with a lesion-based disconnectome map has been taken into account (see http://165.232.73.88/ for a full specification of test abbreviations). Mean_inside: average association with the test inside the ROI of the putative callosal functional projection; Mean_outside: average association with the test outside the ROI of the putative callosal functional projection ; SE_inside: standard error of the association with the test inside the ROI of the putative callosal functional projection; SE_outside: standard error of the association with the test outside the ROI of the putative callosal functional projection; p-unc: uncorrected p value of the Wilcoxon test. p-Bonf: Bonferroni-corrected p value of the Wilcoxon test; Z: Z values; ranksum: Wilcoxon rank sum statistic.

| Domain | DisconnectomeAssociation | Mean_inside | Mean_outside | SE_inside | SE_outside | punc | pBonf | Z | ranksum |
|---|---|---|---|---|---|---|---|---|---|
| Motor | laragrasp.nii | 0.050 | 0.042 | 0.001 | 0.001 | < 0.001 | < 0.001 | 4.515 | 3670 |
| Motor | laragrip.nii | 0.051 | 0.039 | 0.001 | 0.001 | < 0.001 | < 0.001 | 5.197 | 3837 |
| Motor | larapinch.nii | 0.048 | 0.040 | 0.001 | 0.001 | < 0.001 | < 0.001 | 5.165 | 3829 |
| Motor | lgrip.nii | 0.094 | 0.091 | 0.004 | 0.002 | 0.702 | 1.000 | 0.382 | 2659 |
| Motor | lpegs.nii | 0.049 | 0.038 | 0.002 | 0.001 | < 0.001 | < 0.001 | 4.511 | 3669 |
| Motor | lshflex.nii | 0.074 | 0.046 | 0.010 | 0.002 | 0.001 | 0.012 | 3.346 | 3384 |
| Motor | lwrext.nii | 0.057 | 0.046 | 0.001 | 0.001 | < 0.001 | < 0.001 | 5.827 | 3991 |
| Motor | raragrasp.nii | 0.272 | 0.326 | 0.019 | 0.017 | 0.253 | 1.000 | -1.142 | 2285 |
| Motor | raragrip.nii | 0.270 | 0.322 | 0.019 | 0.017 | 0.285 | 1.000 | -1.069 | 2303 |
| Motor | rarapinch.nii | 0.267 | 0.314 | 0.020 | 0.016 | 0.328 | 1.000 | -0.979 | 2325 |
| Motor | rgrip.nii | 0.091 | 0.068 | 0.012 | 0.003 | 0.172 | 1.000 | 1.367 | 2900 |
| Motor | rpegs.nii | 0.309 | 0.315 | 0.018 | 0.014 | 0.930 | 1.000 | -0.088 | 2543 |
| Motor | rshflex.nii | 0.278 | 0.346 | 0.020 | 0.019 | 0.227 | 1.000 | -1.208 | 2269 |
| Motor | rwrext.nii | 0.296 | 0.355 | 0.020 | 0.018 | 0.296 | 1.000 | -1.044 | 2309 |
| Motor | walk_total.nii | 0.019 | 0.010 | 0.002 | 0.001 | < 0.001 | < 0.001 | 5.091 | 3811 |
| Attention | bit_tot_miss.nii | 0.048 | 0.053 | 0.001 | 0.001 | 0.089 | 0.975 | -1.703 | 1671 |
| Attention | bit_ltot_miss.nii | 0.154 | 0.073 | 0.005 | 0.003 | < 0.001 | < 0.001 | 6.952 | 3606 |
| Attention | bit_rtot_miss.nii | 0.061 | 0.060 | 0.002 | 0.001 | 0.821 | 1.000 | -0.226 | 2001 |
| Attention | mes_l_miss.nii | 0.152 | 0.046 | 0.008 | 0.002 | < 0.001 | < 0.001 | 7.717 | 3777 |
| Attention | mes_r_miss.nii | 0.152 | 0.044 | 0.013 | 0.002 | < 0.001 | < 0.001 | 7.646 | 3761 |
| Attention | mes_tot_miss.nii | 0.145 | 0.072 | 0.008 | 0.002 | < 0.001 | < 0.001 | 7.471 | 3722 |
| Attention | pos_acc_avg.nii | 0.039 | 0.033 | 0.003 | 0.001 | 0.018 | 0.198 | 2.365 | 2581 |
| Attention | pos_acc_lv.nii | 0.040 | 0.042 | 0.002 | 0.001 | 0.268 | 1.000 | -1.108 | 1804 |
| Attention | pos_acc_rv.nii | 0.092 | 0.041 | 0.006 | 0.001 | < 0.001 | < 0.001 | 7.198 | 3661 |
| Attention | pos_acc_li.nii | 0.025 | 0.016 | 0.003 | 0.001 | 0.011 | 0.119 | 2.549 | 2622 |
| Attention | pos_acc_ri.nii | 0.040 | 0.035 | 0.002 | 0.001 | 0.024 | 0.264 | 2.258 | 2557 |
| Episodic | hvlt_delay.nii | 0.057 | 0.014 | 0.002 | 0.001 | < 0.001 | < 0.001 | 8.365 | 4612 |
| Episodic | hvlt_delayt.nii | 0.041 | 0.015 | 0.006 | 0.002 | < 0.001 | < 0.001 | 6.755 | 4218 |
| Episodic | hvlt_discrim.nii | 0.039 | 0.023 | 0.014 | 0.003 | 0.126 | 1.000 | -1.531 | 2190 |
| Episodic | hvlt_discrimt.nii | 0.064 | 0.038 | 0.005 | 0.002 | < 0.001 | < 0.001 | 5.226 | 3844 |
| Episodic | hvlt_fa1.nii | 0.094 | 0.057 | 0.010 | 0.003 | < 0.001 | 0.001 | 3.902 | 3520 |
| Episodic | hvlt_fa2.nii | 0.016 | 0.005 | 0.004 | 0.001 | < 0.001 | < 0.001 | 7.262 | 4342 |
| Episodic | hvlt_fa3.nii | 0.077 | 0.051 | 0.003 | 0.003 | < 0.001 | < 0.001 | 4.249 | 3605 |
| Episodic | hvlt_hit.nii | 0.116 | 0.035 | 0.012 | 0.005 | < 0.001 | < 0.001 | 7.000 | 4278 |
| Episodic | hvlt_learn.nii | 0.079 | 0.033 | 0.005 | 0.001 | < 0.001 | < 0.001 | 6.702 | 4205 |
| Episodic | hvlt_perc.nii | 0.037 | 0.027 | 0.004 | 0.002 | 0.017 | 0.167 | 2.393 | 3151 |
| Language Comprehension | commands_raw.nii | 0.089 | 0.043 | 0.009 | 0.002 | < 0.001 | < 0.001 | 6.019 | 3933 |
| Language Comprehension | reading_comp_raw.nii | 0.110 | 0.042 | 0.013 | 0.002 | < 0.001 | < 0.001 | 6.210 | 3979 |

| | | | | | | | | | |
|---|---|---|---|---|---|---|---|---|---|
| Language Comprehension | word_raw.nii | 0.249 | 0.087 | 0.023 | 0.005 | < 0.001 | < 0.001 | 6.819 | 4126 |
| Speech Production | animal_raw.nii | 0.086 | 0.069 | 0.006 | 0.002 | 0.020 | 0.059 | 2.332 | 2668 |
| Speech Production | reading_raw.nii | 0.133 | 0.054 | 0.018 | 0.003 | < 0.001 | < 0.001 | 4.769 | 3222 |
| Speech Production | nonword.nii | 0.152 | 0.071 | 0.022 | 0.003 | 0.004 | 0.011 | 2.900 | 2797 |

#### Confirmatory analysis with the Schaefer et al. (2018) reference atlas

We referred to the Schaefer et al. (2018) atlas, and in particular we chose the version containing 100 ROIs. The template is not symmetrical, so – by definition - it is not possible to aggregate homologous contralateral areas. As far as structural gradients are concerned (Supplementary Tables S2 and S3, Supplementary Figure S1**)**, the use of the Schaefer atlas instead of the cortical Harvard Oxford one produced a significant positive quadratic association (t=2.175, p=0.0321) between distance from the sagittal midline (x axis) and the z axis ["middle" locations pass through the callosum in its most ventral parts]. The linear association between the z coordinate of the center of mass of the ROI and the y location of maximal connectivity barely missed formal significance (p= 0.053). Otherwise, results are identical to those obtained with the cortical Harvard-Oxford Template.

**Supplementary Table S2:** associations between location of maximal connection probability along the antero-posterior (y) axis and the x, y, and z coordinates of the center of mass of each ROI.

| | **Estimate** | **Std. Error** | **t value** | **Pr(>Itl)** |
|---|---|---|---|---|
| **(Intercept)** | -17.780 | 1.011 | -17.591 | < 0.001 |
| **ROI center of mass (x) : linear** | -10.108 | 12.410 | -0.814 | 0.417 |
| **ROI center of mass (x) : quadratic** | -8.464 | 11.384 | -0.743 | 0.459 |
| **ROI center of mass (y) : linear** | 196.989 | 10.191 | 19.330 | < 0.001 |
| **ROI center of mass (y) : quadratic** | 48.566 | 13.477 | 3.604 | 0.001 |
| **ROI center of mass (z) : linear** | 21.900 | 11.155 | 1.963 | 0.053 |
| **ROI center of mass (z) : quadratic** | -0.982 | 11.940 | -0.082 | 0.935 |

**Supplementary Table S3:** associations between location of maximal connection probability along the dorso-ventral (z) axis and the x, y, and z coordinates of the center of mass of each ROI.

| | **Estimate** | **Std. Error** | **t value** | **Pr(>Itl)** |
|---|---|---|---|---|
| **(Intercept)** | 14.880 | 0.394 | 37.771 | < 0.001 |
| **ROI center of mass (x) : linear** | -1.262 | 4.837 | -0.261 | 0.795 |
| **ROI center of mass (x) : quadratic** | 9.652 | 4.437 | 2.175 | 0.032 |
| **ROI center of mass (y) : linear** | -1.356 | 3.972 | -0.341 | 0.734 |
| **ROI center of mass (y) : quadratic** | -33.784 | 5.253 | -6.432 | < 0.001 |
| **ROI center of mass (z) : linear** | 32.100 | 4.348 | 7.383 | < 0.001 |
| **ROI center of mass (z) : quadratic** | -2.944 | 4.654 | -0.633 | 0.529 |

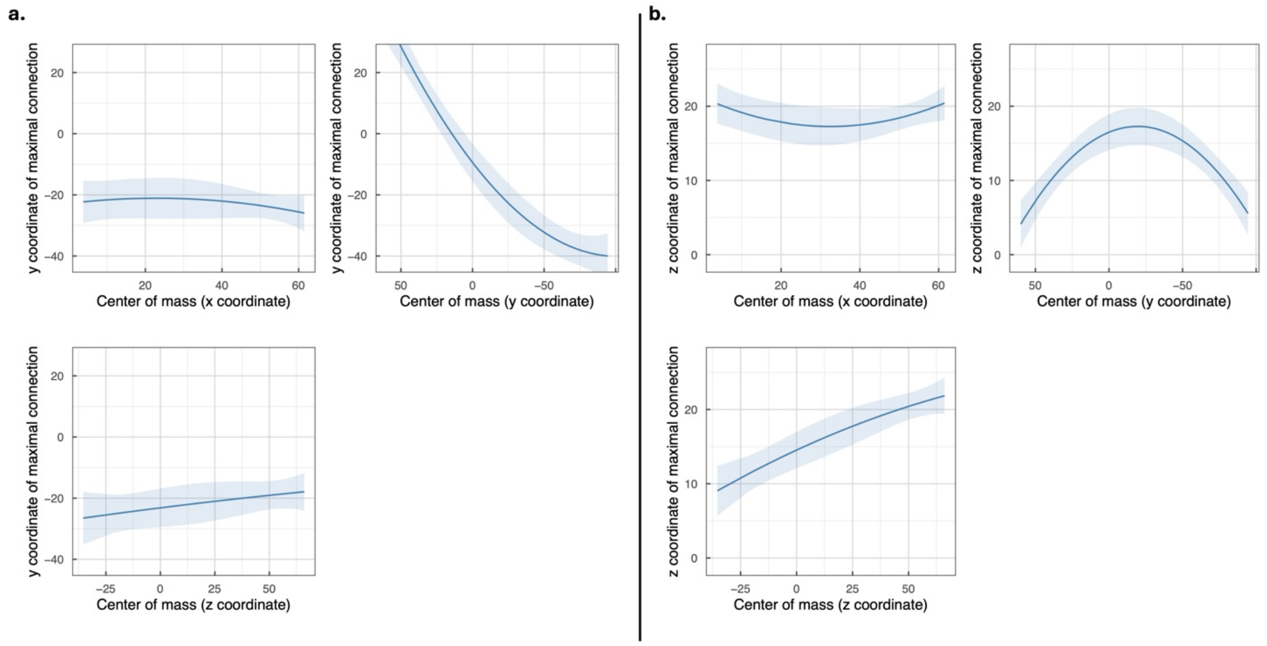

**Supplementary Figure S1.** Estimated association between the stereotactic coordinates of the center of mass (COM) of each ROI and the location of maximal connection probability along the antero-posterior axis (a) or dorso-ventral axis (b) using the Schaefer et al. (2018) atlas. Values in the lateral-medial axis (x coordinates) that are closer to zero indicate areas closer to the sagittal midline. Negative values in the antero-posterior axis (y coordinates) indicate more posterior areas. Negative values in the dorso-ventral axis (z coordinates) indicate more ventral areas.

The estimation of functional projections was carried out in the same way as for the cortical Harvard Oxford template. Crucially, the domains for which functional validation via back-projection succeeded for the Schaefer template are identical to those of the cortical Harvard Oxford atlas (Supplementary Table S4).

**Supplementary Table S4.** Results of the validation analysis (Spearman correlations) using the cortical Harvard-Oxford atlas (left) and the Schaefer et al. (2018) one (right).

| Reference atlas: Harvard Oxford | | | Reference atlas: Schaefer et al., 2018 | | |
|---|---|---|---|---|---|
| **Domain** | **Estimate** | **Pval** | **Domain** | **Estimate** | **Pval** |
| Attention | -0.284 | 0.975 | Attention | -0.104 | 0.848 |
| Auditory | 0.038 | 0.398 | Auditory | 0.056 | 0.292 |
| **Decision** | **0.381** | **0.004** | **Decision** | **0.488** | **<0.001** |
| Emotions | -0.041 | 0.609 | Emotions | 0.154 | 0.063 |
| **Episodic** | **0.310** | **0.016** | **Episodic** | **0.501** | **<0.001** |
| Language Comprehension | -0.313 | 0.985 | Language Comprehension | -0.085 | 0.801 |
| **Motor** | **0.598** | **<0.001** | **Motor** | **0.511** | **<0.001** |
| Semantic | -0.135 | 0.819 | Semantic | -0.093 | 0.821 |
| **Somatosensory** | **0.287** | **0.024** | **Somatosensory** | **0.466** | **<0.001** |
| Speech Production | 0.070 | 0.319 | Speech Production | 0.148 | 0.070 |
| **Visual** | **0.393** | **0.003** | **Visual** | **0.416** | **<0.001** |
| **Working Memory** | **0.375** | **0.004** | **Working Memory** | **0.496** | **<0.001** |

In addition, results of the functional projections showed a remarkable degree of similarity (all voxel-wise correlations [within the callosal mask] > 0.70, all p values < 0.001, see Supplementary Table S5 and Supplementary Figure S2), which is an ultimate testament of the solidity of our approach and ensuing results.

**Supplementary Table S5.** Results of the correlation analysis between the functional projections obtained usin the cortical Harvard Oxford reference atlas and the Schaefer et al (2018) one.

| | Keyword | rho | p |
|---|---|---|---|
| 1 | Emotions | 0.777 | < 0.001 |
| 2 | Visual | 0.959 | < 0.001 |
| 3 | Auditory | 0.761 | < 0.001 |
| 4 | Somatosensory | 0.925 | < 0.001 |

| | | | |
|---|---|---|---|
| 5 | Motor | 0.912 | < 0.001 |
| 6 | Episodic | 0.862 | < 0.001 |
| 7 | Working.Memory | 0.799 | < 0.001 |
| 8 | Semantic | 0.755 | < 0.001 |
| 9 | Decision | 0.881 | < 0.001 |
| 10 | Language.Comprehension | 0.774 | < 0.001 |
| 11 | Speech.Production | 0.718 | < 0.001 |
| 12 | Attention | 0.779 | < 0.001 |

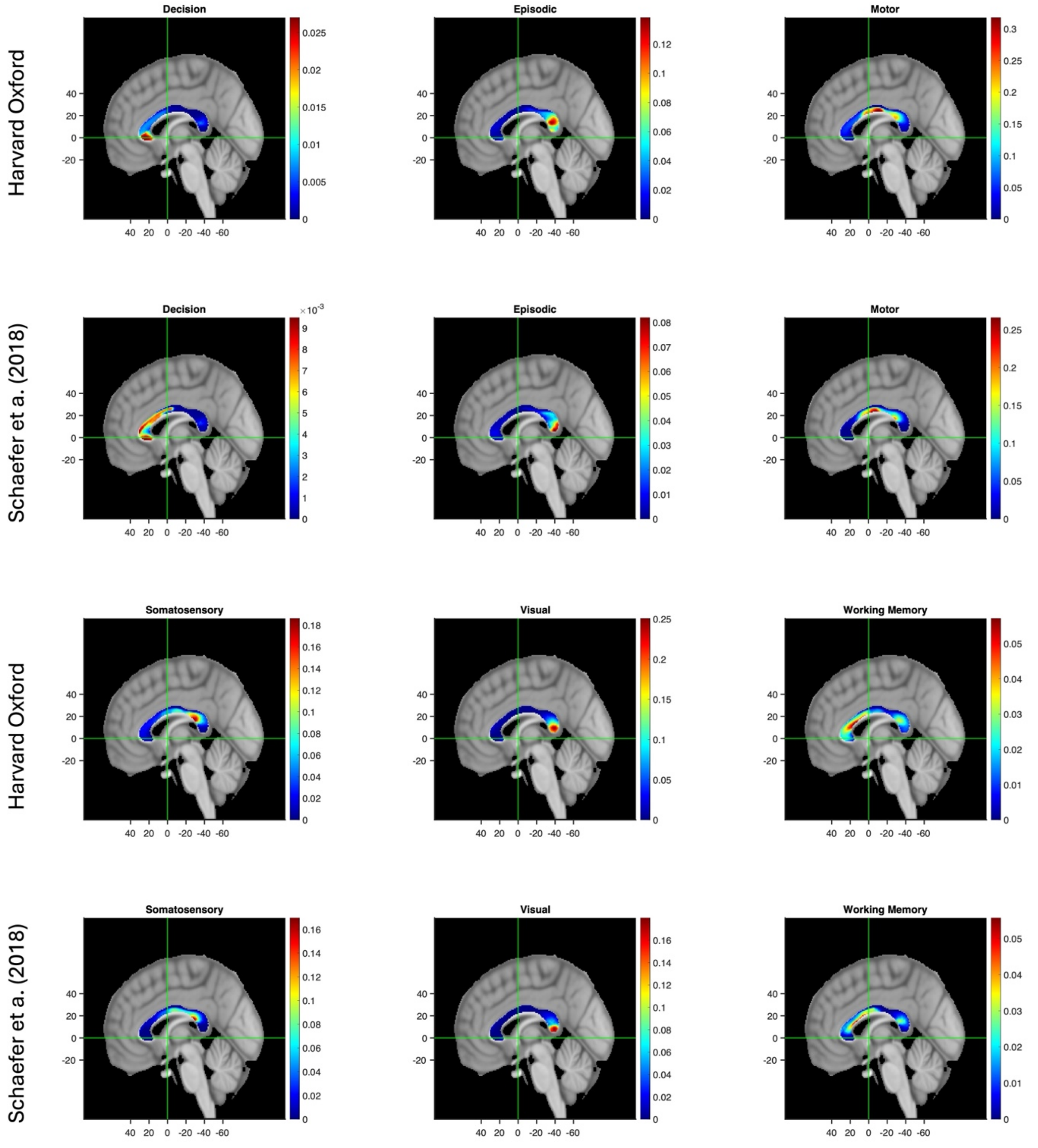

**Supplementary Figure S2.** Rendering of the functional projections obtained using the Harvard Oxford and the Schaefer et al. (2018) atlases.

As for functional gradients (Supplementary Tables S6 and S7), for y coordinates, only the linear (and not the quadratic) gradient for Working Memory turned out to be significant. In any case the direction of the trend found using the Schaefer atlas was identical to that described using the Harvard Oxford one. A similar pattern was observed for the Decision domain and z coordinates (although once again the trend was in the same direction). Working memory and z coordinates showed a positive linear trend (while with the Harvard Oxford atlas the trend was negative). However, the overall (quadratic) trend turned out to be identical across the two analyses.

**Supplementary Table S6:** Functional gradients analysis (Schaefer et al., 2018 atlas): link between the coordinate (y above, z below) in the callosum and average functional projection of the validated cognitive functions.

| | | **Estimate** | **Std. Error** | **t value** | **p value** |
|---|---|---|---|---|---|
| | (Intercept) | -5.592 | 0.070 | -80.040 | < 0.001 |
| | y coordinate : linear | -8.533 | 0.605 | -14.100 | < 0.001 |
| Visual | y coordinate : quadratic | 8.818 | 0.605 | 14.570 | < 0.001 |
| | (Intercept) | -3.801 | 0.058 | -65.424 | < 0.001 |
| | y coordinate : linear | -4.588 | 0.503 | -9.119 | < 0.001 |
| Somatosensory | y coordinate : quadratic | -6.255 | 0.503 | -12.432 | < 0.001 |
| | (Intercept) | -3.122 | 0.054 | -58.326 | < 0.001 |
| | y coordinate : linear | -2.983 | 0.464 | -6.435 | < 0.001 |
| Motor | y coordinate : quadratic | -8.604 | 0.464 | -18.560 | < 0.001 |
| | (Intercept) | -5.134 | 0.039 | -132.620 | < 0.001 |
| | y coordinate : linear | -5.379 | 0.335 | -16.050 | < 0.001 |
| Episodic Memory | y coordinate : quadratic | 7.126 | 0.335 | 21.260 | < 0.001 |
| | (Intercept) | -4.389 | 0.069 | -63.301 | < 0.001 |
| | y coordinate : linear | 2.909 | 0.600 | 4.845 | 0.000 |
| Working Memory | y coordinate : quadratic | -0.142 | 0.600 | -0.236 | 0.814 |
| | (Intercept) | -6.429 | 0.081 | -79.538 | < 0.001 |
| | y coordinate : linear | 10.205 | 0.700 | 14.578 | < 0.001 |
| Decision Making | y coordinate : quadratic | 1.735 | 0.700 | 2.478 | 0.016 |

| | | **Estimate** | **Std. Error** | **t value** | **p value** |
|---|---|---|---|---|---|
| | (Intercept) | -4.930 | 0.138 | -35.837 | < 0.001 |
| | z coordinate : linear | -6.943 | 0.778 | -8.923 | < 0.001 |
| Visual | z coordinate : quadratic | -9.578 | 0.778 | -12.309 | < 0.001 |
| | (Intercept) | -4.624 | 0.127 | -36.372 | < 0.001 |
| | z coordinate : linear | 7.104 | 0.719 | 9.880 | < 0.001 |
| Somatosensory | z coordinate : quadratic | -2.139 | 0.719 | -2.974 | < 0.001 |
| | (Intercept) | -3.967 | 0.107 | -37.006 | < 0.001 |
| | z coordinate : linear | 7.331 | 0.606 | 12.090 | < 0.001 |
| Motor | z coordinate : quadratic | -1.2937 | 0.6064 | -2.133 | 0.0415 |
| | (Intercept) | -4.613 | 0.084 | -54.835 | < 0.001 |
| | z coordinate : linear | -2.956 | 0.476 | -6.212 | < 0.001 |
| Episodic | z coordinate : quadratic | -5.061 | 0.476 | -10.635 | < 0.001 |
| | (Intercept) | -4.649 | 0.072 | -64.153 | < 0.001 |
| | z coordinate : linear | 0.902 | 0.410 | 2.199 | 0.036 |
| Working Memory | z coordinate : quadratic | -4.531 | 0.410 | -11.051 | < 0.001 |
| | (Intercept) | -5.803 | 0.069 | -84.168 | < 0.001 |
| | z coordinate : linear | -3.415 | 0.390 | -8.756 | < 0.001 |
| Decision Making | z coordinate : quadratic | -0.615 | 0.390 | -1.577 | 0.126 |

#### References

Brandt, J. (1991). The Hopkins Verbal Learning Test: Development of a new memory test with six equivalent forms. *The clinical neuropsychologist, 5*(2), 125-142.

Corbetta, M., Ramsey, L., Callejas, A., Baldassarre, A., Hacker, C. D., Siegel, J. S., ... & Shulman, G. L. (2015). Common behavioral clusters and subcortical anatomy in stroke. *Neuron, 85*(5), 927-941.

Dreeben, O. (2008). *Physical therapy clinical handbook for PTAs*. Jones & Bartlett Learning.

Fess, E. E. M. C. (1981). Clinical assessment recommendations. *American society of hand therapists*, 6-8.

Goodglass, H. and E. Kaplan, *The assessment of aphasia and related disorders (2nd ed.)*. 1983, Philadelphia: Liea & Febiger.

Kellor, M., Frost, J., Silberberg, N., Iversen, I., & Cummings, R. (1971). Hand strength and dexterity. *The American journal of occupational therapy: official publication of the American Occupational Therapy Association, 25*(2), 77-83.

Lyle, R. C. (1981). A performance test for assessment of upper limb function in physical rehabilitation treatment and research. *International journal of rehabilitation research, 4*(4), 483-492.

Mesulam, M.M., Principles of Behavioural Neurology: Tests of Directed Attention and Memory (Contemporary Neurology Series). 1985: F.A. Davis Company.

Posner, M.I. and Y. Cohen, Components of visual orienting. *Attention and performance X: Control of language processes*, 1984. 32: p. 531-556.

Schaefer, A., Kong, R., Gordon, E. M., Laumann, T. O., Zuo, X. N., Holmes, A. J., ... & Yeo, B. T. (2018). Local-global parcellation of the human cerebral cortex from intrinsic functional connectivity MRI. *Cerebral cortex, 28*(9), 3095-3114.

Talozzi, L., Forkel, S. J., Pacella, V., Nozais, V., Allart, E., Piscicelli, C., ... & Thiebaut de Schotten, M. (2023). Latent disconnectome prediction of long-term cognitive-behavioural symptoms in stroke. *Brain, 146*(5), 1963-1978.

Whiteside, D. M., Kealey, T., Semla, M., Luu, H., Rice, L., Basso, M. R., & Roper, B. (2016). Verbal fluency: language or executive function measure?. *Applied Neuropsychology: Adult, 23*(1), 29-34.

Wilson, B., Cockburn, J., & Halligan, P. (1987). Development of a behavioral test of visuospatial neglect. *Archives of physical medicine and rehabilitation, 68*(2), 98-102.
